# Megabarcoding dark taxa – Assessing the utility of mass DNA barcoding for phorid fly species discovery

**DOI:** 10.1101/2025.07.21.665927

**Authors:** Jiri Vihavainen, Niina Kiljunen, Pekka Pohjola, Jaclyn McKeown, Eveliina Oinonen, Marko Mutanen, Jaakko Pohjoismäki

## Abstract

Many hyperdiverse, small-bodied insect families contain numerous undescribed species, generally termed “dark taxa”. Scuttle flies (Diptera: Phoridae), being among the most diverse insect groups globally, are a prime example. DNA barcoding can help delineating dark taxa, particularly when integrated with morphology, and/or additional molecular evidence.

We sequenced *COI*-barcodes from 9,120 Finnish phorid specimens and initially identified them using BOLD database. Furthermore, species identifications of all 843 non-*Megaselia* specimens were confirmed morphologically. Initially, the BOLD-based identifications matched the morphological identifications only in 68% of the cases, which resulted from many misidentifications in BOLD. After adjusting the BOLD reference identifications based on morphological analyses of male features, we established a reliable framework for female identification. This is advantageous for future identification of females, as they are often excluded from traditional identification keys.

Only two species was discovered as new to Finland, demonstrating that Finnish non-Megaselia fauna is well-known. Although DNA barcodes show great promise for identifying phorids, incorrectly identified reference sequences remain challenging, not the functionality of *COI* itself. The number of *Megaselia* BINs greatly exceeded the known Finnish species count, with many sequences lacking matches in BOLD. This further highlights *Megaselia* as a particularly dark group, for which genetic tools are essential for uncovering species identities and assessing diversity.

## Introduction

Despite over 260 years of effort in taxonomy, our grasp of the planet’s biodiversity is still just scratching the surface (Godfray 2002). Alongside the ongoing anthropogenic biodiversity crisis, the taxonomists are facing a taxonomic impediment, where inefficiency and lack of resources have slowed down the progress in species delimitation and taxonomic descriptions (Engel et al. 2021). Meanwhile, the species extinction rate continues to exceed the emergence of new species, and the number of taxonomist experts continues to dwindle (Hebert et al. 2003, Engel et al. 2021). It is estimated that the world would need approximately 15,000 taxonomists to identify all life on Earth if based on traditional largely morphological practices. The current number of taxonomists is not nearly enough to document the biodiversity, and their expertise are biased towards better known groups of organisms. Example of this limitation is a nation-wide inventory of insects in Sweden, where after enormous effort of two decades, only 1% of the samples were identified to species level (Karlsson et al. 2020, van Klink et al. 2024). Molecular methods, such as DNA barcoding, could offer an efficient solution to overcoming the taxonomic impediment (Hebert et al. 2003). These methods have been successfully implemented for several groups previously considered challenging or even inaccessible with traditional morphological means (Kekkonen & Hebert 2014, Kekkonen et al. 2015, Meierotto et al. 2019, Sharkey et al. 2021a, Sharkey et al. 2021b, Hartop et al. 2022, Sharkey et al. 2023). While national and regional DNA barcoding initiatives can produce huge quantities of information, an ever-growing number of sequences cannot be assigned to species level, especially those belonging to hyperdiverse, species-rich groups of small-bodied insects (Page 2016, Hausmann et al. 2020, Chimeno et al. 2022, Srivathsan et al. 2023). Groups with most species remaining undescribed have been termed “dark taxa”, often representing taxonomically speciose groups, such as Diptera and Hymenoptera (Hartop et al. 2022). While dark taxa may constitute the majority of all biodiversity in the tropics, they can also represent a significant portion of certain hymenopteran and dipteran families in Europe, despite the long history of taxonomic research in the subcontinent. One such recognized dark taxon are the scuttle flies (Diptera: Phoridae).

The scuttle flies have been proposed to be among the most diverse groups of insects with 4,400 described species world-wide (Phorid Catalog 2024), while 200,000 estimated species are just in Afrotropical region alone (Srivathsan et al. 2019). Also, the larval feeding habits are exceptionally diverse (Disney 1994, Courtney et al. 2017). While many scuttle flies are important scavengers and decomposers of decaying organic matter, there are also many herbivorous, predatory and parasitoid species. For example, the genus *Pseudacteon* Coquillett, 1907 is infamous for parasitizing fire ants by decapitating them during their larval stage and using their empty head capsule for pupation site (Porter 1998). Because of this feature, they have been used as a biological control agent in North America (Maschwitz et al. 2008), although the introduction has not had major successes in controlling the fire ant populations (Morrison & Porter 2005, Callcott et al. 2011). The most diverse genus of scuttle flies is *Megaselia* Rondani, 1856, with over 1,700 described species (Phorid Catalog 2024), with the majority of species remaining undescribed (Hartop et al. 2024). In fact, the *Megaselia* are considered as an extraordinarily diverse genus, even among other species rich insect taxa (Hartop et al. 2022).

Identifying specimens into species using DNA barcodes has been highly successful across a variety of invertebrate groups, such as true bugs (Park et al. 2011), chironomid midges (Brodin et al. 2012), neuropterids (Morinière et al. 2014), bees (Schmidt et al. 2015), beetles (Pentinsaari et al. 2014), tachinid flies (Pohjoismäki et al. 2016), moths (Hausmann et al. 2013) and butterflies (Dincă et al. 2021). Intermediate results have been obtained for families considered to be taxonomically inaccessible, such as highly diverse flies and midges (Morinière et al. 2019). Several national barcode initiatives, for instance in Canada (deWaard et al. 2019), Germany: German Barcode of Life Initiative (GBOL III: Dark Taxa), and in Finland: Finnish Barcode of Life (Roslin et al. 2022) aim to generate DNA barcode reference libraries in respective countries, and some regional consortia (e.g., iBOL Europe, https://iboleurope.org) under the International Barcode of Life (https://ibol.org) research alliance work for the same goal over wider geographic regions.

Although DNA barcoding shows huge promise for completing the inventory of life (Hebert et al. 2003), it has some shortcomings too (Rubinoff et al. 2006, Virgilio et al. 2010, Collins & Cruickshank 2013). Also, because DNA barcoding relies on reference sequences for identification, a sound and comprehensive reference library is required (Hebert et al. 2003, Virgilio et al. 2010). DNA barcoding was initially poorly received by traditional taxonomist societies as resulting a loss of intellectual content (Lipscomb et al. 2003), but Gregory (2005) quickly noted that the DNA barcoding will not replace morphological approach of quantifying life but will be aiding it instead.

Optimally, comprehensive DNA barcode libraries (e.g., Wirta et al. 2016, Roslin et al. 2022) should enable the identification of unknown specimens into species level. This is not only required to facilitate the identification of individual specimens, but also for the purpose of species inventories conducted through metabarcoding (Taberlet et al. 2012). In the case of specimens without a match in the database, operational taxonomic units (OTUs) such as that based on Barcode Index Numbers system (BINs), can be used as units of putative species (Ratnasingham & Hebert 2013). Besides the aforementioned issue with taxonomic impediment preventing the verification of the identity by morphological means, the utility of DNA barcoding for species identification is highly dependent on the reliability of the available databases. The problem is amplified if reference databases contain frequent misidentifications that result in subsequent specimens becoming misidentified and established in other databases through a circular identification process.

In the present work, we sought to elucidate the usability of DNA barcoding to identify Finnish scuttle flies (Diptera: Phoridae). To achieve this, we sequenced the full-length *COI* DNA barcodes for 9,120 scuttle fly specimens trapped throughout Finland and utilized the BOLD identification engine to obtain tentative identities for species already present in the database. Our focus was in non-Megaselian scuttle flies, as they are possible to morphologically identify to species level without years of thorough identification training and, therefore, can be used to confirm tentative species identities. Besides testing the *COI* ability to distinguish scuttle fly species, our combined approach also allowed us to curate and correct non-Megaselian species identifications in the BOLD database.

## Materials and Methods

Insect material was collected using Malaise, or window trap approaches with 80% to absolute ethanol or 70% glycol as collection liquid, respectively. The traps collect mainly flying insects and therefore are unlikely to catch wingless females of some phorid genera in high abundance. The sampling was done across Finland in 14 localities (Figure 1A) from May to October of 2015-2017 and 2020-2022. A more detailed information about sampling is listed in the S-Table 1. The scuttle flies (Diptera: Phoridae) were sorted from the insect material. As scuttle flies were extremely abundant in this material, a randomized subset of 9,120 was taken by placing all sorted scuttle flies from each site and collecting period to separate petri dishes from which individual specimens were picked at random and placed in 96 microplates with 30 microliter of 96% ethanol. The microplates were sent to Canadian Centre for DNA Barcoding (CCDB) Guelph, Canada, where the samples were photographed, and DNA barcoded for the full-length *COI* gene marker (658 bp) using the LepF1 and LepR1 primers (Hebert et al. 2004). The barcoding was done using the CCDB pipeline for Sequel II sequencing (Hebert et al. 2018).

**Figure 1.**
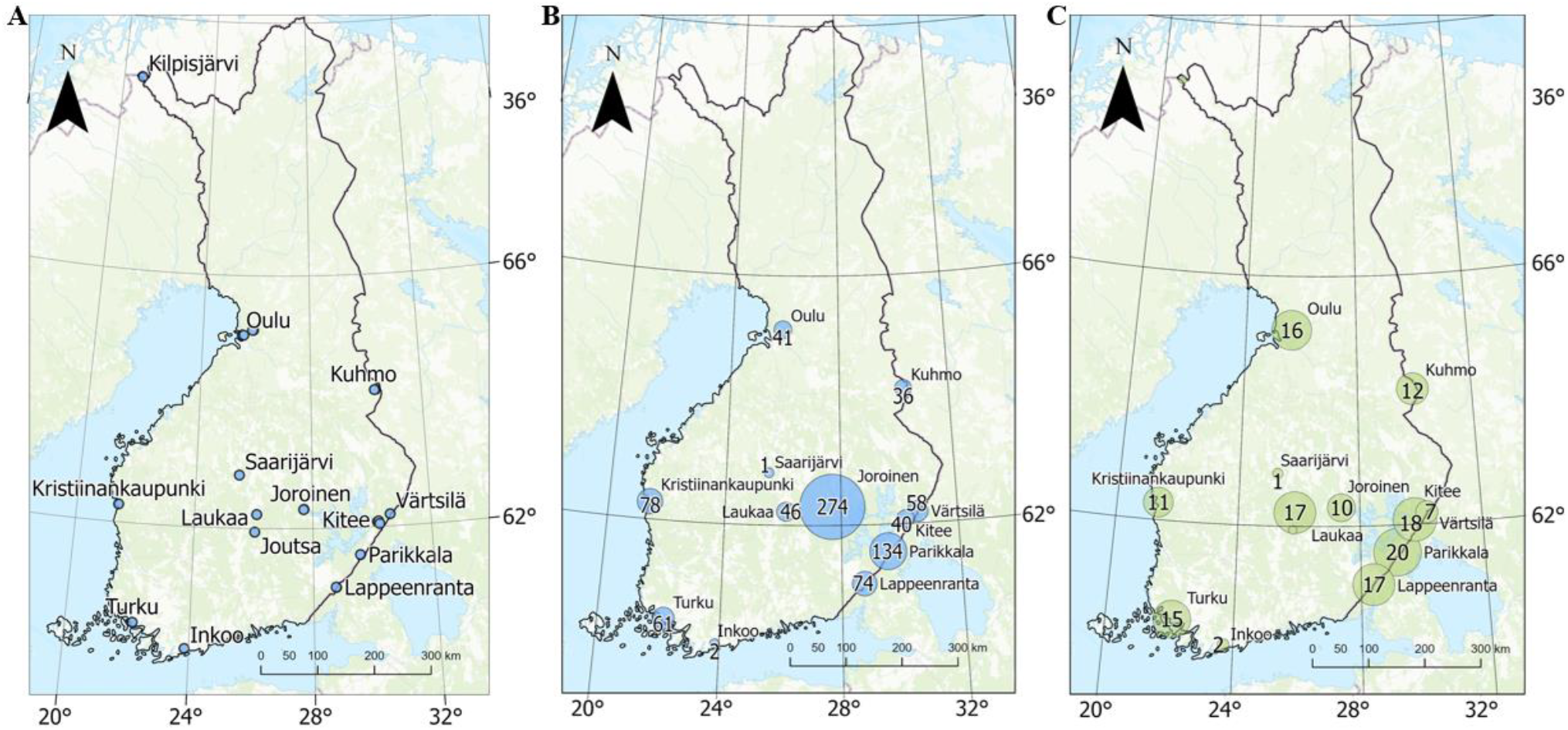
(A) Sampling sites across Finland. Sampling was done using 27 traps in 14 localities. (B) Captured and processed non-*Megaselia* specimens in each locality represented by blue circles according to their abundancy. (C) Count of captured and processed non-*Megaselia* BIN diversity of each locality are represented by green circles depending on their abundancy.

The DNA-based identification of specimens was first done using BOLD Identification Engine (Ratnasingham & Hebert 2007) with 98% K2P sequence similarity used as a proxy for preliminary species level assignment. Each BIN identified only to family level was further identified to genus level using the BOLD Identification Engine with 90% K2P sequence similarity, and the genus assignments were verified using morphology. Subsequently, morphological identification of all non-*Megaselia* species was performed using relevant identification keys (Schmitz et al. 1981, Disney 1983, Gotô 1986, Disney & Withers 2009, Disney 2013, Liu & Yang 2016), where applicable. Due to the lack of identification keys for females, particularly within the genus Phora Latreille, 1796, morphological analyses of females in this genus were limited to determine their sex. Additionally, 29 specimens across several genera were too damaged in order to reliably identify their sex. Female specimens of other genera were identified morphologically, alongside their male counterparts. For *Phora*, females were identified by matching their barcodes to those of morphologically verified males. Barcode-based inference was also applied to females of other genera and degraded specimens to validate the method and ensure accurate species identification. A total of 843 specimens were identified using this combined approach of morphological examination and barcode inference. Among these, one specimen produced a DNA barcode sequence matching *Metopina oligoneura* (Mik, 1867) but was unequivocally identified as *Phalacrotophora berolinensis* Schmitz, 1920 based on morphology, as the two genera are distinctly different. Consequently, this specimen was excluded from further analyses. The source of the contamination remains unclear, as the morphological examination was performed on the sequenced specimen.

## Results

Out of 9,120 analysed specimens, 8,363 yielded a high-quality DNA barcode (>600 bp without stop codons) for further analyses. Validated sequences were assigned into 424 BINS, where the most represented genus was *Megaselia* (Figure 2A). BOLD Identification Engine with 98% similarity assigned 6,014 specimens (72%) to species level, belonging to 171 species (S-Table 2) in total. Of these, 43 were non-*Megaselia* species, represented by 798 specimens. Remaining family level unique BINs were further identified to a genus level with 90% similarity, increasing the number of non-*Megaselia* specimens to 843 belonging to 55 BINs in total. Morphological identification of these specimens verified 15 genera and 51 species, excluding the one aforementioned contaminant. These contained 18 species represented by only one specimen (singletons), while the most numerous species (*Phora pubipes* Schmitz, 1920) represented 15% of the specimens in the samples. In some cases, two BINs were found in a single morphological species (Figure 3A & B). This phenomenon occurred with *Anevrina thoracica* (Meigen, 1804), *Phora obscura* (Zetterstedt, 1848), *P. artifrons* Latreille, 1796, and *Triphleba nudipalpis* (Becker, 1901). Meanwhile, a shared BIN was observed between *Diplonevra florea* and *D. glabra* (Figure 3C). Initial barcode-based identification produced 68% success rate for correct species assignment, before corrections based on morphology were uploaded to the BOLD database. The morphological analyses improved the accuracy of BOLD reference database, increasing the identification success to 100%.

**Figure 2.**
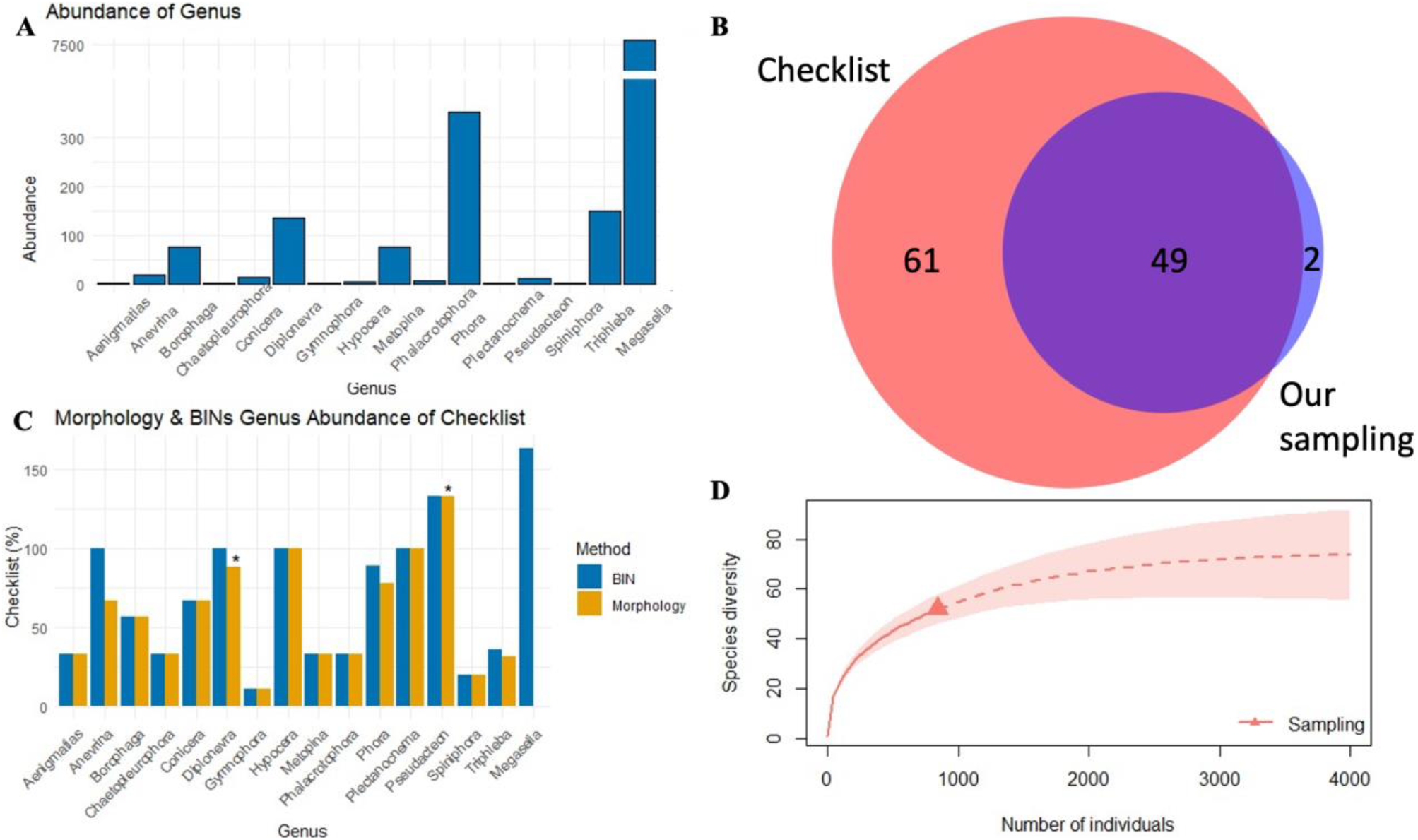
(A) Genus-level representation of the specimens from our sampling. Note how *Megaselia* specimens significantly outnumber other genera. (B) Venn-diagram with known Finnish non- *Megaselia* fauna, and species found in our sampling. The overlap area represents 46% of the known Finnish species. (C) A diagram showing how well each genus was represented in the Malaise trap samples, compared to the species list of Phoridae in FinBIF database (%). Species identified using morphology or BINs, are shown separately. Note that *Megaselia* specimens were not morphologically examined. Some morphologically identified species might have more than one BIN. * Genera with new species to Finland. (D) Rarefaction plot to estimate the species richness of non-*Megaselia* species of Finland.

**Figure 3.**
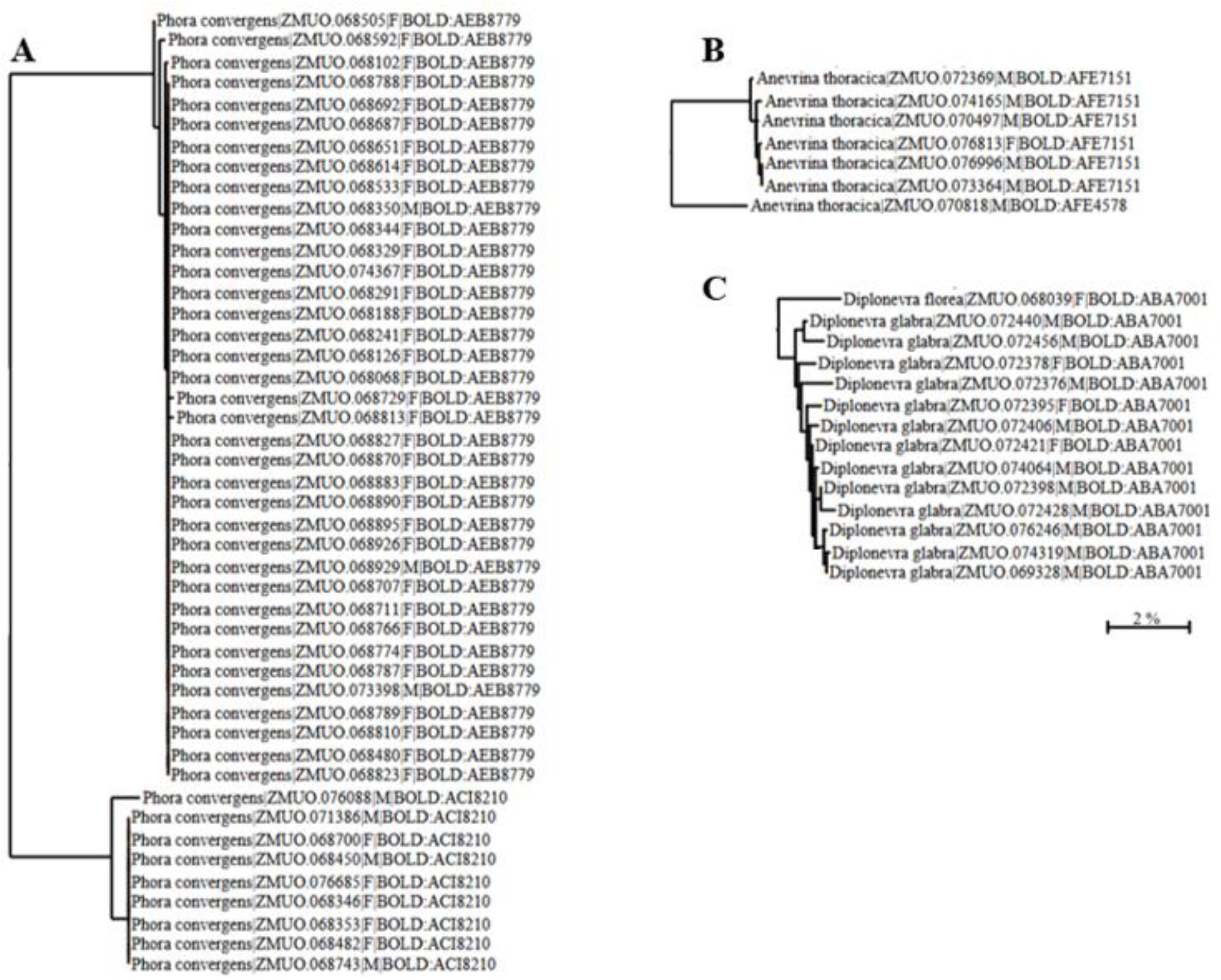
Showcase of NJ tree with examples of species sharing multiple BINs (A & B), and two species sharing the same BIN (C). (A) *Phora convergens* Schmitz, 1920 is split into two BINs, but no geographical pattern was detected. (B) *Anevrina thoracica* is split into two BINs. (C) *Diplonevra florea* and *D. glabra* share the same BIN, although morphological analyses revealed them to represent two separate morphological species. Scale bar: 2% difference.

Cross-referencing the determined species against known Finnish fauna (Kahanpää 2014, FinBIF 2024) revealed our sampling to cover 70% of known Finnish genera (Table 1) and 46% of the non- *Megaselia* species (Figure 2B, Made with DeepVenn (Hulsen 2022)). The morphological analyses verified two new species for Finland: *Diplonevra florea* (Fabricius, 1794), and *Pseudacteon brevicauda* Schmitz, 1925. Each genus was compared against known checklist and database of Finnish species as bar plot (Figure 2C). In the comparison, the BIN count is slightly higher due to some species containing multiple BINs.

**Table 1.**
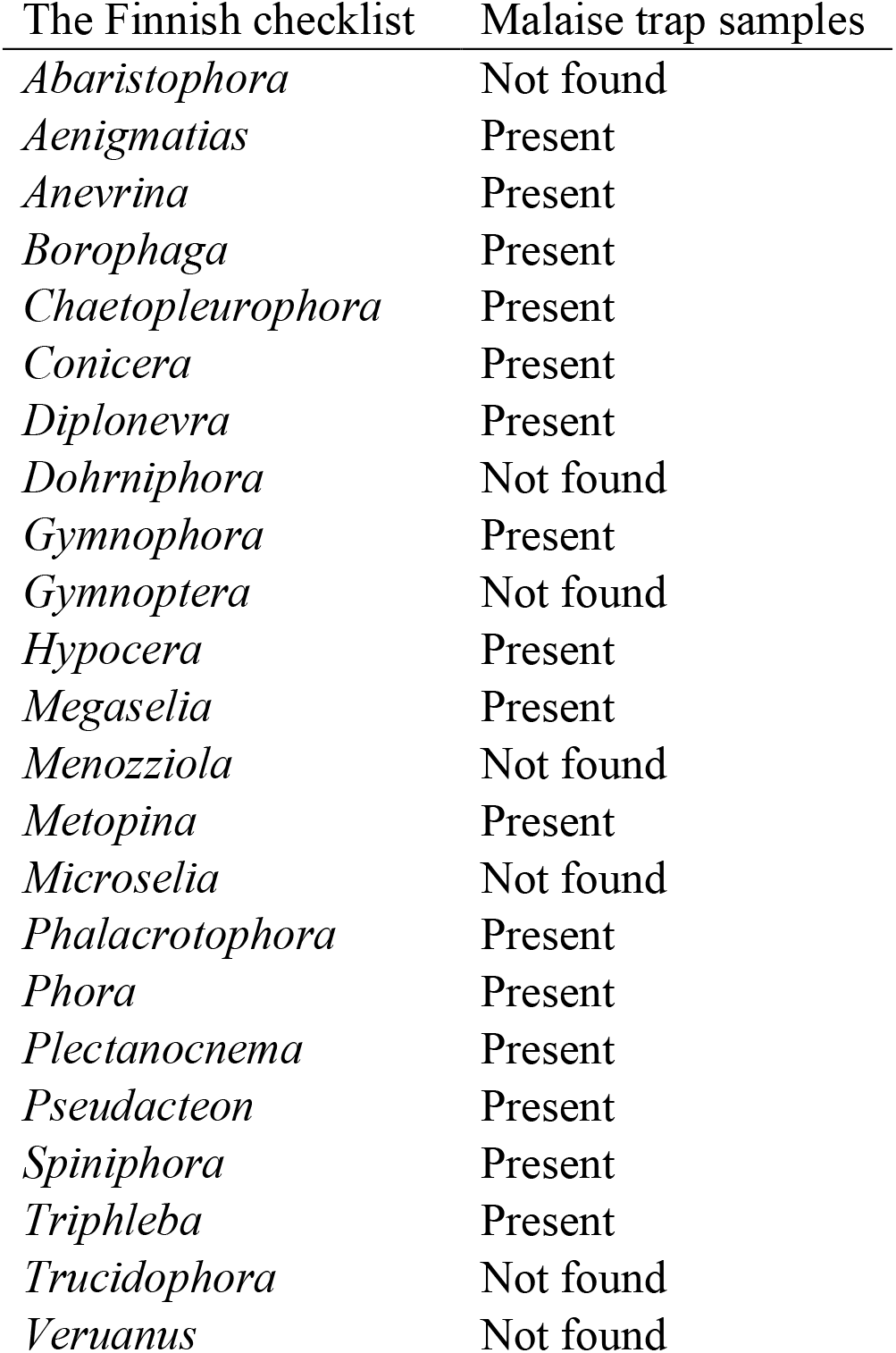
Genus representation among the samples.

| The Finnish checklist | Malaise trap samples |
| --- | --- |
| *Abaristophora* | Not found |
| *Aenigmatias* | Present |
| *Anevrina* | Present |
| *Borophaga* | Present |
| *Chaetopleurophora* | Present |
| *Conicera* | Present |
| *Diplonevra* | Present |
| *Dohrniphora* | Not found |
| *Gymnophora* | Present |
| *Gymnoptera* | Not found |
| *Hypocera* | Present |
| *Megaselia* | Present |
| *Menozziola* | Not found |
| *Metopina* | Present |
| *Microselia* | Not found |
| *Phalacrotophora* | Present |
| *Phora* | Present |
| *Plectanocnema* | Present |
| *Pseudacteon* | Present |
| *Spiniphora* | Present |
| *Triphleba* | Present |
| *Trucidophora* | Not found |
| *Veruanus* | Not found |

The DNA barcodes effectively distinguish the non-*Megaselia* species. The mean minimum of K2P divergence to the nearest neighbour is 9.35% (range 1.4 – 20.9%, SD = 3.46, SE = 0.07), which is higher than reported for Tachinidae (5.51% with 366 species, Pohjoismäki et al. 2016), but lower than in Coleoptera (11.99% with 1872 species, Pentinsaari et al. 2014), but it should be noted that the latter includes multiple families. Similarly, the relatively large K2P divergence in our non- *Megaselia* sample can be attributed to differences between genera, as divergences within species- rich genera with recent radiations, such as Tachinidae (Stireman et al. 2021), tend be smaller. For example, less than 2% divergence was observed between *Diplonevra glabra* Schmitz, 1927 and *D. florea*.

A rarefaction plot (Figure 2D) was generated to estimate the efficiency of our sampling effort. The curve shows that while our sampling effort rapidly covered half of the species, more remain to be discovered, as evidenced also by the number of singleton observations. Our sampling covered 51 non-*Megaselia* species (Figure 2B), and the extrapolations from the rarefaction plot would estimate the Finnish species count for this group to slightly exceed 100 species (Figure 2D). Considering that 110 non-*Megaselia* species were previously known from Finland (Figure 2B), this further highlights the comprehensive current knowledge of the country’s fauna.

## Discussion

To elucidate the utility of DNA barcoding for species inventories of hyperdiverse taxa, we sequenced a geographically representative sample of 9,120 phorid fly specimens from Finland. The barcoding was successful for 8,363 specimens, with samples collected in ethanol yielding the best results. 7,520 of the specimens belong to genus *Megaselia* and 843 to 15 other genera of Phoridae, representing 70% of all genera reported from Finland (Table 1) (S-Table 3). The species identifications for the 843 non-*Megaselia* specimens, representing 55 BINs, were verified by morphologically determining the males (all genera) and females (excluding genus *Phora*), and then inferring the species identities of *Phora* females based on matching male barcodes. This method was also applied to all females to validate its accuracy. Using this approach, a total of 51 non- *Megaselia* scuttle fly species were identified. Since the barcodes were able to unambiguously distinguish the males of different species, we argue that the same approach can be reliably used to identify females. This is particularly important in the contexts such as biomonitoring, where females of many genera are usually overlooked due to identification difficulties. Additionally, this approach provides new opportunities to examine and identify morphological differences between females and aiding in developing new morphological identification keys. Interestingly, *Phora convergens* is divided into two BINs, which may represent two distinct but morphologically cryptic species (Figure 3A). Apart from *P. convergens*, there is only one known species with a similar right epandrial appendage, namely *P. indivisa* known from the Alps and described based on a single specimen (Schmitz et al. 1981, Kahanpää personal communication 2024). However, the specimens do not morphologically match its species description. Sequencing of nuclear markers would probably provide the most effective way to determine if the name *P. convergens* currently contains two biologically distinct species. Other species with multiple BINs include *Anevrina thoracica* (Figure 3B), *Phora obscura, P. artifrons*, and *Triphleba nudipalpis*. Conversely, a shared BIN was observed between *Diplonevra florea* and *D. glabra*, although our results show that the two can also be distinguished by the clustering of their barcodes on NJ-tree (Figure 3C).

Despite including only 14 localities, our sampling reached 46% of the known Finnish non- *Megaselia* fauna. Only two species (*Diplonevra florea* and *Pseudacteon brevicauda*) was new to Finland. This makes Finnish non-*Megaselia* fauna well known, continuing a trend seen in rest of Europe where new non-*Megaselia* species are found scarcely (Hartop et al. 2022, Grundmann & Kappert 2023). Also, in our sampling, the collection was focused mostly to Central and Eastern parts of the country, with only few sites on the coastline (Figure 1B & 1C). With our current sampling, the rarefaction curve did not reach asymptote (Figure 2D), which is likely because our sampling failed to capture several known genera. Then again, the sampling discovered two new species to Finland, indicating the importance of adequate sampling effort needed with this taxon. It is evident from singleton observations that many species are either rare or inhabit a different habitat to those we sampled. Evidently, the genus *Megaselia* is where the majority of new species of Phoridae remain to be discovered. For example, the current FinBIF database contains 226 *Megaselia* species, whereas the BIN count of 369 suggest the true species count being at least a third higher or even more. Previous studies have revealed a similar trend. For example, in the neighbouring country of Sweden (Hartop et al. 2024) 500 new scuttle fly species for the country were found, while in Germany an increase of 12% of species was reported (Caruso et al. 2024). Given the high performance of DNA barcoding in differentiating species of non-*Megaselia*, they could establish a robust framework for species discovery and further taxonomic research also for the *Megaselia*, although this requires verification. This genus is notorious for many taxonomic difficulties (Chimeno et al. 2022, Hartop et al. 2022, Hartop et al. 2024, Caruso et al. 2024), rendering species identification and discovery extremely laborious. One solution could be to complement DNA barcodes with nuclear loci to further validate species identities.

In Sweden, Hartop et al. (2024), estimated the scuttle fly species count to be around 652–713. The true count is difficult to predict, as scuttle flies’ ecology is very diverse, and it seems that rarer species also inhabit specific environments or microhabitats, while the most common species can be found with ease. This can be said from our sampling as well, as we captured 18 singleton species, of which many inhabited different areas. The most abundant species in our material was *Phora pubipes* with 130 specimens, followed by *P. tincta* Schmitz, 1920 with 117 specimens, the two altogether covering 29% of all the non-*Megaselia* specimens. Hartop et al. (2024), alongside others (Disney et al. 1982, Brown 1996, Borkent et al. 2018), concluded that a single sampling approach hardly can capture all taxa as discovery of many species will require a specific trapping method. Therefore, comprehensive sampling would require enormous effort and several trap types to provide an accurate estimate of species count. In this regard, new species will most likely be discovered in future monitoring surveys and by lucky accidents.

Initially, the barcode success rate to identify specimens correctly was as low as 68%, which however turned out to be due to many misidentifications in the reference database rather than poor performance of DNA barcoding. After re-examining the problematic reference specimens through morphological analysis, the success rate increased to 100%, indicating that the main source of error was erroneous identifications of reference specimens. The significance of misidentifications and other ‘operational factors’ was examined and discussed by Mutanen et al. (2016). As a conclusion, the DNA barcodes perform well in identifying species of Phoridae. To this end, it is evident that a comprehensive and validated reference database is a necessity to bring Phoridae and other massively diverse dark taxa among efficient biomonitoring.

## Acknowledgements

The authors would like to thank the Centre for Biodiversity Genomics, Guelph, Canada, for their input to this paper. Special thanks go to Amy Thompson and her team at CBG, Canada, for their invaluable support. We would also like to express our appreciation to following people who assisted with field work: Mikko Vallinmäki, Laura Havukainen, Sylvia Mutanen, Riikka Jarkko, Lari Heikkinen, Oula Kalttopää, Mervi Laaksonen & Sampsa Malmberg. The authors also acknowledge the UEF laboratory personnel for their contributed effort: Vili Jormanainen, Juho Kolehmainen and Jussi Hämäläinen. Authors thank Mr. Jere Kahanpää (Finnish Museum of Natural History, Helsinki, Finland) for his invaluable discussions on potential new species discoveries in the Finnish fauna, as well as aid with morphological analyses. We appreciate Mr. Antti Haarto (Zoological Museum, Turku, Finland) for kindly providing much of the identification literature on Phoridae and, together with Mr. Kaj Winqvist (Turku, Finland), for sharing information on Finnish species.

## Author contributions

JP and MM conceived the study. JV, NK, PP carried out the research. JP and MM designed the methodology. NK carried out the field work. JV carried out molecular and morphological analyses. JM and EO provided visualization resources. JV and JP analysed the data. MM and JP acquired financial resources. JV wrote the first draft with contributions from JP, MM, NK. All authors reviewed the manuscript and have provided critical suggestions/additions to the text.

## Data availability

The complete dataset supporting this study is available on The Barcode of Life Data Systems (https://boldsystems.org), dataset DS-0798.

## References

Borkent A, Brown BV, Adler PH, de Souza Amorim D, Barber K, Bickel D, Boucher S, Brooks SE, Burger J, Burington ZL, Capellari RS, Costa DNR, Cumming JM, Curler G, Dick CW, Epler JH, Fisher E, Gaimari SD, Gelhaus J, Grimaldi DA, Hash J, Hauser M, Hippa H, Ibáñez-Bernal S, Jaschhof M, Kameneva EP, Kerr PH, Korneyev V, Korytkowski CA, Kung G-A, Kvifte GM, Lonsdale O, Marshall SA, Mathis WN, Michelsen V, Naglis S, Norrbom AL, Paiero S, Pape T, Pereira-Colavite A, Pollet M, Rochefort S, Rung A, Runyon JB, Savage J, Silva VC, Sinclair BJ, Skevington JH, Stireman III JO, Swann J, Vilkamaa P, Wheeler T, Whitworth T, Wong M, Wood MD, Woodley N, Yau T, Zavortink TJ, Zumbado MA. 2018. Remarkable fly (Diptera) diversity in a patch of Costa Rican cloud forest: Why inventory is a vital science. Zootaxa 4402(1): 053–090. Doi: 10.11646/zootaxa.4402.1.3.

Brodin Y, Ejdung G, Strandberg J, Lyrholm T. 2012. Improving environmental and biodiversity monitoring in the Baltic Sea using DNA barcoding of Chironomidae (Diptera). Molecular Ecology Resources 13: 996–1004. Doi: 10.1111/1755-0998.12053.

Brown BV. 1996. Phorid newsletter 5. https://phorid.net/newsletters/pnwes5.pdf.

Callcott A-M, Porter SD, Weeks Jr. RD, Graham LC, Johnson SJ, Gilbert LE. 2011. Fire ant decapitating fly cooperative release programs (1994-2008): Two Pseudacteon species, P. tricuspis and P. curvatus, rapidly expand across imported fire ant populations in the southeastern United States. Journal of Insect Science 11:19. Doi: 10.1673/031.011.0119.

Caruso V, Hartop E, Chimeno C, Noori S, Srivathsan A, Haas M, Lee L, Meier R, Whitmore D. 2024. An integrative framework for dark taxa biodiversity assessment at scale: A case study using Megaselia (Diptera, Phoridae). Insect Conservation and Diversity 17: 968–987. Doi: 10.1111/icad.12762.

Chimeno C, Hausmann A, Schmidt S, Raupach MJ, Doczkal D, Baranov V, Hübner J, Höcherl A, Albrecht R, Jaschhof M, Haszprunar G, Hebert PDN. 2022. Peering into the Darkness: DNA Barcoding Reveals Surprisingly High Diversity of Unknown Species of Diptera (Insecta) in Germany. Insects 13: 82. Doi: 10.3390/insects13010082.

Collins RA & Cruickshank RH. 2013. The seven deadly sins of DNA barcoding. Molecular Ecology Resources 13: 969–975. Doi: 10.1111/1755-0998.12046.

Courtney G, Pape T, Skevington J, Sinclair B. 2017. Biodiversity of Diptera. 867 p. Insect Biodiversity: Science and Society, Volume I, 2^nd^ edition. John Wiley & Sons Limited. Doi: 10.1002/9781118945568.ch9.

deWaard JR, Ratnasingham S, Zakharov EV, Borisenko AV, Steinke D, Telfer AC, Perez KHJ, Sones JE, Young MR, Levesque-Beaudin V, Sobel CN, Abrahamyan A, Bessonov K, Blagoev G, deWaard SL, Ho C, Ivanova NV, Layton KKS, Lu L, Manjunath R, McKeown JTA, Milton MA, Miskie R, Monkhouse N, Naik S, Nikolova N, Pentinsaari M, Prosser SWJ, Radulovici AE, Steinke C, Warne CP, Hebert PDN. 2019. A reference library for Canadian invertebrates with 1.5 million barcodes, voucher specimens, and DNA samples. Scientific Data 6: 308. Doi: 10.1038/s41597-019-0320-2.

Dincă V, Dapporto L, Somervuo P, Vodă R, Cuvelier S, Gascoigne-Pees M, Huemer P, Mutanen M, Hebert PDN, Vila R. 2021. High resolution DNA barcode library for European butterflies reveals continental patterns of mitochondrial genetic diversity. Communications Biology 4: 315. Doi: 10.1038/s42003-021-01834-7.

Disney RHL. 1983. Scuttle Flies: Diptera, Phoridae (except Megaselia). Handbooks for the Identification of British Insects Vol. 10, Part 6. Royal Entomological Society of London. London.

Disney RHL. 1994. Scuttle Flies: The Phoridae. Springer Dordrecht. Doi: 10.1007/978-94-011-1288-8.

Disney RHL. 2008. Natural history of the Scuttle Fly, Megaselia scalaris. The Annual Review of Entomology 53: 39–60. Doi: 10.1146/annurev.ento.53.103106.093415.

Disney RHL. 2013. An unusually rich scuttle fly fauna (Diptera, Phoridae) from north of the Arctic Circle in the Kola Peninsula, N.W. Russia. ZooKeys 342: 45–74. Doi: 10.3897/zookeys.342.5772.

Disney RHL & Withers P. 2009. A new species of Pseudacteon Coquillett (Diptera, Phoridae) and a new key to the European species. Fragmenta Faunistica 52(1): 1–12.

Disney RHL, Erzinclioglu Y, de C Henshaw D, Unwin D, Withers P, Woods A. 1982. Collecting methods and adequacy of attempted fauna surveys, with reference to the Diptera. Field Studies 5: 607–621. Doi: 10.5281/zenodo.10742404.

Engel MS, Ceríaco LMP, Daniel GM, Dellapé PM, Löbl I, Marinov M, Reis RE, Young MT, Dubois A, Agarwal I, Lehmann P, Alvarado M, Alvarez N, Andreone F, Araujo-Vieira K, Ascher JS, Baêta D, Baldo D, Bandeira SA, Barden P, Barrasso DA, Bendifallah L, Bockmann FA, Böhme W, Borkent A, Brandão CRF, Busack SD, Bybee SM, Channing A, Chatzimanolis S, Christenhusz MJM, Crisci JV, D’elía G, da Costa LM, Davis SR, de Lucena CAS, Deuve T, Elizalde SF, Faivovich J, Farooq H, Ferguson AW, Gippoliti S, Gonçalves FMP, Gonzalez VH, Greenbaum E, Hinojosa-Díaz IA, Ineich I, Jiang J, Kahono S, Kury AB, Lucinda PHF, Lynch JD, Malécot V, Marques MP, Marris JWM, Mckellar RC, Mendes LF, Nihei SS, Nishikawa K, Ohler A, Orrico VGD, Ota H, Paiva J, Parrinha D, Pauwels OSG, Pereyra MO, Pestana LB, Pinheiro PDP, Prendini L, Prokop J, Rasmussen C, Rödel M-O, Rodrigues MT, Rodríguez SM, Salatnaya H, Sampaio í, Sánchez-García A, Shebl MA, Santos BS, Solórzano-Kraemer MM, Sousa ACA, Stoev P, Teta P, Trape J-F, Van-Dúnem Dos Santos C, Vasudevan K, Vink Cj, Vogel G, Wagner P, Wappler T, Ware Jl, Wedmann S, Zacharie Ck. 2021. The taxonomic impediment: a shortage of taxonomists, not the lack of technical approaches. Zoological Journal of Linnean Society 193: 381–387. Doi: 10.1093/zoolinnean/zlab072.

Finnish Biodiversity Information Facility. FinBIF. 2024. Laji.fi. https://laji.fi/en. Accessed July 2024.

German Barcode of Life. GBOL III: Dark Taxa. https://gbol.bolgermany.de/gbol3/. Accessed November 2023.

Godfray HCJ. 2002. Challenges for taxonomy. Nature 417: 17–19.

Gotô T. 1986. Systematic Study of the Genus Phora Latreille from Japan (Diptera, Phoridae) V*. Kontyu, Tokyo 54(1): 128–142.

Gregory TR. 2005. DNA barcoding does not compete with taxonomy. Nature 434: 1067. Doi: 10.1038/4341067b.

Grundmann B & Kappert J. 2023. The Phoridae (Diptera) of NE-Westphalia: a field study over five years. Fragmenta Faunistica 66(1): 15–36. Doi: 10.3161/00159301FF2023.66.1.015.

Hartop E, Lee L, Srivathsan A, Jones M, Peña-Aguilera P, Ovaskainen O, Roslin T, Meier R. 2024. Resolving biology’s dark matter: species richness, spatiotemporal distribution, and community composition of a dark taxon. BMC Biology 22:215. Doi: 10.1186/s12915-024-02010-z.

Hartop E, Srivathsan A, Ronquist F, Meier R. 2022. Towards Large-Scale Integrative Taxonomy (LIT): Resolving the Data Conundrum for Dark Taxa. Systematic Biology 71(6): 1404–1422.

Hausmann A, Godfray HCJ, Huemer P, Mutanen M, Rougerie R, van Nieukerken EJ, Ratnasingham S, Hebert PDN. 2013. Genetic Patterns in European Geometrid Moths Revealed by the Barcode Index Number (BIN) System. PLoS One 8(12): e84518. Doi: 10.1371/journal.pone.0084518.

Hausmann A, Krogmann L, Peters RS, Rduch V, Schmidt S. 2020. GBOL III: dark taxa. From: GBOL III: Dark Taxa - iBOL Barcode Bulletin.

Hebert PDN, Braukmann TWA, Prosser SWJ, Ratnasingham S, deWaard JR, Ivanova NV, Janzen DH, Hallwachs W, Naik S, Sones JE, Zakharov EV. 2018. A Sequel to Sanger: amplicon sequencing that scales. BMC Genomics 19: 219. Doi: 10.1186/s12864-018-4611-3.

Hebert PDN, Cywinska A, Ball SL, deWaard JR. 2003. Biological identifications through DNA barcodes. Proceedings of the Royal Society B: Biological Sciences 270: 313–321. Doi: 10.1098/rspb.2002.2218.

Hebert PDN, Penton EH, Burns JM, Janzen DH, Hallwachs W. 2004. Ten species in one: DNA barcoding reveals cryptic species in the neotropical skipper butterfly Astraptes fulgerator. Proceedings of the National Academy of Sciences 101(41): 14812–14817. Doi: 10.1073/pnas.0406166101.

Hulsen T. 2022. DeepVenn – a web application for the creation of area-proportional Venn diagrams using the deep learning framework Tensorflow.js. 2210.04597. Doi: 10.48550/arXiv.2210.04597.

International Barcode of Life. https://ibol.org/. Accessed December 2023.

Kahanpää J. 2014. Checklist of the families Lonchopteridae and Phoridae of Finland (Insecta, Diptera). ZooKeys 441: 213–223. Doi: 10.3897/zookeys.441.7197.

Karlsson D, Hartop E, Forshage M, Jaschhof M, Ronquist F. 2020. The Swedish Malaise Trap Project: A 15 Year Retrospective on a Countrywide Insect Inventory. Biodiversity Data Journal 8: e47255. Doi: 10.3897/BDJ.8.e47255.

Kekkonen M & Hebert PDN. 2014. DNA barcode-based delineation of putative species: efficient start for taxonomic workflows. Molecular Ecology Resources 14(4): 706–715. Doi: 10.1111/1755-0998.12233.

Kekkonen M, Mutanen M, Kaila L, Nieminen M, Hebert PDN. 2015. Delineating Species with DNA Barcodes: A Case of Taxon Dependent Method Performance in Moths. PLoS ONE 10(4): e0122481. Doi: 10.1371/journal.pone.0122481.

Lipscomb D, Platnick N, Wheeler Q. 2003. The intellectual content of taxonomy: a comment on DNA taxonomy. Trends in Ecology and Evolution 18(2): 65–66. Doi: 10.1016/S0169-5347(02)00060-5.

Liu G-C & Yang M. 2016. A taxonomic revision of the genus Diplonevra Lioy (Diptera: Phoridae) from China. Zootaxa 4205(1): 031–051. Doi: 10.11646/zootaxa.4205.1.3.

Marshall SA. 2012. Flies: The Natural History and Diversity of Diptera. 616 p. Firefly Books. Richmond Hill. Ontario.

Maschwitz U, Weissflog A, Seebauer S, Disney RHL, Witte V. 2008. Studies of European Ant Decapitating Flies (Diptera: Phoridae): I. Releasers and Phenology of Parasitism of Pseudacteon formicarium. Sociobiology Vol 51(1): 127–140.

Meierotto S, Sharkey MJ, Janzen DH, Hallwachs W, Hebert PDN, Chapman EG, Smith MA. 2019. A revolutionary protocol to describe understudied hyperdiverse taxa and overcome taxonomic impediment. Deutsche Entomologische Zeitschrift 66(2): 119–145. Doi: 10.3897/dez.66.34683.

Morinière J, Balke M, Doczkal D, Geiger MF, Hardulak LA, Haszprunar G, Hausmann A, Hendrich L, Regalado L, Rulik B, Schmidt S, Wägele J-W, Hebert PDN. 2019. A DNA barcode library for 5,200 German flies and midges (Insecta: Diptera) and its implications for metabarcoding-based biomonitoring. Molecular Ecology Resources 19(4): 900–928. Doi: 10.1111/1755-0998.13022.

Morinière J, Hendrich L, Hausmann A, Hebert P, Haszprunar G, Gruppe A. 2014. Barcoding Fauna Bavarica: 78% of the Neuropterida Fauna Barcoded! PLoS One 9(10): e109719. Doi: 10.1371/journal.pone.0109719.

Morrison LW, Porter SD. 2005. Testing for population-level impacts of introduced Pseudacteon tricuspis flies, phorid parasitoids of Solenopsis invicta fire ants. Biological Control 33: 9–19. Doi: 10.1016/j.biocontrol.2005.01.004.

Page RDM. 2016. DNA barcoding and taxonomy: dark taxa and dark texts. Philosophical Transactions of the Royal Society B: Biological Sciences 371(1702): 20150334. Doi: 10.1098/rstb.2015.0334.

Park D-S, Foottit R, Maw E, Hebert PDN. 2011. Barcoding Bugs: DNA-Based Identification of the True Bugs (Insecta: Hemiptera: Heteroptera). PLoS One 6(4): e18749. Doi: 10.1371/journal.pone.0018749.

Pentinsaari M, Hebert PDN, Mutanen M. 2014. Barcoding beetles: a regional survey of 1872 species reveals high identification success and unusually deep interspecific divergences. PLos One 9: e108651. Doi: 10.1371/journal.pone.0108651.

Phorid Catalog. 2024. https://www.phorid.net/pcat/index.php. Accessed December 2024.

Pohjoismäki JLO, Kahanpää J, Mutanen M. 2016. DNA Barcodes for the Northern European Tachnid Flies (Diptera: Tachinidae). PLoS One 11(11): e0164933. 10.1371/journal.pone.0164933.

Porter SD. 1998. Biology and behavior of Pseudacteon decapitating flies (Diptera: Phoridae) that parasitize Solenopsis fire ants (Hymenoptera: Formicidae). The Florida Entomologist, Vol. 81, No. 3: 292–309. Doi: 10.2307/3495920.

Ratnasingham S & Hebert PDN. 2007. BOLD: The Barcode of Life Data System (www.barcodinglife.org). Molecular Ecology Notes 7(3): 355–364. Doi: 10.1111/j.1471-8286.2007.01678.x.

Ratnasingham S & Hebert PDN. 2013. A DNA-Based Registry for All Animal Species: The Barcode Index Number (BIN) System. PLoS One 8(8): e66213. Doi: 10.1371/journal.pone.0066213.

Roslin T, Somervuo P, Pentinsaari M, Hebert PDN, Agda J, Ahlroth P, Anttonen P, Aspi J, Blagoev G, Blanco S, Chan D, Clayhills T, deWaard J, deWaard S, Elliot T, Elo R, Haapala S, Helve E, Ilmonen J, Hirvonen P, Ho C, Itämies J, Ivanov V, Jakovlev J, Juslén A, Jussila R, Kahanpää J, Kaila L, Kaitila J-P, Kakko A, Kakko I, Karhu A, Karjalainen S, Kjaerandsen J, Koskinen J, Laasonen EM, Laasonen L, Laine E, Lampila P, Levesque-Beaudin V, Lu L, Lähteenaro M, Majuri P, Malmberg S, Manjunath R, Martikainen P, Mattila J, McKeown J, Metsälä P, Miklasevskaja M, Miller M, Miskie R, Muinonen A, Mukkala V-P, Naik S, Nikolova N, Nupponen K, Ovaskainen O, Österblad I, Paasivirta L, Pajunen T, Parkko P, Paukkunen J, Penttinen R, Perez K, Pohjoismäki J, Prosser S, Raekunnas M, Rahulan M, Rannisto M, Ratnasingham S, Raukko P, Rinne A, Rintala T, Romo S, Salmela J, Salokannel J, Savolainen R, Schulman L, Sihvonen P, Soliman D, Sones J, Steinke C, Ståhls G, Tabell J, Tiusanen M, Várkonyi G, Vesterinen EJ, Viitanen E, Vikberg V, Viitasaari M, Vilen J, Warne C, Wei C, Winqvist K, Zakharov E, Mutanen M. 2022. A molecular-based identification resource for the arthropods of Finland. Molecular Ecology Resources 22(2): 803–822. Doi: 10.1111/1755-0998.13510.

Rubinoff D, Cameron S, Will K. 2006. A Genomic Perspective on the Shortcomings of Mitochondrial DNA for “Barcoding” Identification. Journal of Heredity 97(6): 581–594. Doi: 10.1093/jhered/esl036.

Schmidt S, Schmid-Egger C, Morinière J, Haszprunar G, Hebert PDN. 2015. DNA barcoding largely supports 250 years of classical taxonomy: identifications for Central European bees (Hymenoptera, Apoidea partim). Molecular Ecology Resources 15(4): 985–1000. Doi: 10.1111/1755-0998.12363.

Schmitz H, Beyer E, Delage A. 1981. 33. Phoridae. Die Fliegen Der Palaearktischen region. Stuttgart. Germany.

Sharkey MJ, Baker A, McCluskey K, Smith A, Naik S, Ratnasingham S, Manjunath R, Perez K, Sones J, D’Souza M, St. Jacques B, Hajibabaei M, Whitfield J, Arias D, Solis A, Metz M, Burns J, Zuñiga R, Phillips-Rodriguez E, Espinoza B, Chacon I, Hebert P, Hallwachs W, Janzen D. 2023. Minimalist revision of Mesochorus Gravenhost, 1829 (Hymenoptera: Ichneumonidae: Mesochorinae) from Área de conservación Guanacaste, Costa Rica, with 158 new species and host records for 129 species. Revista De Biología Tropical 71(S2): e53316. Doi: 10.15517/rev.biol.trop..v71iS2.56316.

Sharkey MJ, Baker A, McCluskey K, Smith A, Naik S, Ratnasingham S, Manjunath R, Perez K, Sones J, D’Souza M, St. Jacques B, Hebert P, Hallwachs W, Janzen D. 2021b. Addendum to a minimalist revision of Costa Rican Braconidae: 28 new species and 23 host records. ZooKeys 1075: 77–136. Doi: 10.3897/zookeys.1075.72197.

Sharkey MJ, Janzen DH, Hallwachs W, Chapman EG, Smith MA, Dapkey T, Brown A, Ratnasingham S, Naik S, Manjunath R, Perez K, Milton M, Hebert P, Shaw SR, Kittel RN, Solis MA, Metz MA, Goldstein PZ, Brown JW, Quicke DLJ, van Achtenberg C, Brown BV, Burns JM. 2021a. Minimalist revision and description of 403 new species in 11 subfamilies of Costa Rican braconid parasitoid wasps, including host records for 219 species. ZooKeys 1013: 1–665. Doi: 10.3897/zookeys.1013.55600.

Stireman JO, Cerretti P, O’Hara JE, Moulton JK. 2021. Extraordinary diversification of the “bristle flies” (Diptera: Tachinidae) and its underlying causes. Biological Journal of the Linnean Society 133: 216–236. Doi: 10.1093/biolinnean/blab010.

Srivathsan A, Ang Y, Heraty JM, Hwang WS, Jusoh WFA, Kutty SN, Puniamoorthy J, Yeo D, Roslin T, Meier R. 2023. Convergence of dominance and neglect in flying insect diversity. Nature Ecology and Evolution 7: 1012–1021.

Srivathsan A, Hartop E, Puniamoorthy J, Lee WT, Kutty SN, Kurina O, Meier R. 2019. Rapid, large-scale species discovery in hyperdiverse taxa using 1D MinION sequencing. BMC Biology 17: 96. Doi: 10.1186/s12915-019-0706-9.

Taberlet P, Coissac E, Hajibabaei M, Rieseberg LH. 2012. Environmental DNA. Molecular Ecology 21: 1789–1793. Doi: 10.1111/j.1365-294X.2012.05542.x.

van Klink R, Sheard JK, Høye TT, Roslin T, Do Nascimento LA, Bauer S. 2024. Towards a toolkit for global insect biodiversity monitoring. Philosophical Transactions Royal Society Publishing B: Biological Sciences 379: 20230101. Doi: 10.1098/rstb.2023.0101.

Virgilio M, Backeljau T, Nevado B, De Meyer M. 2010. Comparative performances of DNA barcoding across insect orders. BMC Bioinformatics 11: 206. Doi: 10.1186/1471-2105-11-206.

Wirta H, Várkonyi G, Rasmussen C, Kaartinen R, Schmidt NM, Hebert PDN, Barták M, Blagoev G, Disney H, Ertl S, Gjelstrup P, Gwiazdowicz DJ, Huldén L, Ilmonen J, Jakovlev J, Jaschhof M, Kahanpää J, Kankaanpää T, Krogh PH, Labbee R, Lettner C, Michelsen V, Nielsen SA, Nielsen TR, Paasivirta L, Pedersen S, Pohjoismäki J, Salmela J, Vilkamaa P, Väre H, von Tschirnhaus M, Roslin T. 2016. Establishing a community-wide DNA barcode library as a new tool for arctic research. Molecular Ecology Resources 16: 809–822. Doi: 10.1111/1755-0998.12489.

