## Supplementary material for "Megabarcoding dark taxa – Assessing the utility of mass DNA barcoding for phorid fly species discovery": S-Table

### Supplementary information

S-Table 1. Specimen details containing sample ID, BIN, species, sampling site, coordinates, collection date and trap type used (WiT = Window Trap, MaT = Malaise Trap).

| Sample ID | BIN | Species | Site | Lat | Lon | Date | Trap |
|---|---|---|---|---|---|---|---|
| ZMUO.068002 | BOLD:ABA7002 | *Phora tincta* | Joroinen | 62.301 | 27.699 | 27-Jun-2021 | WiT |
| ZMUO.068004 | BOLD:AFI7403 | *Phora artifrons* | Joroinen | 62.301 | 27.699 | 27-Jun-2021 | WiT |
| ZMUO.068005 | BOLD:ABA7002 | *Phora tincta* | Joroinen | 62.301 | 27.699 | 27-Jun-2021 | WiT |
| ZMUO.068007 | BOLD:ACQ9655 | *Anevrina unispinosa* | Joroinen | 62.301 | 27.699 | 27-Jun-2021 | WiT |
| ZMUO.068009 | BOLD:ABA7002 | *Phora tincta* | Joroinen | 62.301 | 27.699 | 27-Jun-2021 | WiT |
| ZMUO.068010 | BOLD:ACD2261 | *Phora artifrons* | Joroinen | 62.301 | 27.699 | 27-Jun-2021 | WiT |
| ZMUO.068013 | BOLD:ABA7002 | *Phora tincta* | Joroinen | 62.301 | 27.699 | 27-Jun-2021 | WiT |
| ZMUO.068014 | BOLD:ABA7002 | *Phora tincta* | Joroinen | 62.301 | 27.699 | 27-Jun-2021 | WiT |
| ZMUO.068016 | BOLD:ABA7002 | *Phora tincta* | Joroinen | 62.301 | 27.699 | 27-Jun-2021 | WiT |
| ZMUO.068019 | BOLD:ABA7002 | *Phora tincta* | Joroinen | 62.301 | 27.699 | 27-Jun-2021 | WiT |
| ZMUO.068026 | BOLD:ABA7002 | *Phora tincta* | Joroinen | 62.301 | 27.699 | 27-Jun-2021 | WiT |
| ZMUO.068028 | BOLD:ACD2261 | *Phora artifrons* | Joroinen | 62.304 | 27.71 | 27-Jun-2021 | WiT |
| ZMUO.068034 | BOLD:ACD2261 | *Phora artifrons* | Joroinen | 62.304 | 27.71 | 27-Jun-2021 | WiT |
| ZMUO.068035 | BOLD:ABA7002 | *Phora tincta* | Joroinen | 62.304 | 27.71 | 27-Jun-2021 | WiT |
| ZMUO.068037 | BOLD:ABA7002 | *Phora tincta* | Joroinen | 62.304 | 27.71 | 27-Jun-2021 | WiT |
| ZMUO.068038 | BOLD:ABA7002 | *Phora tincta* | Joroinen | 62.304 | 27.71 | 27-Jun-2021 | WiT |
| ZMUO.068039 | BOLD:ABA7001 | *Diplonevra abdominalis* | Joroinen | 62.304 | 27.71 | 27-Jun-2021 | WiT |
| ZMUO.068046 | BOLD:ABA7002 | *Phora tincta* | Joroinen | 62.304 | 27.71 | 27-Jun-2021 | WiT |
| ZMUO.068050 | BOLD:ACD2261 | *Phora artifrons* | Joroinen | 62.304 | 27.71 | 27-Jun-2021 | WiT |
| ZMUO.068054 | BOLD:ABA7002 | *Phora tincta* | Joroinen | 62.304 | 27.71 | 27-Jun-2021 | WiT |
| ZMUO.068055 | BOLD:ACD2261 | *Phora artifrons* | Joroinen | 62.304 | 27.71 | 27-Jun-2021 | WiT |
| ZMUO.068058 | BOLD:ACD2261 | *Phora artifrons* | Joroinen | 62.304 | 27.71 | 27-Jun-2021 | WiT |
| ZMUO.068062 | BOLD:ABA7002 | *Phora tincta* | Joroinen | 62.306 | 27.706 | 27-Jun-2021 | WiT |
| ZMUO.068064 | BOLD:ACN6664 | *Phora pubipes* | Joroinen | 62.306 | 27.706 | 27-Jun-2021 | WiT |
| ZMUO.068066 | BOLD:ACN6664 | *Phora pubipes* | Joroinen | 62.306 | 27.706 | 27-Jun-2021 | WiT |
| ZMUO.068068 | BOLD:AEB8779 | *Phora convergens* | Joroinen | 62.306 | 27.706 | 27-Jun-2021 | WiT |
| ZMUO.068069 | BOLD:ABA7002 | *Phora tincta* | Joroinen | 62.306 | 27.706 | 27-Jun-2021 | WiT |

| | | | | | | | |
|---|---|---|---|---|---|---|---|
| ZMUO.068075 | BOLD:ACN6664 | *Phora pubipes* | Joroinen | 62.306 | 27.706 | 27-Jun-2021 | WiT |
| ZMUO.068076 | BOLD:ABA7002 | *Phora tincta* | Joroinen | 62.306 | 27.706 | 27-Jun-2021 | WiT |
| ZMUO.068077 | BOLD:ABA7002 | *Phora tincta* | Joroinen | 62.306 | 27.706 | 27-Jun-2021 | WiT |
| ZMUO.068085 | BOLD:ACN6664 | *Phora pubipes* | Joroinen | 62.306 | 27.706 | 27-Jun-2021 | WiT |
| ZMUO.068092 | BOLD:ABA7002 | *Phora tincta* | Joroinen | 62.306 | 27.706 | 27-Jun-2021 | WiT |
| ZMUO.068100 | BOLD:ABA7002 | *Phora tincta* | Joroinen | 62.306 | 27.706 | 27-Jun-2021 | WiT |
| ZMUO.068102 | BOLD:AEB8779 | *Phora convergens* | Joroinen | 62.306 | 27.706 | 27-Jun-2021 | WiT |
| ZMUO.068104 | BOLD:ACN6664 | *Phora pubipes* | Joroinen | 62.306 | 27.706 | 27-Jun-2021 | WiT |
| ZMUO.068105 | BOLD:ABA7002 | *Phora tincta* | Joroinen | 62.306 | 27.706 | 27-Jun-2021 | WiT |
| ZMUO.068109 | BOLD:ACN6664 | *Phora pubipes* | Joroinen | 62.306 | 27.706 | 27-Jun-2021 | WiT |
| ZMUO.068111 | BOLD:ACN6664 | *Phora pubipes* | Joroinen | 62.306 | 27.706 | 27-Jun-2021 | WiT |
| ZMUO.068118 | BOLD:ACN6664 | *Phora pubipes* | Joroinen | 62.306 | 27.706 | 27-Jun-2021 | WiT |
| ZMUO.068124 | BOLD:ABA7002 | *Phora tincta* | Joroinen | 62.306 | 27.706 | 27-Jun-2021 | WiT |
| ZMUO.068126 | BOLD:AEB8779 | *Phora convergens* | Joroinen | 62.306 | 27.706 | 27-Jun-2021 | WiT |
| ZMUO.068128 | BOLD:ABA7002 | *Phora tincta* | Joroinen | 62.306 | 27.706 | 27-Jun-2021 | WiT |
| ZMUO.068129 | BOLD:ABA7002 | *Phora tincta* | Joroinen | 62.306 | 27.706 | 27-Jun-2021 | WiT |
| ZMUO.068130 | BOLD:ABA7002 | *Phora tincta* | Joroinen | 62.306 | 27.706 | 27-Jun-2021 | WiT |
| ZMUO.068141 | BOLD:ABA7002 | *Phora tincta* | Joroinen | 62.306 | 27.706 | 27-Jun-2021 | WiT |
| ZMUO.068153 | BOLD:ACN6664 | *Phora pubipes* | Joroinen | 62.306 | 27.706 | 27-Jun-2021 | WiT |
| ZMUO.068159 | BOLD:ABA7002 | *Phora tincta* | Joroinen | 62.306 | 27.706 | 27-Jun-2021 | WiT |
| ZMUO.068162 | BOLD:ABA7002 | *Phora tincta* | Joroinen | 62.306 | 27.706 | 27-Jun-2021 | WiT |
| ZMUO.068165 | BOLD:ABA7002 | *Phora tincta* | Joroinen | 62.306 | 27.706 | 27-Jun-2021 | WiT |
| ZMUO.068168 | BOLD:ABA7002 | *Phora tincta* | Joroinen | 62.306 | 27.706 | 27-Jun-2021 | WiT |
| ZMUO.068170 | BOLD:ACN6664 | *Phora pubipes* | Joroinen | 62.306 | 27.706 | 27-Jun-2021 | WiT |
| ZMUO.068174 | BOLD:ACN6664 | *Phora pubipes* | Joroinen | 62.306 | 27.706 | 27-Jun-2021 | WiT |
| ZMUO.068180 | BOLD:ABA7002 | *Phora tincta* | Joroinen | 62.306 | 27.706 | 27-Jun-2021 | WiT |
| ZMUO.068188 | BOLD:AEB8779 | *Phora convergens* | Joroinen | 62.306 | 27.706 | 27-Jun-2021 | WiT |
| ZMUO.068189 | BOLD:ABA7002 | *Phora tincta* | Joroinen | 62.306 | 27.706 | 27-Jun-2021 | WiT |
| ZMUO.068190 | BOLD:ACN6664 | *Phora pubipes* | Joroinen | 62.306 | 27.706 | 27-Jun-2021 | WiT |
| ZMUO.068192 | BOLD:ACN6664 | *Phora pubipes* | Joroinen | 62.306 | 27.706 | 27-Jun-2021 | WiT |
| ZMUO.068193 | BOLD:ABA7002 | *Phora tincta* | Joroinen | 62.306 | 27.706 | 27-Jun-2021 | WiT |
| ZMUO.068197 | BOLD:ABA7002 | *Phora tincta* | Joroinen | 62.306 | 27.706 | 27-Jun-2021 | WiT |
| ZMUO.068202 | BOLD:ABA7002 | *Phora tincta* | Joroinen | 62.306 | 27.706 | 27-Jun-2021 | WiT |
| ZMUO.068208 | BOLD:ACN6664 | *Phora pubipes* | Joroinen | 62.306 | 27.706 | 27-Jun-2021 | WiT |
| ZMUO.068213 | BOLD:ACN6664 | *Phora pubipes* | Joroinen | 62.306 | 27.706 | 27-Jun-2021 | WiT |
| ZMUO.068225 | BOLD:ABA7002 | *Phora tincta* | Joroinen | 62.306 | 27.706 | 27-Jun-2021 | WiT |
| ZMUO.068229 | BOLD:ACN6664 | *Phora pubipes* | Joroinen | 62.306 | 27.706 | 27-Jun-2021 | WiT |
| ZMUO.068234 | BOLD:ACN6664 | *Phora pubipes* | Joroinen | 62.306 | 27.706 | 27-Jun-2021 | WiT |
| ZMUO.068239 | BOLD:AFE7212 | *Phora holosericea* | Joroinen | 62.306 | 27.706 | 27-Jun-2021 | WiT |
| ZMUO.068241 | BOLD:AEB8779 | *Phora convergens* | Joroinen | 62.306 | 27.706 | 27-Jun-2021 | WiT |
| ZMUO.068242 | BOLD:ABA7002 | *Phora tincta* | Joroinen | 62.306 | 27.706 | 27-Jun-2021 | WiT |
| ZMUO.068245 | BOLD:ACN6664 | *Phora pubipes* | Joroinen | 62.306 | 27.706 | 27-Jun-2021 | WiT |
| ZMUO.068246 | BOLD:ABA7002 | *Phora tincta* | Joroinen | 62.306 | 27.706 | 27-Jun-2021 | WiT |
| ZMUO.068250 | BOLD:ABA7002 | *Phora tincta* | Joroinen | 62.306 | 27.706 | 27-Jun-2021 | WiT |
| ZMUO.068251 | BOLD:ACN6664 | *Phora pubipes* | Joroinen | 62.306 | 27.706 | 27-Jun-2021 | WiT |
| ZMUO.068259 | BOLD:ACN6664 | *Phora pubipes* | Joroinen | 62.306 | 27.706 | 27-Jun-2021 | WiT |
| ZMUO.068271 | BOLD:ABA7002 | *Phora tincta* | Joroinen | 62.306 | 27.706 | 27-Jun-2021 | WiT |
| ZMUO.068272 | BOLD:ABA7002 | *Phora tincta* | Joroinen | 62.306 | 27.706 | 27-Jun-2021 | WiT |

| | | | | | | | |
|---|---|---|---|---|---|---|---|
| ZMUO.068273 | BOLD:ACQ9655 | *Anevrina unispinosa* | Joroinen | 62.306 | 27.706 | 27-Jun-2021 | WiT |
| ZMUO.068276 | BOLD:ACN6664 | *Phora pubipes* | Joroinen | 62.306 | 27.706 | 27-Jun-2021 | WiT |
| ZMUO.068288 | BOLD:ABA7002 | *Phora tincta* | Joroinen | 62.306 | 27.706 | 27-Jun-2021 | WiT |
| ZMUO.068290 | BOLD:ABA7002 | *Phora tincta* | Joroinen | 62.306 | 27.706 | 27-Jun-2021 | WiT |
| ZMUO.068291 | BOLD:AEB8779 | *Phora convergens* | Joroinen | 62.306 | 27.706 | 27-Jun-2021 | WiT |
| ZMUO.068292 | BOLD:ACN6664 | *Phora pubipes* | Joroinen | 62.306 | 27.706 | 27-Jun-2021 | WiT |
| ZMUO.068294 | BOLD:ABA7002 | *Phora tincta* | Joroinen | 62.306 | 27.706 | 27-Jun-2021 | WiT |
| ZMUO.068297 | BOLD:ABA7002 | *Phora tincta* | Joroinen | 62.306 | 27.706 | 27-Jun-2021 | WiT |
| ZMUO.068303 | BOLD:ABA7002 | *Phora tincta* | Joroinen | 62.306 | 27.706 | 27-Jun-2021 | WiT |
| ZMUO.068309 | BOLD:ABA7002 | *Phora tincta* | Joroinen | 62.306 | 27.706 | 27-Jun-2021 | WiT |
| ZMUO.068311 | BOLD:ABA7002 | *Phora tincta* | Joroinen | 62.306 | 27.706 | 27-Jun-2021 | WiT |
| ZMUO.068313 | BOLD:ACN6664 | *Phora pubipes* | Joroinen | 62.306 | 27.706 | 27-Jun-2021 | WiT |
| ZMUO.068316 | BOLD:ACN6664 | *Phora pubipes* | Joroinen | 62.306 | 27.706 | 27-Jun-2021 | WiT |
| ZMUO.068318 | BOLD:ABA7002 | *Phora tincta* | Joroinen | 62.306 | 27.706 | 27-Jun-2021 | WiT |
| ZMUO.068321 | BOLD:ACN6664 | *Phora pubipes* | Joroinen | 62.306 | 27.706 | 27-Jun-2021 | WiT |
| ZMUO.068328 | BOLD:ACN6664 | *Phora pubipes* | Joroinen | 62.306 | 27.706 | 27-Jun-2021 | WiT |
| ZMUO.068329 | BOLD:AEB8779 | *Phora convergens* | Joroinen | 62.306 | 27.706 | 27-Jun-2021 | WiT |
| ZMUO.068332 | BOLD:ACN6664 | *Phora pubipes* | Joroinen | 62.306 | 27.706 | 27-Jun-2021 | WiT |
| ZMUO.068338 | BOLD:ABA7002 | *Phora tincta* | Joroinen | 62.306 | 27.706 | 27-Jun-2021 | WiT |
| ZMUO.068341 | BOLD:ACN6664 | *Phora pubipes* | Joroinen | 62.304 | 27.71 | 27-Jun-2021 | WiT |
| ZMUO.068344 | BOLD:AEB8779 | *Phora convergens* | Joroinen | 62.304 | 27.71 | 27-Jun-2021 | WiT |
| ZMUO.068346 | BOLD:ACI8210 | *Phora convergens* | Joroinen | 62.304 | 27.71 | 27-Jun-2021 | WiT |
| ZMUO.068350 | BOLD:AEB8779 | *Phora convergens* | Joroinen | 62.304 | 27.71 | 27-Jun-2021 | WiT |
| ZMUO.068352 | BOLD:ABA7002 | *Phora tincta* | Joroinen | 62.304 | 27.71 | 27-Jun-2021 | WiT |
| ZMUO.068353 | BOLD:ACI8210 | *Phora convergens* | Joroinen | 62.304 | 27.71 | 27-Jun-2021 | WiT |
| ZMUO.068359 | BOLD:ACN6664 | *Phora pubipes* | Joroinen | 62.304 | 27.71 | 27-Jun-2021 | WiT |
| ZMUO.068363 | BOLD:ABA7002 | *Phora tincta* | Joroinen | 62.304 | 27.71 | 27-Jun-2021 | WiT |
| ZMUO.068367 | BOLD:ACN6664 | *Phora pubipes* | Joroinen | 62.304 | 27.71 | 27-Jun-2021 | WiT |
| ZMUO.068374 | BOLD:ABA7002 | *Phora tincta* | Joroinen | 62.304 | 27.71 | 27-Jun-2021 | WiT |
| ZMUO.068376 | BOLD:ACN6664 | *Phora pubipes* | Joroinen | 62.304 | 27.71 | 27-Jun-2021 | WiT |
| ZMUO.068383 | BOLD:ACN6664 | *Phora pubipes* | Joroinen | 62.304 | 27.71 | 27-Jun-2021 | WiT |
| ZMUO.068387 | BOLD:ACN6664 | *Phora pubipes* | Joroinen | 62.304 | 27.71 | 27-Jun-2021 | WiT |
| ZMUO.068388 | BOLD:ACN6664 | *Phora pubipes* | Joroinen | 62.304 | 27.71 | 27-Jun-2021 | WiT |
| ZMUO.068395 | BOLD:ACN6664 | *Phora pubipes* | Joroinen | 62.304 | 27.71 | 27-Jun-2021 | WiT |
| ZMUO.068399 | BOLD:ACN6664 | *Phora pubipes* | Joroinen | 62.304 | 27.71 | 27-Jun-2021 | WiT |
| ZMUO.068402 | BOLD:ABA7002 | *Phora tincta* | Joroinen | 62.304 | 27.71 | 27-Jun-2021 | WiT |
| ZMUO.068403 | BOLD:ACN6664 | *Phora pubipes* | Joroinen | 62.304 | 27.71 | 27-Jun-2021 | WiT |
| ZMUO.068405 | BOLD:ABA7002 | *Phora tincta* | Joroinen | 62.304 | 27.71 | 27-Jun-2021 | WiT |
| ZMUO.068411 | BOLD:ACN6664 | *Phora pubipes* | Joroinen | 62.304 | 27.71 | 27-Jun-2021 | WiT |
| ZMUO.068415 | BOLD:ACN6664 | *Phora pubipes* | Joroinen | 62.304 | 27.71 | 27-Jun-2021 | WiT |
| ZMUO.068418 | BOLD:ACN6664 | *Phora pubipes* | Joroinen | 62.304 | 27.71 | 27-Jun-2021 | WiT |
| ZMUO.068428 | BOLD:ACN6664 | *Phora pubipes* | Joroinen | 62.304 | 27.71 | 27-Jun-2021 | WiT |
| ZMUO.068434 | BOLD:ACN6664 | *Phora pubipes* | Joroinen | 62.304 | 27.71 | 27-Jun-2021 | WiT |
| ZMUO.068435 | BOLD:ABA7002 | *Phora tincta* | Joroinen | 62.304 | 27.71 | 27-Jun-2021 | WiT |
| ZMUO.068439 | BOLD:ACN6664 | *Phora pubipes* | Joroinen | 62.304 | 27.71 | 27-Jun-2021 | WiT |
| ZMUO.068450 | BOLD:ACI8210 | *Phora convergens* | Joroinen | 62.304 | 27.71 | 27-Jun-2021 | WiT |
| ZMUO.068451 | BOLD:ACN6664 | *Phora pubipes* | Joroinen | 62.304 | 27.71 | 27-Jun-2021 | WiT |
| ZMUO.068458 | BOLD:ACN6664 | *Phora pubipes* | Joroinen | 62.304 | 27.71 | 27-Jun-2021 | WiT |

| | | | | | | | |
|---|---|---|---|---|---|---|---|
| ZMUO.068462 | BOLD:ACN6664 | *Phora pubipes* | Joroinen | 62.304 | 27.71 | 27-Jun-2021 | WiT |
| ZMUO.068471 | BOLD:ACN6664 | *Phora pubipes* | Joroinen | 62.304 | 27.71 | 27-Jun-2021 | WiT |
| ZMUO.068480 | BOLD:AEB8779 | *Phora convergens* | Joroinen | 62.304 | 27.71 | 27-Jun-2021 | WiT |
| ZMUO.068482 | BOLD:ACI8210 | *Phora convergens* | Joroinen | 62.304 | 27.71 | 27-Jun-2021 | WiT |
| ZMUO.068487 | BOLD:ACN6664 | *Phora pubipes* | Joroinen | 62.304 | 27.71 | 27-Jun-2021 | WiT |
| ZMUO.068488 | BOLD:ACN6664 | *Phora pubipes* | Joroinen | 62.304 | 27.71 | 27-Jun-2021 | WiT |
| ZMUO.068501 | BOLD:ACN6664 | *Phora pubipes* | Joroinen | 62.304 | 27.709 | 27-Jun-2021 | WiT |
| ZMUO.068502 | BOLD:ACN6664 | *Phora pubipes* | Joroinen | 62.304 | 27.709 | 27-Jun-2021 | WiT |
| ZMUO.068504 | BOLD:ACN6664 | *Phora pubipes* | Joroinen | 62.304 | 27.709 | 27-Jun-2021 | WiT |
| ZMUO.068505 | BOLD:AEB8779 | *Phora convergens* | Joroinen | 62.304 | 27.709 | 27-Jun-2021 | WiT |
| ZMUO.068521 | BOLD:ACN6664 | *Phora pubipes* | Joroinen | 62.304 | 27.709 | 27-Jun-2021 | WiT |
| ZMUO.068528 | BOLD:ACN6664 | *Phora pubipes* | Joroinen | 62.304 | 27.709 | 27-Jun-2021 | WiT |
| ZMUO.068533 | BOLD:AEB8779 | *Phora convergens* | Joroinen | 62.304 | 27.709 | 27-Jun-2021 | WiT |
| ZMUO.068535 | BOLD:ACN6664 | *Phora pubipes* | Joroinen | 62.304 | 27.709 | 27-Jun-2021 | WiT |
| ZMUO.068536 | BOLD:ABA7002 | *Phora tincta* | Joroinen | 62.304 | 27.709 | 27-Jun-2021 | WiT |
| ZMUO.068540 | BOLD:ACN6664 | *Phora pubipes* | Joroinen | 62.304 | 27.709 | 27-Jun-2021 | WiT |
| ZMUO.068542 | BOLD:ACN6664 | *Phora pubipes* | Joroinen | 62.304 | 27.709 | 27-Jun-2021 | WiT |
| ZMUO.068546 | BOLD:ACN6664 | *Phora pubipes* | Joroinen | 62.304 | 27.709 | 27-Jun-2021 | WiT |
| ZMUO.068553 | BOLD:ABA7002 | *Phora tincta* | Joroinen | 62.304 | 27.709 | 27-Jun-2021 | WiT |
| ZMUO.068554 | BOLD:ACN6664 | *Phora pubipes* | Joroinen | 62.304 | 27.709 | 27-Jun-2021 | WiT |
| ZMUO.068558 | BOLD:ACN6664 | *Phora pubipes* | Joroinen | 62.304 | 27.709 | 27-Jun-2021 | WiT |
| ZMUO.068567 | BOLD:ACN6664 | *Phora pubipes* | Joroinen | 62.304 | 27.709 | 27-Jun-2021 | WiT |
| ZMUO.068568 | BOLD:ACN6664 | *Phora pubipes* | Joroinen | 62.304 | 27.709 | 27-Jun-2021 | WiT |
| ZMUO.068571 | BOLD:ACN6664 | *Phora pubipes* | Joroinen | 62.304 | 27.709 | 27-Jun-2021 | WiT |
| ZMUO.068573 | BOLD:ACN6664 | *Phora pubipes* | Joroinen | 62.304 | 27.709 | 27-Jun-2021 | WiT |
| ZMUO.068574 | BOLD:ABA7002 | *Phora tincta* | Joroinen | 62.304 | 27.709 | 27-Jun-2021 | WiT |
| ZMUO.068576 | BOLD:ACN6664 | *Phora pubipes* | Joroinen | 62.304 | 27.709 | 27-Jun-2021 | WiT |
| ZMUO.068582 | BOLD:ACN6664 | *Phora pubipes* | Joroinen | 62.304 | 27.709 | 27-Jun-2021 | WiT |
| ZMUO.068583 | BOLD:ACN6664 | *Phora pubipes* | Joroinen | 62.304 | 27.709 | 27-Jun-2021 | WiT |
| ZMUO.068592 | BOLD:AEB8779 | *Phora convergens* | Joroinen | 62.304 | 27.709 | 27-Jun-2021 | WiT |
| ZMUO.068593 | BOLD:ACN6664 | *Phora pubipes* | Joroinen | 62.304 | 27.709 | 27-Jun-2021 | WiT |
| ZMUO.068597 | BOLD:ACN6664 | *Phora pubipes* | Joroinen | 62.304 | 27.709 | 27-Jun-2021 | WiT |
| ZMUO.068599 | BOLD:ACN6664 | *Phora pubipes* | Joroinen | 62.304 | 27.709 | 27-Jun-2021 | WiT |
| ZMUO.068601 | BOLD:ACN6664 | *Phora pubipes* | Joroinen | 62.304 | 27.709 | 27-Jun-2021 | WiT |
| ZMUO.068605 | BOLD:ABA7002 | *Phora tincta* | Joroinen | 62.304 | 27.709 | 27-Jun-2021 | WiT |
| ZMUO.068614 | BOLD:AEB8779 | *Phora convergens* | Joroinen | 62.304 | 27.709 | 27-Jun-2021 | WiT |
| ZMUO.068625 | BOLD:ABA7002 | *Phora tincta* | Joroinen | 62.304 | 27.709 | 27-Jun-2021 | WiT |
| ZMUO.068626 | BOLD:ABA7002 | *Phora tincta* | Joroinen | 62.304 | 27.709 | 27-Jun-2021 | WiT |
| ZMUO.068631 | BOLD:ACN6664 | *Phora pubipes* | Joroinen | 62.304 | 27.709 | 27-Jun-2021 | WiT |
| ZMUO.068637 | BOLD:ACN6664 | *Phora pubipes* | Joroinen | 62.304 | 27.709 | 27-Jun-2021 | WiT |
| ZMUO.068639 | BOLD:ACN6664 | *Phora pubipes* | Joroinen | 62.304 | 27.709 | 27-Jun-2021 | WiT |
| ZMUO.068640 | BOLD:ACN6664 | *Phora pubipes* | Joroinen | 62.304 | 27.709 | 27-Jun-2021 | WiT |
| ZMUO.068646 | BOLD:ACN6664 | *Phora pubipes* | Joroinen | 62.304 | 27.709 | 27-Jun-2021 | WiT |
| ZMUO.068651 | BOLD:AEB8779 | *Phora convergens* | Joroinen | 62.304 | 27.709 | 27-Jun-2021 | WiT |
| ZMUO.068653 | BOLD:ACQ9655 | *Anevrina unispinosa* | Joroinen | 62.304 | 27.709 | 27-Jun-2021 | WiT |
| ZMUO.068658 | BOLD:ACN6664 | *Phora pubipes* | Joroinen | 62.304 | 27.709 | 27-Jun-2021 | WiT |
| ZMUO.068669 | BOLD:ACN6664 | *Phora pubipes* | Joroinen | 62.304 | 27.709 | 27-Jun-2021 | WiT |
| ZMUO.068670 | BOLD:ACN6664 | *Phora pubipes* | Joroinen | 62.304 | 27.709 | 27-Jun-2021 | WiT |

| | | | | | | | |
|---|---|---|---|---|---|---|---|
| ZMUO.068680 | BOLD:ACN6664 | *Phora pubipes* | Joroinen | 62.304 | 27.709 | 27-Jun-2021 | WiT |
| ZMUO.068684 | BOLD:ACN6664 | *Phora pubipes* | Joroinen | 62.304 | 27.709 | 27-Jun-2021 | WiT |
| ZMUO.068687 | BOLD:AEB8779 | *Phora convergens* | Joroinen | 62.3 | 27.701 | 27-Jun-2021 | WiT |
| ZMUO.068688 | BOLD:ABA7002 | *Phora tincta* | Joroinen | 62.3 | 27.701 | 27-Jun-2021 | WiT |
| ZMUO.068692 | BOLD:AEB8779 | *Phora convergens* | Joroinen | 62.3 | 27.701 | 27-Jun-2021 | WiT |
| ZMUO.068695 | BOLD:ABA7002 | *Phora tincta* | Joroinen | 62.3 | 27.701 | 27-Jun-2021 | WiT |
| ZMUO.068700 | BOLD:ACI8210 | *Phora convergens* | Joroinen | 62.3 | 27.701 | 27-Jun-2021 | WiT |
| ZMUO.068707 | BOLD:AEB8779 | *Phora convergens* | Joroinen | 62.3 | 27.701 | 27-Jun-2021 | WiT |
| ZMUO.068711 | BOLD:AEB8779 | *Phora convergens* | Joroinen | 62.3 | 27.701 | 27-Jun-2021 | WiT |
| ZMUO.068712 | BOLD:ABA7002 | *Phora tincta* | Joroinen | 62.3 | 27.701 | 27-Jun-2021 | WiT |
| ZMUO.068714 | BOLD:ACN6664 | *Phora pubipes* | Joroinen | 62.3 | 27.701 | 27-Jun-2021 | WiT |
| ZMUO.068729 | BOLD:AEB8779 | *Phora convergens* | Joroinen | 62.3 | 27.701 | 27-Jun-2021 | WiT |
| ZMUO.068743 | BOLD:ACI8210 | *Phora convergens* | Joroinen | 62.3 | 27.701 | 27-Jun-2021 | WiT |
| ZMUO.068744 | BOLD:AFE7212 | *Phora holosericea* | Joroinen | 62.3 | 27.701 | 27-Jun-2021 | WiT |
| ZMUO.068746 | BOLD:ACN6664 | *Phora pubipes* | Joroinen | 62.3 | 27.701 | 27-Jun-2021 | WiT |
| ZMUO.068748 | BOLD:ABA7002 | *Phora tincta* | Joroinen | 62.3 | 27.701 | 27-Jun-2021 | WiT |
| ZMUO.068756 | BOLD:ACN6664 | *Phora pubipes* | Joroinen | 62.3 | 27.701 | 27-Jun-2021 | WiT |
| ZMUO.068757 | BOLD:ABA7002 | *Phora tincta* | Joroinen | 62.3 | 27.701 | 27-Jun-2021 | WiT |
| ZMUO.068760 | BOLD:ACN6664 | *Phora pubipes* | Joroinen | 62.3 | 27.701 | 27-Jun-2021 | WiT |
| ZMUO.068763 | BOLD:ACN6664 | *Phora pubipes* | Joroinen | 62.3 | 27.701 | 27-Jun-2021 | WiT |
| ZMUO.068764 | BOLD:ABA7002 | *Phora tincta* | Joroinen | 62.3 | 27.701 | 27-Jun-2021 | WiT |
| ZMUO.068766 | BOLD:AEB8779 | *Phora convergens* | Joroinen | 62.3 | 27.701 | 27-Jun-2021 | WiT |
| ZMUO.068769 | BOLD:ACN6664 | *Phora pubipes* | Joroinen | 62.3 | 27.701 | 27-Jun-2021 | WiT |
| ZMUO.068774 | BOLD:AEB8779 | *Phora convergens* | Joroinen | 62.3 | 27.701 | 27-Jun-2021 | WiT |
| ZMUO.068777 | BOLD:ABA7002 | *Phora tincta* | Joroinen | 62.3 | 27.701 | 27-Jun-2021 | WiT |
| ZMUO.068780 | BOLD:ACN6664 | *Phora pubipes* | Joroinen | 62.3 | 27.701 | 27-Jun-2021 | WiT |
| ZMUO.068784 | BOLD:ACN6664 | *Phora pubipes* | Joroinen | 62.3 | 27.701 | 27-Jun-2021 | WiT |
| ZMUO.068787 | BOLD:AEB8779 | *Phora convergens* | Joroinen | 62.3 | 27.701 | 27-Jun-2021 | WiT |
| ZMUO.068788 | BOLD:AEB8779 | *Phora convergens* | Joroinen | 62.3 | 27.701 | 27-Jun-2021 | WiT |
| ZMUO.068789 | BOLD:AEB8779 | *Phora convergens* | Joroinen | 62.3 | 27.701 | 27-Jun-2021 | WiT |
| ZMUO.068792 | BOLD:ACN6664 | *Phora pubipes* | Joroinen | 62.3 | 27.701 | 27-Jun-2021 | WiT |
| ZMUO.068793 | BOLD:ACN6664 | *Phora pubipes* | Joroinen | 62.3 | 27.701 | 27-Jun-2021 | WiT |
| ZMUO.068796 | BOLD:ACN6664 | *Phora pubipes* | Joroinen | 62.3 | 27.701 | 27-Jun-2021 | WiT |
| ZMUO.068800 | BOLD:ABA7002 | *Phora tincta* | Joroinen | 62.3 | 27.701 | 27-Jun-2021 | WiT |
| ZMUO.068802 | BOLD:ACN6664 | *Phora pubipes* | Joroinen | 62.3 | 27.701 | 27-Jun-2021 | WiT |
| ZMUO.068804 | BOLD:ABA7002 | *Phora tincta* | Joroinen | 62.3 | 27.701 | 27-Jun-2021 | WiT |
| ZMUO.068805 | BOLD:ACN6664 | *Phora pubipes* | Joroinen | 62.3 | 27.701 | 27-Jun-2021 | WiT |
| ZMUO.068807 | BOLD:ACN6664 | *Phora pubipes* | Joroinen | 62.3 | 27.701 | 27-Jun-2021 | WiT |
| ZMUO.068808 | BOLD:ACN6664 | *Phora pubipes* | Joroinen | 62.3 | 27.701 | 27-Jun-2021 | WiT |
| ZMUO.068810 | BOLD:AEB8779 | *Phora convergens* | Joroinen | 62.3 | 27.701 | 27-Jun-2021 | WiT |
| ZMUO.068812 | BOLD:ABA7002 | *Phora tincta* | Joroinen | 62.3 | 27.701 | 27-Jun-2021 | WiT |
| ZMUO.068813 | BOLD:AEB8779 | *Phora convergens* | Joroinen | 62.3 | 27.701 | 27-Jun-2021 | WiT |
| ZMUO.068814 | BOLD:ABA7002 | *Phora tincta* | Joroinen | 62.3 | 27.701 | 27-Jun-2021 | WiT |
| ZMUO.068819 | BOLD:ABA7002 | *Phora tincta* | Joroinen | 62.3 | 27.701 | 27-Jun-2021 | WiT |
| ZMUO.068821 | BOLD:ACN6664 | *Phora pubipes* | Joroinen | 62.3 | 27.701 | 27-Jun-2021 | WiT |
| ZMUO.068822 | BOLD:ACN6664 | *Phora pubipes* | Joroinen | 62.3 | 27.701 | 27-Jun-2021 | WiT |
| ZMUO.068823 | BOLD:AEB8779 | *Phora convergens* | Joroinen | 62.3 | 27.701 | 27-Jun-2021 | WiT |
| ZMUO.068827 | BOLD:AEB8779 | *Phora convergens* | Joroinen | 62.3 | 27.701 | 27-Jun-2021 | WiT |

| | | | | | | | |
|---|---|---|---|---|---|---|---|
| ZMUO.068829 | BOLD:ACN6664 | *Phora pubipes* | Joroinen | 62.3 | 27.701 | 27-Jun-2021 | WiT |
| ZMUO.068832 | BOLD:ABA7002 | *Phora tincta* | Joroinen | 62.3 | 27.701 | 27-Jun-2021 | WiT |
| ZMUO.068834 | BOLD:ACN6664 | *Phora pubipes* | Joroinen | 62.3 | 27.701 | 27-Jun-2021 | WiT |
| ZMUO.068837 | BOLD:ABA7002 | *Phora tincta* | Joroinen | 62.3 | 27.701 | 27-Jun-2021 | WiT |
| ZMUO.068841 | BOLD:ACN6664 | *Phora pubipes* | Joroinen | 62.3 | 27.701 | 27-Jun-2021 | WiT |
| ZMUO.068843 | BOLD:ACN6664 | *Phora pubipes* | Joroinen | 62.3 | 27.701 | 27-Jun-2021 | WiT |
| ZMUO.068844 | BOLD:ACN6664 | *Phora pubipes* | Joroinen | 62.3 | 27.701 | 27-Jun-2021 | WiT |
| ZMUO.068853 | BOLD:ACN6664 | *Phora pubipes* | Joroinen | 62.3 | 27.701 | 27-Jun-2021 | WiT |
| ZMUO.068855 | BOLD:ACN6664 | *Phora pubipes* | Joroinen | 62.3 | 27.701 | 27-Jun-2021 | WiT |
| ZMUO.068856 | BOLD:ACN6664 | *Phora pubipes* | Joroinen | 62.3 | 27.701 | 27-Jun-2021 | WiT |
| ZMUO.068857 | BOLD:ABA7002 | *Phora tincta* | Joroinen | 62.3 | 27.701 | 27-Jun-2021 | WiT |
| ZMUO.068860 | BOLD:ACN6664 | *Phora pubipes* | Joroinen | 62.3 | 27.701 | 27-Jun-2021 | WiT |
| ZMUO.068861 | BOLD:ABA7002 | *Phora tincta* | Joroinen | 62.3 | 27.701 | 27-Jun-2021 | WiT |
| ZMUO.068866 | BOLD:ABA7002 | *Phora tincta* | Joroinen | 62.3 | 27.701 | 27-Jun-2021 | WiT |
| ZMUO.068868 | BOLD:ACN6664 | *Phora pubipes* | Joroinen | 62.3 | 27.701 | 27-Jun-2021 | WiT |
| ZMUO.068869 | BOLD:ABA7002 | *Phora tincta* | Joroinen | 62.3 | 27.701 | 27-Jun-2021 | WiT |
| ZMUO.068870 | BOLD:AEB8779 | *Phora convergens* | Joroinen | 62.3 | 27.701 | 27-Jun-2021 | WiT |
| ZMUO.068874 | BOLD:ACN6664 | *Phora pubipes* | Joroinen | 62.3 | 27.701 | 27-Jun-2021 | WiT |
| ZMUO.068875 | BOLD:ACN6664 | *Phora pubipes* | Joroinen | 62.3 | 27.701 | 27-Jun-2021 | WiT |
| ZMUO.068877 | BOLD:ABA7002 | *Phora tincta* | Joroinen | 62.3 | 27.701 | 27-Jun-2021 | WiT |
| ZMUO.068881 | BOLD:ABA7002 | *Phora tincta* | Joroinen | 62.3 | 27.701 | 27-Jun-2021 | WiT |
| ZMUO.068882 | BOLD:ABA7002 | *Phora tincta* | Joroinen | 62.3 | 27.701 | 27-Jun-2021 | WiT |
| ZMUO.068883 | BOLD:AEB8779 | *Phora convergens* | Joroinen | 62.3 | 27.701 | 27-Jun-2021 | WiT |
| ZMUO.068884 | BOLD:ACN6664 | *Phora pubipes* | Joroinen | 62.3 | 27.701 | 27-Jun-2021 | WiT |
| ZMUO.068885 | BOLD:ABA7002 | *Phora tincta* | Joroinen | 62.3 | 27.701 | 27-Jun-2021 | WiT |
| ZMUO.068886 | BOLD:ACN6664 | *Phora pubipes* | Joroinen | 62.3 | 27.701 | 27-Jun-2021 | WiT |
| ZMUO.068889 | BOLD:ACB4872 | *Phora atra* | Joroinen | 62.3 | 27.701 | 27-Jun-2021 | WiT |
| ZMUO.068890 | BOLD:AEB8779 | *Phora convergens* | Joroinen | 62.3 | 27.701 | 27-Jun-2021 | WiT |
| ZMUO.068895 | BOLD:AEB8779 | *Phora convergens* | Joroinen | 62.3 | 27.701 | 27-Jun-2021 | WiT |
| ZMUO.068897 | BOLD:ACN6664 | *Phora pubipes* | Joroinen | 62.3 | 27.701 | 27-Jun-2021 | WiT |
| ZMUO.068898 | BOLD:ABA7002 | *Phora tincta* | Joroinen | 62.3 | 27.701 | 27-Jun-2021 | WiT |
| ZMUO.068901 | BOLD:ACN6664 | *Phora pubipes* | Joroinen | 62.3 | 27.701 | 27-Jun-2021 | WiT |
| ZMUO.068905 | BOLD:ABA7002 | *Phora tincta* | Joroinen | 62.3 | 27.701 | 27-Jun-2021 | WiT |
| ZMUO.068907 | BOLD:ABA7002 | *Phora tincta* | Joroinen | 62.3 | 27.701 | 27-Jun-2021 | WiT |
| ZMUO.068912 | BOLD:ABA7002 | *Phora tincta* | Joroinen | 62.3 | 27.701 | 27-Jun-2021 | WiT |
| ZMUO.068913 | BOLD:ABA7002 | *Phora tincta* | Joroinen | 62.3 | 27.701 | 27-Jun-2021 | WiT |
| ZMUO.068917 | BOLD:ACN6664 | *Phora pubipes* | Joroinen | 62.3 | 27.701 | 27-Jun-2021 | WiT |
| ZMUO.068919 | BOLD:ABA7002 | *Phora tincta* | Joroinen | 62.3 | 27.701 | 27-Jun-2021 | WiT |
| ZMUO.068921 | BOLD:ACN6664 | *Phora pubipes* | Joroinen | 62.3 | 27.701 | 27-Jun-2021 | WiT |
| ZMUO.068922 | BOLD:ACN6664 | *Phora pubipes* | Joroinen | 62.3 | 27.701 | 27-Jun-2021 | WiT |
| ZMUO.068923 | BOLD:ABA7002 | *Phora tincta* | Joroinen | 62.3 | 27.701 | 27-Jun-2021 | WiT |
| ZMUO.068924 | BOLD:ACN6664 | *Phora pubipes* | Joroinen | 62.3 | 27.701 | 27-Jun-2021 | WiT |
| ZMUO.068926 | BOLD:AEB8779 | *Phora convergens* | Joroinen | 62.3 | 27.701 | 27-Jun-2021 | WiT |
| ZMUO.068929 | BOLD:AEB8779 | *Phora convergens* | Joroinen | 62.3 | 27.701 | 27-Jun-2021 | WiT |
| ZMUO.068932 | BOLD:ABA7002 | *Phora tincta* | Joroinen | 62.3 | 27.701 | 27-Jun-2021 | WiT |
| ZMUO.068933 | BOLD:ACN6664 | *Phora pubipes* | Joroinen | 62.3 | 27.701 | 27-Jun-2021 | WiT |
| ZMUO.068934 | BOLD:ACN6664 | *Phora pubipes* | Joroinen | 62.3 | 27.701 | 27-Jun-2021 | WiT |
| ZMUO.068935 | BOLD:ACN6664 | *Phora pubipes* | Joroinen | 62.3 | 27.701 | 27-Jun-2021 | WiT |

| | | | | | | | |
|---|---|---|---|---|---|---|---|
| ZMUO.068936 | BOLD:ACN6664 | *Phora pubipes* | Joroinen | 62.3 | 27.701 | 27-Jun-2021 | WiT |
| ZMUO.068937 | BOLD:ACN6664 | *Phora pubipes* | Joroinen | 62.3 | 27.701 | 27-Jun-2021 | WiT |
| ZMUO.068938 | BOLD:ABA7002 | *Phora tincta* | Joroinen | 62.3 | 27.701 | 27-Jun-2021 | WiT |
| ZMUO.068945 | BOLD:ABA7002 | *Phora tincta* | Joroinen | 62.3 | 27.701 | 27-Jun-2021 | WiT |
| ZMUO.068949 | BOLD:ABA7002 | *Phora tincta* | Joroinen | 62.3 | 27.701 | 27-Jun-2021 | WiT |
| ZMUO.069039 | BOLD:ADM5152 | *Triphleba antricola* | Inkoo | 60.069 | 23.863 | 13-Jul-2020 | MaT |
| ZMUO.069053 | BOLD:ACB3701 | *Diplonevra concinna* | Inkoo | 60.069 | 23.863 | 22-Aug-2020 | MaT |
| ZMUO.069322 | BOLD:ABA7001 | *Diplonevra glabra* | Kristiinankaupunki | 62.282 | 21.409 | 23-Jun-2020 | MaT |
| ZMUO.069328 | BOLD:ABA7001 | *Diplonevra glabra* | Kristiinankaupunki | 62.282 | 21.409 | 23-Jun-2020 | MaT |
| ZMUO.069338 | BOLD:ABA7001 | *Diplonevra glabra* | Kristiinankaupunki | 62.282 | 21.409 | 23-Jun-2020 | MaT |
| ZMUO.069345 | BOLD:ABA7001 | *Diplonevra glabra* | Kristiinankaupunki | 62.282 | 21.409 | 23-Jun-2020 | MaT |
| ZMUO.069359 | BOLD:ACE2645 | *Conicera floricola* | Kristiinankaupunki | 62.282 | 21.409 | 23-Jun-2020 | MaT |
| ZMUO.069385 | BOLD:ADM8144 | *Phora obscura* | Kristiinankaupunki | 62.282 | 21.409 | 23-Jun-2020 | MaT |
| ZMUO.069392 | BOLD:ABA7002 | *Phora tincta* | Kristiinankaupunki | 62.282 | 21.409 | 23-Jun-2020 | MaT |
| ZMUO.069412 | BOLD:AAG3236 | *Diplonevra nitidula* | Kristiinankaupunki | 62.282 | 21.409 | 23-Jun-2020 | MaT |
| ZMUO.069423 | BOLD:ABA7002 | *Phora tincta* | Kristiinankaupunki | 62.282 | 21.409 | 23-Jun-2020 | MaT |
| ZMUO.069441 | BOLD:ACO9836 | *Triphleba distinguenda* | Kristiinankaupunki | 62.282 | 21.409 | 14-Jul-2020 | MaT |
| ZMUO.069468 | BOLD:AAG3236 | *Diplonevra nitidula* | Kristiinankaupunki | 62.282 | 21.409 | 14-Jul-2020 | MaT |
| ZMUO.069485 | BOLD:ACO9836 | *Triphleba distinguenda* | Kristiinankaupunki | 62.282 | 21.409 | 14-Jul-2020 | MaT |
| ZMUO.069486 | BOLD:ACO9836 | *Triphleba distinguenda* | Kristiinankaupunki | 62.282 | 21.409 | 14-Jul-2020 | MaT |
| ZMUO.069501 | BOLD:ACB4872 | *Phora atra* | Kristiinankaupunki | 62.282 | 21.409 | 14-Jul-2020 | MaT |
| ZMUO.069515 | BOLD:AAG3236 | *Diplonevra nitidula* | Kristiinankaupunki | 62.282 | 21.409 | 14-Jul-2020 | MaT |
| ZMUO.069516 | BOLD:ACO9836 | *Triphleba distinguenda* | Kristiinankaupunki | 62.282 | 21.409 | 14-Jul-2020 | MaT |
| ZMUO.069528 | BOLD:ADM5152 | *Triphleba antricola* | Kristiinankaupunki | 62.282 | 21.409 | 14-Jul-2020 | MaT |
| ZMUO.069536 | BOLD:AAG3236 | *Diplonevra nitidula* | Kristiinankaupunki | 62.282 | 21.409 | 14-Jul-2020 | MaT |
| ZMUO.069558 | BOLD:ACO9836 | *Triphleba distinguenda* | Kristiinankaupunki | 62.282 | 21.409 | 14-Jul-2020 | MaT |
| ZMUO.069560 | BOLD:ACO9836 | *Triphleba distinguenda* | Kristiinankaupunki | 62.282 | 21.409 | 04-Aug-2020 | MaT |
| ZMUO.069564 | BOLD:ACO9836 | *Triphleba distinguenda* | Kristiinankaupunki | 62.282 | 21.409 | 04-Aug-2020 | MaT |
| ZMUO.069565 | BOLD:ACO9836 | *Triphleba distinguenda* | Kristiinankaupunki | 62.282 | 21.409 | 04-Aug-2020 | MaT |
| ZMUO.069568 | BOLD:ADM5152 | *Triphleba antricola* | Kristiinankaupunki | 62.282 | 21.409 | 04-Aug-2020 | MaT |
| ZMUO.069570 | BOLD:ACO9836 | *Triphleba distinguenda* | Kristiinankaupunki | 62.282 | 21.409 | 04-Aug-2020 | MaT |
| ZMUO.069571 | BOLD:ACO9836 | *Triphleba distinguenda* | Kristiinankaupunki | 62.282 | 21.409 | 04-Aug-2020 | MaT |
| ZMUO.069579 | BOLD:ACD2464 | *Borophaga carinifrons* | Kristiinankaupunki | 62.282 | 21.409 | 04-Aug-2020 | MaT |
| ZMUO.069580 | BOLD:ACD2464 | *Borophaga carinifrons* | Kristiinankaupunki | 62.282 | 21.409 | 04-Aug-2020 | MaT |
| ZMUO.069585 | BOLD:ACO9836 | *Triphleba distinguenda* | Kristiinankaupunki | 62.282 | 21.409 | 04-Aug-2020 | MaT |
| ZMUO.069586 | BOLD:ACO9836 | *Triphleba distinguenda* | Kristiinankaupunki | 62.282 | 21.409 | 04-Aug-2020 | MaT |
| ZMUO.069587 | BOLD:ACD2464 | *Borophaga carinifrons* | Kristiinankaupunki | 62.282 | 21.409 | 04-Aug-2020 | MaT |
| ZMUO.069588 | BOLD:ACD2464 | *Borophaga carinifrons* | Kristiinankaupunki | 62.282 | 21.409 | 04-Aug-2020 | MaT |
| ZMUO.069589 | BOLD:ADM5152 | *Triphleba antricola* | Kristiinankaupunki | 62.282 | 21.409 | 04-Aug-2020 | MaT |
| ZMUO.069590 | BOLD:ACO9836 | *Triphleba distinguenda* | Kristiinankaupunki | 62.282 | 21.409 | 04-Aug-2020 | MaT |
| ZMUO.069595 | BOLD:ACD2464 | *Borophaga carinifrons* | Kristiinankaupunki | 62.282 | 21.409 | 04-Aug-2020 | MaT |
| ZMUO.069597 | BOLD:ACO9836 | *Triphleba distinguenda* | Kristiinankaupunki | 62.282 | 21.409 | 04-Aug-2020 | MaT |
| ZMUO.069603 | BOLD:ACO9836 | *Triphleba distinguenda* | Kristiinankaupunki | 62.282 | 21.409 | 04-Aug-2020 | MaT |
| ZMUO.069606 | BOLD:ADM5152 | *Triphleba antricola* | Kristiinankaupunki | 62.282 | 21.409 | 04-Aug-2020 | MaT |
| ZMUO.069611 | BOLD:ACO9836 | *Triphleba distinguenda* | Kristiinankaupunki | 62.282 | 21.409 | 04-Aug-2020 | MaT |
| ZMUO.069613 | BOLD:ACO9836 | *Triphleba distinguenda* | Kristiinankaupunki | 62.282 | 21.409 | 04-Aug-2020 | MaT |
| ZMUO.069615 | BOLD:ACD2464 | *Borophaga carinifrons* | Kristiinankaupunki | 62.282 | 21.409 | 04-Aug-2020 | MaT |
| ZMUO.069621 | BOLD:ACD2464 | *Borophaga carinifrons* | Kristiinankaupunki | 62.282 | 21.409 | 04-Aug-2020 | MaT |

| | | | | | | | |
|---|---|---|---|---|---|---|---|
| ZMUO.069623 | BOLD:ACO9836 | *Triphleba distinguenda* | Kristiinankaupunki | 62.282 | 21.409 | 04-Aug-2020 | MaT |
| ZMUO.069624 | BOLD:ACO9836 | *Triphleba distinguenda* | Kristiinankaupunki | 62.282 | 21.409 | 04-Aug-2020 | MaT |
| ZMUO.069627 | BOLD:ACO9836 | *Triphleba distinguenda* | Kristiinankaupunki | 62.282 | 21.409 | 04-Aug-2020 | MaT |
| ZMUO.069629 | BOLD:ACO9836 | *Triphleba distinguenda* | Kristiinankaupunki | 62.282 | 21.409 | 04-Aug-2020 | MaT |
| ZMUO.069630 | BOLD:ACB3701 | *Diplonevra concinna* | Kristiinankaupunki | 62.282 | 21.409 | 04-Aug-2020 | MaT |
| ZMUO.069638 | BOLD:ACO9836 | *Triphleba distinguenda* | Kristiinankaupunki | 62.282 | 21.409 | 04-Aug-2020 | MaT |
| ZMUO.069652 | BOLD:ACO9836 | *Triphleba distinguenda* | Kristiinankaupunki | 62.282 | 21.409 | 04-Aug-2020 | MaT |
| ZMUO.069653 | BOLD:ACO9836 | *Triphleba distinguenda* | Kristiinankaupunki | 62.282 | 21.409 | 04-Aug-2020 | MaT |
| ZMUO.069656 | BOLD:ACD2464 | *Borophaga carinifrons* | Kristiinankaupunki | 62.282 | 21.409 | 23-Aug-2020 | MaT |
| ZMUO.069657 | BOLD:ACD2464 | *Borophaga carinifrons* | Kristiinankaupunki | 62.282 | 21.409 | 23-Aug-2020 | MaT |
| ZMUO.069658 | BOLD:ACD2464 | *Borophaga carinifrons* | Kristiinankaupunki | 62.282 | 21.409 | 23-Aug-2020 | MaT |
| ZMUO.069659 | BOLD:ACD2464 | *Borophaga carinifrons* | Kristiinankaupunki | 62.282 | 21.409 | 23-Aug-2020 | MaT |
| ZMUO.069662 | BOLD:ACQ3921 | *Phora stictica* | Kristiinankaupunki | 62.282 | 21.409 | 23-Aug-2020 | MaT |
| ZMUO.069663 | BOLD:ACD2464 | *Borophaga carinifrons* | Kristiinankaupunki | 62.282 | 21.409 | 23-Aug-2020 | MaT |
| ZMUO.069664 | BOLD:ACB3701 | *Diplonevra concinna* | Kristiinankaupunki | 62.282 | 21.409 | 23-Aug-2020 | MaT |
| ZMUO.069665 | BOLD:ACD2464 | *Borophaga carinifrons* | Kristiinankaupunki | 62.282 | 21.409 | 23-Aug-2020 | MaT |
| ZMUO.069667 | BOLD:ACD2464 | *Borophaga carinifrons* | Kristiinankaupunki | 62.282 | 21.409 | 23-Aug-2020 | MaT |
| ZMUO.069668 | BOLD:ACD2464 | *Borophaga carinifrons* | Kristiinankaupunki | 62.282 | 21.409 | 23-Aug-2020 | MaT |
| ZMUO.069669 | BOLD:ACD2464 | *Borophaga carinifrons* | Kristiinankaupunki | 62.282 | 21.409 | 23-Aug-2020 | MaT |
| ZMUO.069670 | BOLD:ACD2464 | *Borophaga carinifrons* | Kristiinankaupunki | 62.282 | 21.409 | 23-Aug-2020 | MaT |
| ZMUO.069671 | BOLD:ACD2464 | *Borophaga carinifrons* | Kristiinankaupunki | 62.282 | 21.409 | 23-Aug-2020 | MaT |
| ZMUO.069672 | BOLD:ACD2464 | *Borophaga carinifrons* | Kristiinankaupunki | 62.282 | 21.409 | 23-Aug-2020 | MaT |
| ZMUO.069674 | BOLD:ACD2464 | *Borophaga carinifrons* | Kristiinankaupunki | 62.282 | 21.409 | 23-Aug-2020 | MaT |
| ZMUO.069678 | BOLD:ACD2464 | *Borophaga carinifrons* | Kristiinankaupunki | 62.282 | 21.409 | 23-Aug-2020 | MaT |
| ZMUO.069679 | BOLD:ACD2464 | *Borophaga carinifrons* | Kristiinankaupunki | 62.282 | 21.409 | 23-Aug-2020 | MaT |
| ZMUO.069681 | BOLD:ACO9836 | *Triphleba distinguenda* | Kristiinankaupunki | 62.282 | 21.409 | 23-Aug-2020 | MaT |
| ZMUO.069683 | BOLD:ACD2464 | *Borophaga carinifrons* | Kristiinankaupunki | 62.282 | 21.409 | 23-Aug-2020 | MaT |
| ZMUO.069685 | BOLD:ACD2464 | *Borophaga carinifrons* | Kristiinankaupunki | 62.282 | 21.409 | 23-Aug-2020 | MaT |
| ZMUO.069686 | BOLD:ACD2464 | *Borophaga carinifrons* | Kristiinankaupunki | 62.282 | 21.409 | 23-Aug-2020 | MaT |
| ZMUO.069687 | BOLD:ACD2464 | *Borophaga carinifrons* | Kristiinankaupunki | 62.282 | 21.409 | 23-Aug-2020 | MaT |
| ZMUO.069688 | BOLD:ACO9836 | *Triphleba distinguenda* | Kristiinankaupunki | 62.282 | 21.409 | 23-Aug-2020 | MaT |
| ZMUO.069689 | BOLD:ACD2464 | *Borophaga carinifrons* | Kristiinankaupunki | 62.282 | 21.409 | 23-Aug-2020 | MaT |
| ZMUO.069690 | BOLD:ACD2464 | *Borophaga carinifrons* | Kristiinankaupunki | 62.282 | 21.409 | 23-Aug-2020 | MaT |
| ZMUO.069691 | BOLD:ACD2464 | *Borophaga carinifrons* | Kristiinankaupunki | 62.282 | 21.409 | 23-Aug-2020 | MaT |
| ZMUO.069692 | BOLD:ADM5152 | *Triphleba antricola* | Kristiinankaupunki | 62.282 | 21.409 | 23-Aug-2020 | MaT |
| ZMUO.069696 | BOLD:ACD2464 | *Borophaga carinifrons* | Kristiinankaupunki | 62.282 | 21.409 | 23-Aug-2020 | MaT |
| ZMUO.069697 | BOLD:ACB3701 | *Diplonevra concinna* | Kristiinankaupunki | 62.282 | 21.409 | 23-Aug-2020 | MaT |
| ZMUO.069703 | BOLD:ABA7002 | *Phora tincta* | Turku | 60.432 | 22.162 | 22-Jun-2020 | MaT |
| ZMUO.069704 | BOLD:ABA7002 | *Phora tincta* | Turku | 60.432 | 22.162 | 22-Jun-2020 | MaT |
| ZMUO.069716 | BOLD:ABA7002 | *Phora tincta* | Turku | 60.432 | 22.162 | 22-Jun-2020 | MaT |
| ZMUO.069719 | BOLD:ABA7002 | *Phora tincta* | Turku | 60.432 | 22.162 | 22-Jun-2020 | MaT |
| ZMUO.069721 | BOLD:ADM5152 | *Triphleba antricola* | Turku | 60.432 | 22.162 | 22-Jun-2020 | MaT |
| ZMUO.069733 | BOLD:ADM5152 | *Triphleba antricola* | Turku | 60.432 | 22.162 | 22-Jun-2020 | MaT |
| ZMUO.069735 | BOLD:ABA7001 | *Diplonevra glabra* | Turku | 60.432 | 22.162 | 22-Jun-2020 | MaT |
| ZMUO.069784 | BOLD:ADM5152 | *Triphleba antricola* | Turku | 60.432 | 22.162 | 22-Jun-2020 | MaT |
| ZMUO.069787 | BOLD:ACD2261 | *Phora artifrons* | Turku | 60.432 | 22.162 | 22-Jun-2020 | MaT |
| ZMUO.069792 | BOLD:ADM5152 | *Triphleba antricola* | Turku | 60.432 | 22.162 | 22-Jun-2020 | MaT |
| ZMUO.069839 | BOLD:ABA7002 | *Phora tincta* | Turku | 60.432 | 22.162 | 22-Jun-2020 | MaT |

| | | | | | | | |
|---|---|---|---|---|---|---|---|
| ZMUO.069860 | BOLD:ABA7002 | *Phora tincta* | Turku | 60.432 | 22.162 | 22-Jun-2020 | MaT |
| ZMUO.069900 | BOLD:ABA7002 | *Phora tincta* | Turku | 60.432 | 22.162 | 14-Jul-2020 | MaT |
| ZMUO.069914 | BOLD:ADM5152 | *Triphleba antricola* | Turku | 60.432 | 22.162 | 14-Jul-2020 | MaT |
| ZMUO.069952 | BOLD:ADM5152 | *Triphleba antricola* | Turku | 60.432 | 22.162 | 14-Jul-2020 | MaT |
| ZMUO.069978 | BOLD:ADM5152 | *Triphleba antricola* | Turku | 60.432 | 22.162 | 14-Jul-2020 | MaT |
| ZMUO.069990 | BOLD:ADM5152 | *Triphleba antricola* | Turku | 60.432 | 22.162 | 14-Jul-2020 | MaT |
| ZMUO.070003 | BOLD:AAP9864 | *Triphleba subcompleta* | Turku | 60.432 | 22.162 | 14-Jul-2020 | MaT |
| ZMUO.070022 | BOLD:ADM5152 | *Triphleba antricola* | Turku | 60.432 | 22.162 | 14-Jul-2020 | MaT |
| ZMUO.070028 | BOLD:AAP9864 | *Triphleba subcompleta* | Turku | 60.432 | 22.162 | 14-Jul-2020 | MaT |
| ZMUO.070032 | BOLD:ACQ9655 | *Anevrina unispinosa* | Turku | 60.432 | 22.162 | 14-Jul-2020 | MaT |
| ZMUO.070041 | BOLD:ACB3401 | *Chaetopleurophora erythronota* | Turku | 60.432 | 22.162 | 14-Jul-2020 | MaT |
| ZMUO.070044 | BOLD:ADM5152 | *Triphleba antricola* | Turku | 60.432 | 22.162 | 14-Jul-2020 | MaT |
| ZMUO.070046 | BOLD:ADM5152 | *Triphleba antricola* | Turku | 60.432 | 22.162 | 14-Jul-2020 | MaT |
| ZMUO.070050 | BOLD:ADM5152 | *Triphleba antricola* | Turku | 60.432 | 22.162 | 14-Jul-2020 | MaT |
| ZMUO.070056 | BOLD:ACK2513 | *Spiniphora excisa* | Turku | 60.432 | 22.162 | 14-Jul-2020 | MaT |
| ZMUO.070077 | BOLD:ADM5152 | *Triphleba antricola* | Turku | 60.432 | 22.162 | 14-Jul-2020 | MaT |
| ZMUO.070086 | BOLD:ADM5152 | *Triphleba antricola* | Turku | 60.432 | 22.162 | 14-Jul-2020 | MaT |
| ZMUO.070101 | BOLD:ADM5152 | *Triphleba antricola* | Turku | 60.432 | 22.162 | 04-Aug-2020 | MaT |
| ZMUO.070103 | BOLD:ACB3412 | *Triphleba nudipalpis* | Turku | 60.432 | 22.162 | 04-Aug-2020 | MaT |
| ZMUO.070115 | BOLD:ACK2513 | *Spiniphora excisa* | Turku | 60.432 | 22.162 | 04-Aug-2020 | MaT |
| ZMUO.070118 | BOLD:AAP9864 | *Triphleba subcompleta* | Turku | 60.432 | 22.162 | 04-Aug-2020 | MaT |
| ZMUO.070128 | BOLD:ADM5152 | *Triphleba antricola* | Turku | 60.432 | 22.162 | 04-Aug-2020 | MaT |
| ZMUO.070145 | BOLD:ADM5152 | *Triphleba antricola* | Turku | 60.432 | 22.162 | 04-Aug-2020 | MaT |
| ZMUO.070146 | BOLD:ADM5152 | *Triphleba antricola* | Turku | 60.432 | 22.162 | 04-Aug-2020 | MaT |
| ZMUO.070156 | BOLD:ACO9836 | *Triphleba distinguenda* | Turku | 60.432 | 22.162 | 04-Aug-2020 | MaT |
| ZMUO.070173 | BOLD:ADM5152 | *Triphleba antricola* | Turku | 60.432 | 22.162 | 04-Aug-2020 | MaT |
| ZMUO.070180 | BOLD:ADM5152 | *Triphleba antricola* | Turku | 60.432 | 22.162 | 04-Aug-2020 | MaT |
| ZMUO.070187 | BOLD:ADM5152 | *Triphleba antricola* | Turku | 60.432 | 22.162 | 04-Aug-2020 | MaT |
| ZMUO.070188 | BOLD:ADM5152 | *Triphleba antricola* | Turku | 60.432 | 22.162 | 04-Aug-2020 | MaT |
| ZMUO.070190 | BOLD:AAG3236 | *Diplonevra nitidula* | Turku | 60.432 | 22.162 | 04-Aug-2020 | MaT |
| ZMUO.070202 | BOLD:ADM5152 | *Triphleba antricola* | Turku | 60.432 | 22.162 | 04-Aug-2020 | MaT |
| ZMUO.070218 | BOLD:ACO9836 | *Triphleba distinguenda* | Turku | 60.432 | 22.162 | 04-Aug-2020 | MaT |
| ZMUO.070223 | BOLD:AAG3236 | *Diplonevra nitidula* | Turku | 60.432 | 22.162 | 04-Aug-2020 | MaT |
| ZMUO.070227 | BOLD:ABA7002 | *Phora tincta* | Turku | 60.432 | 22.162 | 04-Aug-2020 | MaT |
| ZMUO.070230 | BOLD:ADM5152 | *Triphleba antricola* | Turku | 60.432 | 22.162 | 04-Aug-2020 | MaT |
| ZMUO.070231 | BOLD:ADM5152 | *Triphleba antricola* | Turku | 60.432 | 22.162 | 04-Aug-2020 | MaT |
| ZMUO.070285 | BOLD:ACB3701 | *Diplonevra concinna* | Turku | 60.432 | 22.162 | 23-Aug-2020 | MaT |
| ZMUO.070294 | BOLD:ACN6664 | *Phora pubipes* | Turku | 60.432 | 22.162 | 23-Aug-2020 | MaT |
| ZMUO.070298 | BOLD:ADM5152 | *Triphleba antricola* | Turku | 60.432 | 22.162 | 23-Aug-2020 | MaT |
| ZMUO.070299 | BOLD:ACB3701 | *Diplonevra concinna* | Turku | 60.432 | 22.162 | 23-Aug-2020 | MaT |
| ZMUO.070304 | BOLD:ADA0453 | *Triphleba citreiformis* | Turku | 60.432 | 22.162 | 23-Aug-2020 | MaT |
| ZMUO.070308 | BOLD:ADM5152 | *Triphleba antricola* | Turku | 60.432 | 22.162 | 23-Aug-2020 | MaT |
| ZMUO.070311 | BOLD:ACD2464 | *Borophaga carinifrons* | Turku | 60.432 | 22.162 | 23-Aug-2020 | MaT |
| ZMUO.070316 | BOLD:ACB3701 | *Diplonevra concinna* | Turku | 60.432 | 22.162 | 23-Aug-2020 | MaT |
| ZMUO.070329 | BOLD:ACB3701 | *Diplonevra concinna* | Turku | 60.432 | 22.162 | 23-Aug-2020 | MaT |
| ZMUO.070335 | BOLD:ADM5152 | *Triphleba antricola* | Turku | 60.432 | 22.162 | 23-Aug-2020 | MaT |
| ZMUO.070343 | BOLD:AAG3236 | *Diplonevra nitidula* | Turku | 60.432 | 22.162 | 23-Aug-2020 | MaT |
| ZMUO.070361 | BOLD:ADM5152 | *Triphleba antricola* | Turku | 60.432 | 22.162 | 23-Aug-2020 | MaT |

| | | | | | | | |
|---|---|---|---|---|---|---|---|
| ZMUO.070365 | BOLD:ACB3412 | *Triphleba nudipalpis* | Turku | 60.432 | 22.162 | 23-Aug-2020 | MaT |
| ZMUO.070366 | BOLD:AAG3236 | *Diplonevra nitidula* | Turku | 60.432 | 22.162 | 23-Aug-2020 | MaT |
| ZMUO.070443 | BOLD:ACD2444 | *Hypocera mordellaria* | Kuhmo | 64.168 | 30.347 | 24-Jun-2020 | MaT |
| ZMUO.070444 | BOLD:ABA7001 | *Diplonevra glabra* | Kuhmo | 64.168 | 30.347 | 24-Jun-2020 | MaT |
| ZMUO.070456 | BOLD:AAG3249 | *Phora obscura* | Kuhmo | 64.168 | 30.347 | 24-Jun-2020 | MaT |
| ZMUO.070460 | BOLD:ACD2261 | *Phora artifrons* | Kuhmo | 64.168 | 30.347 | 24-Jun-2020 | MaT |
| ZMUO.070477 | BOLD:ACD2261 | *Phora artifrons* | Kuhmo | 64.168 | 30.347 | 15-Jul-2020 | MaT |
| ZMUO.070493 | BOLD:ABA7001 | *Diplonevra glabra* | Kuhmo | 64.168 | 30.347 | 15-Jul-2020 | MaT |
| ZMUO.070497 | BOLD:AFE7151 | *Anevrina thoracica* | Kuhmo | 64.168 | 30.347 | 15-Jul-2020 | MaT |
| ZMUO.070568 | BOLD:ADM5152 | *Triphleba antricola* | Kuhmo | 64.168 | 30.347 | 15-Aug-2020 | MaT |
| ZMUO.070576 | BOLD:ACD2464 | *Borophaga carinifrons* | Kuhmo | 64.168 | 30.347 | 15-Aug-2020 | MaT |
| ZMUO.070577 | BOLD:ADM5152 | *Triphleba antricola* | Kuhmo | 64.168 | 30.347 | 15-Aug-2020 | MaT |
| ZMUO.070596 | BOLD:AAG3236 | *Diplonevra nitidula* | Kuhmo | 64.168 | 30.347 | 15-Aug-2020 | MaT |
| ZMUO.070600 | BOLD:AAG3236 | *Diplonevra nitidula* | Kuhmo | 64.168 | 30.347 | 15-Aug-2020 | MaT |
| ZMUO.070606 | BOLD:ADM5152 | *Triphleba antricola* | Kuhmo | 64.168 | 30.347 | 15-Aug-2020 | MaT |
| ZMUO.070608 | BOLD:ADM5152 | *Triphleba antricola* | Kuhmo | 64.168 | 30.347 | 15-Aug-2020 | MaT |
| ZMUO.070623 | BOLD:ACD2464 | *Borophaga carinifrons* | Kuhmo | 64.168 | 30.347 | 15-Aug-2020 | MaT |
| ZMUO.070645 | BOLD:ADM5152 | *Triphleba antricola* | Kuhmo | 64.168 | 30.347 | 15-Aug-2020 | MaT |
| ZMUO.070647 | BOLD:AAG3236 | *Diplonevra nitidula* | Kuhmo | 64.168 | 30.347 | 15-Aug-2020 | MaT |
| ZMUO.070648 | BOLD:ACD2464 | *Borophaga carinifrons* | Kuhmo | 64.168 | 30.347 | 15-Aug-2020 | MaT |
| ZMUO.070656 | BOLD:ADM5152 | *Triphleba antricola* | Kuhmo | 64.168 | 30.347 | 15-Aug-2020 | MaT |
| ZMUO.070662 | BOLD:AAG3236 | *Diplonevra nitidula* | Kuhmo | 64.168 | 30.347 | 24-Aug-2020 | MaT |
| ZMUO.070681 | BOLD:ACD2464 | *Borophaga carinifrons* | Kuhmo | 64.168 | 30.347 | 24-Aug-2020 | MaT |
| ZMUO.070690 | BOLD:ADM5152 | *Triphleba antricola* | Kuhmo | 64.168 | 30.347 | 24-Aug-2020 | MaT |
| ZMUO.070691 | BOLD:AAG3236 | *Diplonevra nitidula* | Kuhmo | 64.168 | 30.347 | 24-Aug-2020 | MaT |
| ZMUO.070697 | BOLD:AAG3236 | *Diplonevra nitidula* | Kuhmo | 64.168 | 30.347 | 24-Aug-2020 | MaT |
| ZMUO.070705 | BOLD:ACD2464 | *Borophaga carinifrons* | Kuhmo | 64.168 | 30.347 | 24-Aug-2020 | MaT |
| ZMUO.070709 | BOLD:ACD2464 | *Borophaga carinifrons* | Kuhmo | 64.168 | 30.347 | 24-Aug-2020 | MaT |
| ZMUO.070713 | BOLD:ACD2464 | *Borophaga carinifrons* | Kuhmo | 64.168 | 30.347 | 24-Aug-2020 | MaT |
| ZMUO.070717 | BOLD:ADM5152 | *Triphleba antricola* | Kuhmo | 64.168 | 30.347 | 24-Aug-2020 | MaT |
| ZMUO.070718 | BOLD:ACD2464 | *Borophaga carinifrons* | Kuhmo | 64.168 | 30.347 | 24-Aug-2020 | MaT |
| ZMUO.070736 | BOLD:ACD2464 | *Borophaga carinifrons* | Kuhmo | 64.168 | 30.347 | 24-Aug-2020 | MaT |
| ZMUO.070790 | BOLD:ACF6067 | *Phora dubia* | Kuhmo | 64.166 | 30.309 | 27-Jun-2022 | MaT |
| ZMUO.070809 | BOLD:ACD2261 | *Phora artifrons* | Kuhmo | 64.166 | 30.309 | 27-Jun-2022 | MaT |
| ZMUO.070818 | BOLD:AFE4578 | *Anevrina thoracica* | Kuhmo | 64.166 | 30.309 | 27-Jun-2022 | MaT |
| ZMUO.070826 | BOLD:ACF6067 | *Phora dubia* | Kuhmo | 64.166 | 30.309 | 27-Jun-2022 | MaT |
| ZMUO.070921 | BOLD:ACE1464 | *Phalacrotophora fasciata* | Kuhmo | 64.166 | 30.309 | 29-Jul-2022 | MaT |
| ZMUO.070982 | BOLD:ACN6664 | *Phora pubipes* | Kuhmo | 64.166 | 30.309 | 26-Aug-2022 | MaT |
| ZMUO.071184 | BOLD:AAN8685 | *Conicera dauci* | Oulu | 65.137 | 25.885 | 09-Jul-2015 | MaT |
| ZMUO.071206 | BOLD:ADM5152 | *Triphleba antricola* | Oulu | 65.137 | 25.885 | 09-Jul-2015 | MaT |
| ZMUO.071213 | BOLD:ABA7002 | *Phora tincta* | Oulu | 65.137 | 25.885 | 09-Jul-2015 | MaT |
| ZMUO.071243 | BOLD:ACD2463 | *Borophaga agilis* | Oulu | 65.137 | 25.885 | 15-Jul-2015 | MaT |
| ZMUO.071283 | BOLD:ACD2463 | *Borophaga agilis* | Oulu | 65.137 | 25.885 | 15-Jul-2015 | MaT |
| ZMUO.071290 | BOLD:ACD2463 | *Borophaga agilis* | Oulu | 65.137 | 25.885 | 15-Jul-2015 | MaT |
| ZMUO.071297 | BOLD:ACD2463 | *Borophaga agilis* | Oulu | 65.137 | 25.885 | 15-Jul-2015 | MaT |
| ZMUO.071310 | BOLD:ACD2463 | *Borophaga agilis* | Oulu | 65.137 | 25.885 | 15-Jul-2015 | MaT |
| ZMUO.071343 | BOLD:ACD2464 | *Borophaga carinifrons* | Oulu | 65.137 | 25.885 | 18-Aug-2015 | MaT |
| ZMUO.071361 | BOLD:ACD2464 | *Borophaga carinifrons* | Oulu | 65.137 | 25.885 | 18-Aug-2015 | MaT |

| | | | | | | | |
|---|---|---|---|---|---|---|---|
| ZMUO.071374 | BOLD:ACD2463 | *Borophaga agilis* | Oulu | 65.137 | 25.885 | 18-Aug-2015 | MaT |
| ZMUO.071375 | BOLD:ACD2463 | *Borophaga agilis* | Oulu | 65.137 | 25.885 | 18-Aug-2015 | MaT |
| ZMUO.071376 | BOLD:ACD2464 | *Borophaga carinifrons* | Oulu | 65.137 | 25.885 | 18-Aug-2015 | MaT |
| ZMUO.071380 | BOLD:ACD2463 | *Borophaga agilis* | Oulu | 65.137 | 25.885 | 18-Aug-2015 | MaT |
| ZMUO.071383 | BOLD:ACD2464 | *Borophaga carinifrons* | Oulu | 65.137 | 25.885 | 18-Aug-2015 | MaT |
| ZMUO.071386 | BOLD:ACI8210 | *Phora convergens* | Oulu | 65.137 | 25.885 | 18-Aug-2015 | MaT |
| ZMUO.071388 | BOLD:ACN6664 | *Phora pubipes* | Oulu | 65.137 | 25.885 | 18-Aug-2015 | MaT |
| ZMUO.071390 | BOLD:ACD2463 | *Borophaga agilis* | Oulu | 65.137 | 25.885 | 18-Aug-2015 | MaT |
| ZMUO.071407 | BOLD:ACD2464 | *Borophaga carinifrons* | Oulu | 65.137 | 25.885 | 18-Aug-2015 | MaT |
| ZMUO.071413 | BOLD:ACD2464 | *Borophaga carinifrons* | Oulu | 65.137 | 25.885 | 18-Aug-2015 | MaT |
| ZMUO.071463 | BOLD:AAG3348 | *Triphleba aequalis* | Oulu | 65.137 | 25.851 | 29-Sep-2015 | MaT |
| ZMUO.071485 | BOLD:ACF0365 | *Triphleba bicornuta* | Oulu | 65.137 | 25.851 | 29-Sep-2015 | MaT |
| ZMUO.071489 | BOLD:AAG3348 | *Triphleba aequalis* | Oulu | 65.137 | 25.851 | 29-Sep-2015 | MaT |
| ZMUO.071490 | BOLD:ACF0365 | *Triphleba bicornuta* | Oulu | 65.137 | 25.851 | 29-Sep-2015 | MaT |
| ZMUO.071549 | BOLD:AAG3348 | *Triphleba aequalis* | Oulu | 65.137 | 25.851 | 30-Oct-2015 | MaT |
| ZMUO.071583 | BOLD:ACF0365 | *Triphleba bicornuta* | Oulu | 65.137 | 25.851 | 30-Oct-2015 | MaT |
| ZMUO.071588 | BOLD:ACF0365 | *Triphleba bicornuta* | Oulu | 65.137 | 25.851 | 30-Oct-2015 | MaT |
| ZMUO.071594 | BOLD:AAG3348 | *Triphleba aequalis* | Oulu | 65.137 | 25.851 | 30-Oct-2015 | MaT |
| ZMUO.071637 | BOLD:AFE8790 | *Plectanocnema nudipes* | Oulu | 65.063 | 25.469 | 25-May-2016 | MaT |
| ZMUO.071741 | BOLD:ADA0451 | *Borophaga femorata* | Oulu | 65.063 | 25.469 | 06-Jun-2016 | MaT |
| ZMUO.071782 | BOLD:AAP3611 | *Borophaga subsultans* | Oulu | 65.063 | 25.469 | 06-Jun-2016 | MaT |
| ZMUO.071948 | BOLD:ACD2464 | *Borophaga carinifrons* | Oulu | 65.063 | 25.469 | 08-Aug-2016 | MaT |
| ZMUO.071964 | BOLD:ACD2463 | *Borophaga agilis* | Oulu | 65.063 | 25.469 | 08-Aug-2016 | MaT |
| ZMUO.071977 | BOLD:ACD2463 | *Borophaga agilis* | Oulu | 65.063 | 25.469 | 08-Aug-2016 | MaT |
| ZMUO.071978 | BOLD:ACD2463 | *Borophaga agilis* | Oulu | 65.063 | 25.469 | 08-Aug-2016 | MaT |
| ZMUO.072123 | BOLD:ACE3722 | *Diplonevra florescens* | Kitee | 62.078 | 30.197 | 18-May-2017 | MaT |
| ZMUO.072264 | BOLD:ACD2444 | *Hypocera mordellaria* | Kitee | 62.078 | 30.197 | 05-Jul-2017 | MaT |
| ZMUO.072369 | BOLD:AFE7151 | *Anevrina thoracica* | Kitee | 62.076 | 30.18 | 23-May-2016 | MaT |
| ZMUO.072376 | BOLD:ABA7001 | *Diplonevra glabra* | Kitee | 62.076 | 30.18 | 23-May-2016 | MaT |
| ZMUO.072378 | BOLD:ABA7001 | *Diplonevra glabra* | Kitee | 62.076 | 30.18 | 23-May-2016 | MaT |
| ZMUO.072385 | BOLD:ABA7001 | *Diplonevra glabra* | Kitee | 62.076 | 30.18 | 23-May-2016 | MaT |
| ZMUO.072394 | BOLD:ACF6067 | *Phora dubia* | Kitee | 62.076 | 30.18 | 06-Jun-2016 | MaT |
| ZMUO.072395 | BOLD:ABA7001 | *Diplonevra glabra* | Kitee | 62.076 | 30.18 | 06-Jun-2016 | MaT |
| ZMUO.072398 | BOLD:ABA7001 | *Diplonevra glabra* | Kitee | 62.076 | 30.18 | 06-Jun-2016 | MaT |
| ZMUO.072406 | BOLD:ABA7001 | *Diplonevra glabra* | Kitee | 62.076 | 30.18 | 06-Jun-2016 | MaT |
| ZMUO.072407 | BOLD:ABA7001 | *Diplonevra glabra* | Kitee | 62.076 | 30.18 | 06-Jun-2016 | MaT |
| ZMUO.072411 | BOLD:ABA7001 | *Diplonevra glabra* | Kitee | 62.076 | 30.18 | 06-Jun-2016 | MaT |
| ZMUO.072420 | BOLD:ABA7001 | *Diplonevra glabra* | Kitee | 62.076 | 30.18 | 06-Jun-2016 | MaT |
| ZMUO.072421 | BOLD:ABA7001 | *Diplonevra glabra* | Kitee | 62.076 | 30.18 | 06-Jun-2016 | MaT |
| ZMUO.072428 | BOLD:ABA7001 | *Diplonevra glabra* | Kitee | 62.076 | 30.18 | 06-Jun-2016 | MaT |
| ZMUO.072440 | BOLD:ABA7001 | *Diplonevra glabra* | Kitee | 62.076 | 30.18 | 06-Jun-2016 | MaT |
| ZMUO.072442 | BOLD:ABA7001 | *Diplonevra glabra* | Kitee | 62.076 | 30.18 | 06-Jun-2016 | MaT |
| ZMUO.072443 | BOLD:ACF6067 | *Phora dubia* | Kitee | 62.076 | 30.18 | 06-Jun-2016 | MaT |
| ZMUO.072455 | BOLD:ABA7001 | *Diplonevra glabra* | Kitee | 62.076 | 30.18 | 06-Jun-2016 | MaT |
| ZMUO.072456 | BOLD:ABA7001 | *Diplonevra glabra* | Kitee | 62.076 | 30.18 | 06-Jun-2016 | MaT |
| ZMUO.072479 | BOLD:ACE2645 | *Conicera floricola* | Kitee | 62.076 | 30.18 | 21-Jul-2016 | MaT |
| ZMUO.072557 | BOLD:ACN6664 | *Phora pubipes* | Kitee | 62.076 | 30.18 | 01-Aug-2016 | MaT |
| ZMUO.072662 | BOLD:AAP3611 | *Borophaga subsultans* | Kitee | 62.088 | 30.223 | 15-Jun-2017 | MaT |

| | | | | | | | |
|---|---|---|---|---|---|---|---|
| ZMUO.072665 | BOLD:ABA7002 | *Phora tincta* | Kitee | 62.088 | 30.223 | 15-Jun-2017 | MaT |
| ZMUO.072669 | BOLD:ACF6067 | *Phora dubia* | Kitee | 62.088 | 30.223 | 24-Jun-2017 | MaT |
| ZMUO.072694 | BOLD:ABA7001 | *Diplonevra glabra* | Kitee | 62.088 | 30.223 | 24-Jun-2017 | MaT |
| ZMUO.072715 | BOLD:ADM5152 | *Triphleba antricola* | Kitee | 62.088 | 30.223 | 05-Jul-2017 | MaT |
| ZMUO.072745 | BOLD:ACE2645 | *Conicera floricola* | Kitee | 62.088 | 30.223 | 16-Jul-2017 | MaT |
| ZMUO.072790 | BOLD:ADM5152 | *Triphleba antricola* | Kitee | 62.088 | 30.223 | 24-Jul-2017 | MaT |
| ZMUO.072826 | BOLD:ADM5152 | *Triphleba antricola* | Kitee | 62.088 | 30.223 | 02-Aug-2017 | MaT |
| ZMUO.073041 | BOLD:ACQ9655 | *Anevrina unispinosa* | Oulu | 65.059 | 25.52 | 12-Jul-2022 | MaT |
| ZMUO.073048 | BOLD:AAP3611 | *Borophaga subsultans* | Oulu | 65.059 | 25.52 | 12-Jul-2022 | MaT |
| ZMUO.073261 | BOLD:ABA7002 | *Phora tincta* | Oulu | 65.059 | 25.52 | 02-Aug-2022 | MaT |
| ZMUO.073364 | BOLD:AFE7151 | *Anevrina thoracica* | Oulu | 65.059 | 25.52 | 02-Aug-2022 | MaT |
| ZMUO.073398 | BOLD:AEB8779 | *Phora convergens* | Oulu | 65.059 | 25.52 | 15-Aug-2022 | MaT |
| ZMUO.073471 | BOLD:AEH1787 | *Phora occidentata* | Oulu | 65.059 | 25.52 | 30-Aug-2022 | MaT |
| ZMUO.073538 | BOLD:ABA7001 | *Diplonevra glabra* | Laukaa | 62.223 | 26.099 | 21-Jun-2021 | MaT |
| ZMUO.073542 | BOLD:ACD2444 | *Hypocera mordellaria* | Laukaa | 62.223 | 26.099 | 21-Jun-2021 | MaT |
| ZMUO.073544 | BOLD:ABA7001 | *Diplonevra glabra* | Laukaa | 62.223 | 26.099 | 21-Jun-2021 | MaT |
| ZMUO.073564 | BOLD:ACF8062 | *Triphleba papillata* | Laukaa | 62.223 | 26.099 | 21-Jun-2021 | MaT |
| ZMUO.073580 | BOLD:ADM5152 | *Triphleba antricola* | Laukaa | 62.223 | 26.099 | 21-Jun-2021 | MaT |
| ZMUO.073590 | BOLD:ABA7001 | *Diplonevra glabra* | Laukaa | 62.223 | 26.099 | 21-Jun-2021 | MaT |
| ZMUO.073592 | BOLD:ABA7001 | *Diplonevra glabra* | Laukaa | 62.223 | 26.099 | 21-Jun-2021 | MaT |
| ZMUO.073602 | BOLD:ABA7001 | *Diplonevra glabra* | Laukaa | 62.223 | 26.099 | 21-Jun-2021 | MaT |
| ZMUO.073639 | BOLD:AAG3236 | *Diplonevra nitidula* | Laukaa | 62.223 | 26.099 | 21-Jun-2021 | MaT |
| ZMUO.073645 | BOLD:ACB3394 | *Phora bullata* | Laukaa | 62.223 | 26.099 | 21-Jun-2021 | MaT |
| ZMUO.073712 | BOLD:AAG3236 | *Diplonevra nitidula* | Laukaa | 62.223 | 26.099 | 11-Jul-2021 | MaT |
| ZMUO.073719 | BOLD:ACE1464 | *Phalacrotophora fasciata* | Laukaa | 62.223 | 26.099 | 11-Jul-2021 | MaT |
| ZMUO.073723 | BOLD:ADM5152 | *Triphleba antricola* | Laukaa | 62.223 | 26.099 | 11-Jul-2021 | MaT |
| ZMUO.073724 | BOLD:ADM5152 | *Triphleba antricola* | Laukaa | 62.223 | 26.099 | 11-Jul-2021 | MaT |
| ZMUO.073736 | BOLD:ACE1464 | *Phalacrotophora fasciata* | Laukaa | 62.223 | 26.099 | 11-Jul-2021 | MaT |
| ZMUO.073768 | BOLD:ABW2086 | *Pseudacteon fennicus* | Laukaa | 62.223 | 26.099 | 11-Jul-2021 | MaT |
| ZMUO.073771 | BOLD:ACE2645 | *Conicera floricola* | Laukaa | 62.223 | 26.099 | 11-Jul-2021 | MaT |
| ZMUO.073792 | BOLD:ACE1464 | *Phalacrotophora fasciata* | Laukaa | 62.223 | 26.099 | 11-Jul-2021 | MaT |
| ZMUO.073803 | BOLD:ACF4704 | *Metopina oligoneura* | Laukaa | 62.223 | 26.099 | 11-Jul-2021 | MaT |
| ZMUO.073814 | BOLD:ACR0978 | *Pseudacteon formicarum* | Laukaa | 62.223 | 26.099 | 11-Jul-2021 | MaT |
| ZMUO.073825 | BOLD:ADM5152 | *Triphleba antricola* | Laukaa | 62.223 | 26.099 | 11-Jul-2021 | MaT |
| ZMUO.073836 | BOLD:ACE1464 | *Phalacrotophora fasciata* | Laukaa | 62.223 | 26.099 | 11-Jul-2021 | MaT |
| ZMUO.073849 | BOLD:ACQ3746 | *Gymnophora arcuata* | Laukaa | 62.223 | 26.099 | 11-Jul-2021 | MaT |
| ZMUO.073851 | BOLD:AAG3236 | *Diplonevra nitidula* | Laukaa | 62.223 | 26.099 | 11-Jul-2021 | MaT |
| ZMUO.073854 | BOLD:AAG3236 | *Diplonevra nitidula* | Laukaa | 62.223 | 26.099 | 11-Jul-2021 | MaT |
| ZMUO.073857 | BOLD:AAG3236 | *Diplonevra nitidula* | Laukaa | 62.223 | 26.099 | 11-Jul-2021 | MaT |
| ZMUO.073862 | BOLD:AAG3236 | *Diplonevra nitidula* | Laukaa | 62.223 | 26.099 | 11-Jul-2021 | MaT |
| ZMUO.073887 | BOLD:ADM5152 | *Triphleba antricola* | Laukaa | 62.223 | 26.099 | 11-Jul-2021 | MaT |
| ZMUO.073890 | BOLD:ACE1464 | *Phalacrotophora fasciata* | Laukaa | 62.223 | 26.099 | 11-Jul-2021 | MaT |
| ZMUO.073899 | BOLD:ACF4704 | *Metopina oligoneura* | Laukaa | 62.223 | 26.099 | 11-Jul-2021 | MaT |
| ZMUO.073914 | BOLD:ADM5152 | *Triphleba antricola* | Laukaa | 62.223 | 26.099 | 11-Jul-2021 | MaT |
| ZMUO.073923 | BOLD:ADM5152 | *Triphleba antricola* | Laukaa | 62.223 | 26.099 | 11-Jul-2021 | MaT |

| | | | | | | | |
|---|---|---|---|---|---|---|---|
| ZMUO.073933 | BOLD:AAG3236 | *Diplonevra nitidula* | Laukaa | 62.223 | 26.099 | 11-Jul-2021 | MaT |
| ZMUO.073941 | BOLD:ADM5152 | *Triphleba antricola* | Laukaa | 62.223 | 26.099 | 11-Jul-2021 | MaT |
| ZMUO.073944 | BOLD:ACJ1099 | *Conicera schnittmanni* | Laukaa | 62.223 | 26.099 | 20-Jul-2021 | MaT |
| ZMUO.073949 | BOLD:ADM5152 | *Triphleba antricola* | Laukaa | 62.223 | 26.099 | 20-Jul-2021 | MaT |
| ZMUO.073952 | BOLD:ACD2464 | *Borophaga carinifrons* | Laukaa | 62.223 | 26.099 | 20-Jul-2021 | MaT |
| ZMUO.073953 | BOLD:ACF4704 | *Metopina oligoneura* | Laukaa | 62.223 | 26.099 | 20-Jul-2021 | MaT |
| ZMUO.073964 | BOLD:ACO9836 | *Triphleba distinguenda* | Laukaa | 62.223 | 26.099 | 20-Jul-2021 | MaT |
| ZMUO.073970 | BOLD:ACO7758 | *Pseudacteon brevicauda* | Laukaa | 62.223 | 26.099 | 20-Jul-2021 | MaT |
| ZMUO.073976 | BOLD:ADM5152 | *Triphleba antricola* | Laukaa | 62.223 | 26.099 | 20-Jul-2021 | MaT |
| ZMUO.073983 | BOLD:ACZ8391 | *Diplonevra pilosella* | Laukaa | 62.223 | 26.099 | 20-Jul-2021 | MaT |
| ZMUO.074007 | BOLD:ADM5152 | *Triphleba antricola* | Laukaa | 62.223 | 26.099 | 20-Jul-2021 | MaT |
| ZMUO.074016 | BOLD:ADM5152 | *Triphleba antricola* | Laukaa | 62.223 | 26.099 | 20-Jul-2021 | MaT |
| ZMUO.074022 | BOLD:ACD2464 | *Borophaga carinifrons* | Laukaa | 62.223 | 26.099 | 20-Jul-2021 | MaT |
| ZMUO.074023 | BOLD:ADM5152 | *Triphleba antricola* | Laukaa | 62.223 | 26.099 | 20-Jul-2021 | MaT |
| ZMUO.074042 | BOLD:ABA7001 | *Diplonevra glabra* | Vartsila | 62.192 | 30.627 | 21-Jun-2021 | MaT |
| ZMUO.074050 | BOLD:AAG3236 | *Diplonevra nitidula* | Vartsila | 62.192 | 30.627 | 21-Jun-2021 | MaT |
| ZMUO.074056 | BOLD:AAG3236 | *Diplonevra nitidula* | Vartsila | 62.192 | 30.627 | 21-Jun-2021 | MaT |
| ZMUO.074060 | BOLD:ABA7001 | *Diplonevra glabra* | Vartsila | 62.192 | 30.627 | 21-Jun-2021 | MaT |
| ZMUO.074064 | BOLD:ABA7001 | *Diplonevra glabra* | Vartsila | 62.192 | 30.627 | 21-Jun-2021 | MaT |
| ZMUO.074089 | BOLD:ABA7001 | *Diplonevra glabra* | Vartsila | 62.192 | 30.627 | 21-Jun-2021 | MaT |
| ZMUO.074095 | BOLD:ABA7001 | *Diplonevra glabra* | Vartsila | 62.192 | 30.627 | 21-Jun-2021 | MaT |
| ZMUO.074096 | BOLD:ABA7001 | *Diplonevra glabra* | Vartsila | 62.192 | 30.627 | 21-Jun-2021 | MaT |
| ZMUO.074098 | BOLD:ABA7001 | *Diplonevra glabra* | Vartsila | 62.192 | 30.627 | 21-Jun-2021 | MaT |
| ZMUO.074100 | BOLD:ABA7001 | *Diplonevra glabra* | Vartsila | 62.192 | 30.627 | 21-Jun-2021 | MaT |
| ZMUO.074102 | BOLD:AAG3236 | *Diplonevra nitidula* | Vartsila | 62.192 | 30.627 | 21-Jun-2021 | MaT |
| ZMUO.074124 | BOLD:AAG3236 | *Diplonevra nitidula* | Vartsila | 62.192 | 30.627 | 19-Jul-2021 | MaT |
| ZMUO.074130 | BOLD:ACD2463 | *Borophaga agilis* | Vartsila | 62.192 | 30.627 | 19-Jul-2021 | MaT |
| ZMUO.074133 | BOLD:AAG3236 | *Diplonevra nitidula* | Vartsila | 62.192 | 30.627 | 19-Jul-2021 | MaT |
| ZMUO.074134 | BOLD:AAG3236 | *Diplonevra nitidula* | Vartsila | 62.192 | 30.627 | 19-Jul-2021 | MaT |
| ZMUO.074137 | BOLD:ABA7001 | *Diplonevra glabra* | Vartsila | 62.192 | 30.627 | 19-Jul-2021 | MaT |
| ZMUO.074141 | BOLD:ACQ9655 | *Anevrina unispinosa* | Vartsila | 62.192 | 30.627 | 19-Jul-2021 | MaT |
| ZMUO.074150 | BOLD:AAG3236 | *Diplonevra nitidula* | Vartsila | 62.192 | 30.627 | 19-Jul-2021 | MaT |
| ZMUO.074164 | BOLD:ACB3701 | *Diplonevra concinna* | Vartsila | 62.192 | 30.627 | 19-Jul-2021 | MaT |
| ZMUO.074165 | BOLD:AFE7151 | *Anevrina thoracica* | Vartsila | 62.192 | 30.627 | 19-Jul-2021 | MaT |
| ZMUO.074167 | BOLD:AAG3236 | *Diplonevra nitidula* | Vartsila | 62.192 | 30.627 | 19-Jul-2021 | MaT |
| ZMUO.074205 | BOLD:ABA7001 | *Diplonevra glabra* | Vartsila | 62.192 | 30.627 | 19-Jul-2021 | MaT |
| ZMUO.074208 | BOLD:AAG3236 | *Diplonevra nitidula* | Vartsila | 62.192 | 30.627 | 19-Jul-2021 | MaT |
| ZMUO.074209 | BOLD:ACD2463 | *Borophaga agilis* | Vartsila | 62.192 | 30.627 | 19-Jul-2021 | MaT |
| ZMUO.074212 | BOLD:ACQ9655 | *Anevrina unispinosa* | Vartsila | 62.192 | 30.627 | 19-Jul-2021 | MaT |
| ZMUO.074213 | BOLD:AAG3236 | *Diplonevra nitidula* | Vartsila | 62.192 | 30.627 | 19-Jul-2021 | MaT |
| ZMUO.074214 | BOLD:AAG3236 | *Diplonevra nitidula* | Vartsila | 62.192 | 30.627 | 19-Jul-2021 | MaT |
| ZMUO.074215 | BOLD:ACQ9655 | *Anevrina unispinosa* | Vartsila | 62.192 | 30.627 | 19-Jul-2021 | MaT |
| ZMUO.074217 | BOLD:AAG3236 | *Diplonevra nitidula* | Vartsila | 62.192 | 30.627 | 19-Jul-2021 | MaT |
| ZMUO.074228 | BOLD:ABA7001 | *Diplonevra glabra* | Vartsila | 62.192 | 30.627 | 19-Jul-2021 | MaT |
| ZMUO.074229 | BOLD:AAG3236 | *Diplonevra nitidula* | Vartsila | 62.192 | 30.627 | 19-Jul-2021 | MaT |
| ZMUO.074231 | BOLD:AAG3236 | *Diplonevra nitidula* | Vartsila | 62.192 | 30.627 | 19-Jul-2021 | MaT |
| ZMUO.074232 | BOLD:AAG3236 | *Diplonevra nitidula* | Vartsila | 62.192 | 30.627 | 19-Jul-2021 | MaT |
| ZMUO.074234 | BOLD:AAG3236 | *Diplonevra nitidula* | Vartsila | 62.192 | 30.627 | 19-Jul-2021 | MaT |

| | | | | | | | |
|---|---|---|---|---|---|---|---|
| ZMUO.074241 | BOLD:AAG3236 | *Diplonevra nitidula* | Vartsila | 62.192 | 30.627 | 19-Jul-2021 | MaT |
| ZMUO.074242 | BOLD:AAG3236 | *Diplonevra nitidula* | Vartsila | 62.192 | 30.627 | 19-Jul-2021 | MaT |
| ZMUO.074243 | BOLD:ACD2463 | *Borophaga agilis* | Vartsila | 62.192 | 30.627 | 19-Jul-2021 | MaT |
| ZMUO.074248 | BOLD:AAG3236 | *Diplonevra nitidula* | Vartsila | 62.192 | 30.627 | 19-Jul-2021 | MaT |
| ZMUO.074257 | BOLD:AAG3236 | *Diplonevra nitidula* | Vartsila | 62.192 | 30.627 | 19-Jul-2021 | MaT |
| ZMUO.074281 | BOLD:AAG3236 | *Diplonevra nitidula* | Vartsila | 62.192 | 30.627 | 19-Jul-2021 | MaT |
| ZMUO.074282 | BOLD:AAG3236 | *Diplonevra nitidula* | Vartsila | 62.192 | 30.627 | 19-Jul-2021 | MaT |
| ZMUO.074288 | BOLD:AAG3236 | *Diplonevra nitidula* | Vartsila | 62.192 | 30.627 | 19-Jul-2021 | MaT |
| ZMUO.074294 | BOLD:AAG3236 | *Diplonevra nitidula* | Vartsila | 62.192 | 30.627 | 19-Jul-2021 | MaT |
| ZMUO.074298 | BOLD:ABA7001 | *Diplonevra glabra* | Vartsila | 62.192 | 30.627 | 19-Jul-2021 | MaT |
| ZMUO.074309 | BOLD:AAG3236 | *Diplonevra nitidula* | Vartsila | 62.192 | 30.627 | 19-Jul-2021 | MaT |
| ZMUO.074311 | BOLD:ACB3701 | *Diplonevra concinna* | Vartsila | 62.192 | 30.627 | 19-Jul-2021 | MaT |
| ZMUO.074319 | BOLD:ABA7001 | *Diplonevra glabra* | Vartsila | 62.192 | 30.627 | 19-Jul-2021 | MaT |
| ZMUO.074323 | BOLD:AAG3236 | *Diplonevra nitidula* | Vartsila | 62.192 | 30.627 | 19-Jul-2021 | MaT |
| ZMUO.074332 | BOLD:ACQ9655 | *Anevrina unispinosa* | Vartsila | 62.192 | 30.627 | 19-Jul-2021 | MaT |
| ZMUO.074336 | BOLD:AAG3236 | *Diplonevra nitidula* | Vartsila | 62.192 | 30.627 | 19-Jul-2021 | MaT |
| ZMUO.074344 | BOLD:ACD2463 | *Borophaga agilis* | Vartsila | 62.192 | 30.627 | 19-Jul-2021 | MaT |
| ZMUO.074347 | BOLD:ACD2463 | *Borophaga agilis* | Vartsila | 62.192 | 30.627 | 19-Jul-2021 | MaT |
| ZMUO.074354 | BOLD:AAG3236 | *Diplonevra nitidula* | Vartsila | 62.192 | 30.627 | 19-Jul-2021 | MaT |
| ZMUO.074356 | BOLD:ABA7001 | *Diplonevra glabra* | Vartsila | 62.192 | 30.627 | 19-Jul-2021 | MaT |
| ZMUO.074357 | BOLD:AAG3236 | *Diplonevra nitidula* | Vartsila | 62.192 | 30.627 | 19-Jul-2021 | MaT |
| ZMUO.074363 | BOLD:AAG3236 | *Diplonevra nitidula* | Vartsila | 62.192 | 30.627 | 19-Jul-2021 | MaT |
| ZMUO.074367 | BOLD:AEB8779 | *Phora convergens* | Vartsila | 62.192 | 30.627 | 19-Jul-2021 | MaT |
| ZMUO.074388 | BOLD:AAG3236 | *Diplonevra nitidula* | Vartsila | 62.192 | 30.627 | 19-Jul-2021 | MaT |
| ZMUO.074396 | BOLD:ACF4704 | *Metopina oligoneura* | Parikkala | 61.566 | 29.561 | 21-Jun-2021 | MaT |
| ZMUO.074403 | BOLD:ADM5152 | *Triphleba antricola* | Parikkala | 61.566 | 29.561 | 21-Jun-2021 | MaT |
| ZMUO.074404 | BOLD:ACB3412 | *Triphleba nudipalpis* | Parikkala | 61.566 | 29.561 | 21-Jun-2021 | MaT |
| ZMUO.074410 | BOLD:ACF8494 | *Conicera similis* | Parikkala | 61.566 | 29.561 | 21-Jun-2021 | MaT |
| ZMUO.074414 | BOLD:AAG3236 | *Diplonevra nitidula* | Parikkala | 61.566 | 29.561 | 21-Jun-2021 | MaT |
| ZMUO.074419 | BOLD:ACF4704 | *Metopina oligoneura* | Parikkala | 61.566 | 29.561 | 21-Jun-2021 | MaT |
| ZMUO.074422 | BOLD:AAG3236 | *Diplonevra nitidula* | Parikkala | 61.566 | 29.561 | 21-Jun-2021 | MaT |
| ZMUO.074425 | BOLD:ACF4704 | *Metopina oligoneura* | Parikkala | 61.566 | 29.561 | 21-Jun-2021 | MaT |
| ZMUO.074461 | BOLD:ACF4704 | *Metopina oligoneura* | Parikkala | 61.566 | 29.561 | 21-Jun-2021 | MaT |
| ZMUO.074466 | BOLD:ACF4704 | *Metopina oligoneura* | Parikkala | 61.566 | 29.561 | 21-Jun-2021 | MaT |
| ZMUO.074473 | BOLD:ACB3394 | *Phora bullata* | Parikkala | 61.566 | 29.561 | 21-Jun-2021 | MaT |
| ZMUO.074485 | BOLD:ACF4704 | *Metopina oligoneura* | Parikkala | 61.566 | 29.561 | 21-Jun-2021 | MaT |
| ZMUO.074505 | BOLD:ACE2645 | *Conicera floricola* | Parikkala | 61.566 | 29.561 | 21-Jun-2021 | MaT |
| ZMUO.074507 | BOLD:ACF8494 | *Conicera similis* | Parikkala | 61.566 | 29.561 | 21-Jun-2021 | MaT |
| ZMUO.074516 | BOLD:ACF4704 | *Metopina oligoneura* | Parikkala | 61.566 | 29.561 | 21-Jun-2021 | MaT |
| ZMUO.074518 | BOLD:ADM5152 | *Triphleba antricola* | Parikkala | 61.566 | 29.561 | 21-Jun-2021 | MaT |
| ZMUO.074535 | BOLD:ACF4704 | *Metopina oligoneura* | Parikkala | 61.566 | 29.561 | 21-Jun-2021 | MaT |
| ZMUO.074537 | BOLD:ACR0978 | *Pseudacteon formicarum* | Parikkala | 61.566 | 29.561 | 21-Jun-2021 | MaT |
| ZMUO.074545 | BOLD:ACF4704 | *Metopina oligoneura* | Parikkala | 61.566 | 29.561 | 21-Jun-2021 | MaT |
| ZMUO.074547 | BOLD:ACE2645 | *Conicera floricola* | Parikkala | 61.566 | 29.561 | 21-Jun-2021 | MaT |
| ZMUO.074549 | BOLD:ACF4704 | *Metopina oligoneura* | Parikkala | 61.566 | 29.561 | 21-Jun-2021 | MaT |
| ZMUO.074561 | BOLD:ACF4704 | *Metopina oligoneura* | Parikkala | 61.566 | 29.561 | 21-Jun-2021 | MaT |
| ZMUO.074562 | BOLD:ACF4704 | *Metopina oligoneura* | Parikkala | 61.566 | 29.561 | 21-Jun-2021 | MaT |
| ZMUO.074563 | BOLD:ACF4704 | *Metopina oligoneura* | Parikkala | 61.566 | 29.561 | 21-Jun-2021 | MaT |

| | | | | | | | |
|---|---|---|---|---|---|---|---|
| ZMUO.074567 | BOLD:ACF4704 | *Metopina oligoneura* | Parikkala | 61.566 | 29.561 | 21-Jun-2021 | MaT |
| ZMUO.074569 | BOLD:ACF4704 | *Metopina oligoneura* | Parikkala | 61.566 | 29.561 | 21-Jun-2021 | MaT |
| ZMUO.074571 | BOLD:ACR0978 | *Pseudacteon formicarum* | Parikkala | 61.566 | 29.561 | 21-Jun-2021 | MaT |
| ZMUO.074572 | BOLD:ACF4704 | *Metopina oligoneura* | Parikkala | 61.566 | 29.561 | 21-Jun-2021 | MaT |
| ZMUO.074576 | BOLD:ACF4704 | *Metopina oligoneura* | Parikkala | 61.566 | 29.561 | 21-Jun-2021 | MaT |
| ZMUO.074581 | BOLD:ACF4704 | *Metopina oligoneura* | Parikkala | 61.566 | 29.561 | 21-Jun-2021 | MaT |
| ZMUO.074586 | BOLD:ACF4704 | *Metopina oligoneura* | Parikkala | 61.566 | 29.561 | 21-Jun-2021 | MaT |
| ZMUO.074590 | BOLD:ACF4704 | *Metopina oligoneura* | Parikkala | 61.566 | 29.561 | 21-Jun-2021 | MaT |
| ZMUO.074595 | BOLD:ACF4704 | *Metopina oligoneura* | Parikkala | 61.566 | 29.561 | 21-Jun-2021 | MaT |
| ZMUO.074598 | BOLD:AAG3236 | *Diplonevra nitidula* | Parikkala | 61.566 | 29.561 | 21-Jun-2021 | MaT |
| ZMUO.074599 | BOLD:ACF4704 | *Metopina oligoneura* | Parikkala | 61.566 | 29.561 | 21-Jun-2021 | MaT |
| ZMUO.074608 | BOLD:ACF4704 | *Metopina oligoneura* | Parikkala | 61.566 | 29.561 | 21-Jun-2021 | MaT |
| ZMUO.074612 | BOLD:AAG3236 | *Diplonevra nitidula* | Parikkala | 61.566 | 29.561 | 21-Jun-2021 | MaT |
| ZMUO.074621 | BOLD:ACF4704 | *Metopina oligoneura* | Parikkala | 61.566 | 29.561 | 21-Jun-2021 | MaT |
| ZMUO.074624 | BOLD:ACF4704 | *Metopina oligoneura* | Parikkala | 61.566 | 29.561 | 21-Jun-2021 | MaT |
| ZMUO.074638 | BOLD:ACF4704 | *Metopina oligoneura* | Parikkala | 61.566 | 29.561 | 21-Jun-2021 | MaT |
| ZMUO.074639 | BOLD:ACF4704 | *Metopina oligoneura* | Parikkala | 61.566 | 29.561 | 21-Jun-2021 | MaT |
| ZMUO.074640 | BOLD:ACF4704 | *Metopina oligoneura* | Parikkala | 61.566 | 29.561 | 21-Jun-2021 | MaT |
| ZMUO.074648 | BOLD:ACF4704 | *Metopina oligoneura* | Parikkala | 61.566 | 29.561 | 21-Jun-2021 | MaT |
| ZMUO.074657 | BOLD:ADM5152 | *Triphleba antricola* | Parikkala | 61.566 | 29.561 | 21-Jun-2021 | MaT |
| ZMUO.074661 | BOLD:ACF4704 | *Metopina oligoneura* | Parikkala | 61.566 | 29.561 | 21-Jun-2021 | MaT |
| ZMUO.074666 | BOLD:ACP3737 | *Phora edentata* | Parikkala | 61.566 | 29.561 | 21-Jun-2021 | MaT |
| ZMUO.074674 | BOLD:ACF4704 | *Metopina oligoneura* | Parikkala | 61.566 | 29.561 | 21-Jun-2021 | MaT |
| ZMUO.074677 | BOLD:ACF4704 | *Metopina oligoneura* | Parikkala | 61.566 | 29.561 | 21-Jun-2021 | MaT |
| ZMUO.074680 | BOLD:ACF4704 | *Metopina oligoneura* | Parikkala | 61.566 | 29.561 | 21-Jun-2021 | MaT |
| ZMUO.074681 | BOLD:ACF4704 | *Metopina oligoneura* | Parikkala | 61.566 | 29.561 | 21-Jun-2021 | MaT |
| ZMUO.074682 | BOLD:ACF4704 | *Metopina oligoneura* | Parikkala | 61.566 | 29.561 | 21-Jun-2021 | MaT |
| ZMUO.074687 | BOLD:ACF4704 | *Metopina oligoneura* | Parikkala | 61.566 | 29.561 | 21-Jun-2021 | MaT |
| ZMUO.074702 | BOLD:ACF4704 | *Metopina oligoneura* | Parikkala | 61.566 | 29.561 | 21-Jun-2021 | MaT |
| ZMUO.074704 | BOLD:ACF4704 | *Metopina oligoneura* | Parikkala | 61.566 | 29.561 | 21-Jun-2021 | MaT |
| ZMUO.074708 | BOLD:ACF4704 | *Metopina oligoneura* | Parikkala | 61.566 | 29.561 | 21-Jun-2021 | MaT |
| ZMUO.074714 | BOLD:ADW4894 | *Pseudacteon lundbecki* | Parikkala | 61.566 | 29.561 | 21-Jun-2021 | MaT |
| ZMUO.074718 | BOLD:ACF4704 | *Metopina oligoneura* | Parikkala | 61.566 | 29.561 | 21-Jun-2021 | MaT |
| ZMUO.074720 | BOLD:ACF4704 | *Metopina oligoneura* | Parikkala | 61.566 | 29.561 | 21-Jun-2021 | MaT |
| ZMUO.074721 | BOLD:ABA7002 | *Phora tincta* | Parikkala | 61.566 | 29.561 | 21-Jun-2021 | MaT |
| ZMUO.074724 | BOLD:ACF4704 | *Metopina oligoneura* | Parikkala | 61.566 | 29.561 | 21-Jun-2021 | MaT |
| ZMUO.074727 | BOLD:ACF4704 | *Metopina oligoneura* | Parikkala | 61.566 | 29.561 | 21-Jun-2021 | MaT |
| ZMUO.074729 | BOLD:ACF4704 | *Metopina oligoneura* | Parikkala | 61.566 | 29.561 | 21-Jun-2021 | MaT |
| ZMUO.074731 | BOLD:ACF4704 | *Metopina oligoneura* | Parikkala | 61.566 | 29.561 | 21-Jun-2021 | MaT |
| ZMUO.074733 | BOLD:ACD2463 | *Borophaga agilis* | Parikkala | 61.566 | 29.561 | 21-Jun-2021 | MaT |
| ZMUO.074740 | BOLD:ACF4704 | *Metopina oligoneura* | Parikkala | 61.566 | 29.561 | 21-Jun-2021 | MaT |
| ZMUO.074743 | BOLD:ACF4704 | *Metopina oligoneura* | Parikkala | 61.566 | 29.561 | 21-Jun-2021 | MaT |
| ZMUO.074746 | BOLD:ACF4704 | *Metopina oligoneura* | Parikkala | 61.566 | 29.561 | 21-Jun-2021 | MaT |
| ZMUO.074751 | BOLD:ADM5152 | *Triphleba antricola* | Parikkala | 61.566 | 29.561 | 21-Jun-2021 | MaT |
| ZMUO.074754 | BOLD:ACF4704 | *Metopina oligoneura* | Parikkala | 61.566 | 29.561 | 21-Jun-2021 | MaT |
| ZMUO.074755 | BOLD:ACF4704 | *Metopina oligoneura* | Parikkala | 61.566 | 29.561 | 21-Jun-2021 | MaT |
| ZMUO.074757 | BOLD:ACF4704 | *Metopina oligoneura* | Parikkala | 61.566 | 29.561 | 21-Jun-2021 | MaT |
| ZMUO.074766 | BOLD:ACF4704 | *Metopina oligoneura* | Parikkala | 61.566 | 29.561 | 21-Jun-2021 | MaT |

| | | | | | | | |
|---|---|---|---|---|---|---|---|
| ZMUO.074768 | BOLD:ACF4704 | *Metopina oligoneura* | Parikkala | 61.566 | 29.561 | 21-Jun-2021 | MaT |
| ZMUO.074769 | BOLD:ACF4704 | *Metopina oligoneura* | Parikkala | 61.566 | 29.561 | 21-Jun-2021 | MaT |
| ZMUO.074773 | BOLD:ACF4704 | *Metopina oligoneura* | Parikkala | 61.566 | 29.561 | 21-Jun-2021 | MaT |
| ZMUO.074777 | BOLD:ACF4704 | *Metopina oligoneura* | Parikkala | 61.566 | 29.561 | 21-Jun-2021 | MaT |
| ZMUO.074778 | BOLD:ACF4704 | *Metopina oligoneura* | Parikkala | 61.566 | 29.561 | 21-Jun-2021 | MaT |
| ZMUO.074781 | BOLD:ACF4704 | *Metopina oligoneura* | Parikkala | 61.566 | 29.561 | 21-Jun-2021 | MaT |
| ZMUO.074782 | BOLD:ACB3871 | *Aenigmatias lubbockii* | Parikkala | 61.566 | 29.561 | 21-Jun-2021 | MaT |
| ZMUO.074785 | BOLD:ACF4704 | *Metopina oligoneura* | Parikkala | 61.566 | 29.561 | 21-Jun-2021 | MaT |
| ZMUO.074789 | BOLD:AAG3236 | *Diplonevra nitidula* | Parikkala | 61.566 | 29.561 | 21-Jun-2021 | MaT |
| ZMUO.074796 | BOLD:ACB3701 | *Diplonevra concinna* | Parikkala | 61.566 | 29.561 | 21-Jun-2021 | MaT |
| ZMUO.074797 | BOLD:ACP3737 | *Phora edentata* | Parikkala | 61.566 | 29.561 | 21-Jun-2021 | MaT |
| ZMUO.074808 | BOLD:ACF4704 | *Metopina oligoneura* | Parikkala | 61.566 | 29.561 | 19-Jul-2021 | MaT |
| ZMUO.074819 | BOLD:ACF4704 | *Metopina oligoneura* | Parikkala | 61.566 | 29.561 | 19-Jul-2021 | MaT |
| ZMUO.074831 | BOLD:ACF4704 | *Metopina oligoneura* | Parikkala | 61.566 | 29.561 | 19-Jul-2021 | MaT |
| ZMUO.074834 | BOLD:ADM8144 | *Phora obscura* | Parikkala | 61.566 | 29.561 | 19-Jul-2021 | MaT |
| ZMUO.074848 | BOLD:ACP3737 | *Phora edentata* | Parikkala | 61.566 | 29.561 | 19-Jul-2021 | MaT |
| ZMUO.074862 | BOLD:ACD5037 | *Triphleba nudipalpis* | Parikkala | 61.566 | 29.561 | 19-Jul-2021 | MaT |
| ZMUO.074864 | BOLD:ADM5152 | *Triphleba antricola* | Parikkala | 61.566 | 29.561 | 19-Jul-2021 | MaT |
| ZMUO.074867 | BOLD:ACD2261 | *Phora artifrons* | Parikkala | 61.566 | 29.561 | 19-Jul-2021 | MaT |
| ZMUO.074879 | BOLD:ADM5152 | *Triphleba antricola* | Parikkala | 61.566 | 29.561 | 19-Jul-2021 | MaT |
| ZMUO.074881 | BOLD:ABA7001 | *Diplonevra glabra* | Parikkala | 61.566 | 29.561 | 19-Jul-2021 | MaT |
| ZMUO.074885 | BOLD:ACB3394 | *Phora bullata* | Parikkala | 61.566 | 29.561 | 19-Jul-2021 | MaT |
| ZMUO.074896 | BOLD:ACF4704 | *Metopina oligoneura* | Parikkala | 61.566 | 29.561 | 19-Jul-2021 | MaT |
| ZMUO.074898 | BOLD:ACB3412 | *Triphleba nudipalpis* | Parikkala | 61.566 | 29.561 | 19-Jul-2021 | MaT |
| ZMUO.074902 | BOLD:ACF4704 | *Metopina oligoneura* | Parikkala | 61.566 | 29.561 | 19-Jul-2021 | MaT |
| ZMUO.074904 | BOLD:ACF4704 | *Metopina oligoneura* | Parikkala | 61.566 | 29.561 | 19-Jul-2021 | MaT |
| ZMUO.074907 | BOLD:ACF6067 | *Phora dubia* | Parikkala | 61.566 | 29.561 | 19-Jul-2021 | MaT |
| ZMUO.074912 | BOLD:ACF4704 | *Metopina oligoneura* | Parikkala | 61.566 | 29.561 | 19-Jul-2021 | MaT |
| ZMUO.074914 | BOLD:ACF6067 | *Phora dubia* | Parikkala | 61.566 | 29.561 | 19-Jul-2021 | MaT |
| ZMUO.074918 | BOLD:ADM5152 | *Triphleba antricola* | Parikkala | 61.566 | 29.561 | 19-Jul-2021 | MaT |
| ZMUO.074929 | BOLD:ACF4704 | *Metopina oligoneura* | Parikkala | 61.566 | 29.561 | 19-Jul-2021 | MaT |
| ZMUO.074949 | BOLD:ADM5152 | *Triphleba antricola* | Parikkala | 61.566 | 29.561 | 19-Jul-2021 | MaT |
| ZMUO.074974 | BOLD:ACF4704 | *Metopina oligoneura* | Parikkala | 61.566 | 29.561 | 19-Jul-2021 | MaT |
| ZMUO.074982 | BOLD:ACF4704 | *Metopina oligoneura* | Parikkala | 61.566 | 29.561 | 19-Jul-2021 | MaT |
| ZMUO.074987 | BOLD:ACF6067 | *Phora dubia* | Parikkala | 61.566 | 29.561 | 19-Jul-2021 | MaT |
| ZMUO.074989 | BOLD:ABA7002 | *Phora tincta* | Parikkala | 61.566 | 29.561 | 19-Jul-2021 | MaT |
| ZMUO.074990 | BOLD:ACF6067 | *Phora dubia* | Parikkala | 61.566 | 29.561 | 19-Jul-2021 | MaT |
| ZMUO.074992 | BOLD:ACB3412 | *Triphleba nudipalpis* | Parikkala | 61.566 | 29.561 | 19-Jul-2021 | MaT |
| ZMUO.074996 | BOLD:ACB3394 | *Phora bullata* | Parikkala | 61.566 | 29.561 | 19-Jul-2021 | MaT |
| ZMUO.075025 | BOLD:ACB3412 | *Triphleba nudipalpis* | Parikkala | 61.566 | 29.561 | 19-Jul-2021 | MaT |
| ZMUO.075026 | BOLD:ACF6067 | *Phora dubia* | Parikkala | 61.566 | 29.561 | 19-Jul-2021 | MaT |
| ZMUO.075047 | BOLD:ACF4704 | *Metopina oligoneura* | Parikkala | 61.566 | 29.561 | 19-Jul-2021 | MaT |
| ZMUO.075053 | BOLD:ADW4894 | *Pseudacteon lundbecki* | Parikkala | 61.566 | 29.561 | 19-Jul-2021 | MaT |
| ZMUO.075059 | BOLD:ACB3412 | *Triphleba nudipalpis* | Parikkala | 61.566 | 29.561 | 19-Jul-2021 | MaT |
| ZMUO.075067 | BOLD:ACF6067 | *Phora dubia* | Parikkala | 61.566 | 29.561 | 19-Jul-2021 | MaT |
| ZMUO.075072 | BOLD:ACE2645 | *Conicera floricola* | Parikkala | 61.566 | 29.561 | 19-Jul-2021 | MaT |
| ZMUO.075074 | BOLD:ACP3737 | *Phora edentata* | Parikkala | 61.566 | 29.561 | 19-Jul-2021 | MaT |
| ZMUO.075081 | BOLD:ADM8636 | *Phora hamata* | Parikkala | 61.566 | 29.561 | 19-Jul-2021 | MaT |

| | | | | | | | |
|---|---|---|---|---|---|---|---|
| ZMUO.075099 | BOLD:ACF6067 | *Phora dubia* | Parikkala | 61.566 | 29.561 | 19-Jul-2021 | MaT |
| ZMUO.075115 | BOLD:ABA7002 | *Phora tincta* | Parikkala | 61.566 | 29.561 | 19-Jul-2021 | MaT |
| ZMUO.075123 | BOLD:ACE2645 | *Conicera floricola* | Parikkala | 61.566 | 29.561 | 19-Jul-2021 | MaT |
| ZMUO.075145 | BOLD:ADM5152 | *Triphleba antricola* | Parikkala | 61.566 | 29.561 | 19-Jul-2021 | MaT |
| ZMUO.075153 | BOLD:ACF4704 | *Metopina oligoneura* | Parikkala | 61.566 | 29.561 | 19-Jul-2021 | MaT |
| ZMUO.075155 | BOLD:ACF4704 | *Metopina oligoneura* | Parikkala | 61.566 | 29.561 | 19-Jul-2021 | MaT |
| ZMUO.075156 | BOLD:ACB3412 | *Triphleba nudipalpis* | Parikkala | 61.566 | 29.561 | 19-Jul-2021 | MaT |
| ZMUO.075177 | BOLD:ACF6067 | *Phora dubia* | Parikkala | 61.566 | 29.561 | 19-Jul-2021 | MaT |
| ZMUO.075183 | BOLD:ABA7002 | *Phora tincta* | Parikkala | 61.566 | 29.561 | 19-Jul-2021 | MaT |
| ZMUO.075190 | BOLD:ADM5152 | *Triphleba antricola* | Parikkala | 61.566 | 29.561 | 19-Jul-2021 | MaT |
| ZMUO.075191 | BOLD:ADM5152 | *Triphleba antricola* | Parikkala | 61.566 | 29.561 | 19-Jul-2021 | MaT |
| ZMUO.075208 | BOLD:ABA7001 | *Diplonevra glabra* | Parikkala | 61.566 | 29.561 | 19-Jul-2021 | MaT |
| ZMUO.075226 | BOLD:ACF6067 | *Phora dubia* | Parikkala | 61.566 | 29.561 | 19-Jul-2021 | MaT |
| ZMUO.075238 | BOLD:ADM5152 | *Triphleba antricola* | Parikkala | 61.566 | 29.561 | 19-Jul-2021 | MaT |
| ZMUO.075272 | BOLD:ACD2261 | *Phora artifrons* | Saarijarvi | 62.842 | 25.483 | 22-Jun-2021 | MaT |
| ZMUO.075554 | BOLD:ACR0978 | *Pseudacteon formicarum* | Kitee | 62.045 | 30.261 | 24-Jul-2016 | MaT |
| ZMUO.075604 | BOLD:ACR0791 | *Triphleba lugubris* | Kitee | 62.045 | 30.261 | 30-Jul-2016 | MaT |
| ZMUO.075873 | BOLD:ACB3653 | *Diplonevra freyi* | Kitee | 62.045 | 30.261 | 19-Jul-2016 | MaT |
| ZMUO.075970 | BOLD:ACB3394 | *Phora bullata* | Kitee | 62.045 | 30.261 | 19-Jul-2016 | MaT |
| ZMUO.076023 | BOLD:ACO9836 | *Triphleba distinguenda* | Kitee | 62.045 | 30.261 | 19-Jul-2016 | MaT |
| ZMUO.076032 | BOLD:ACO7758 | *Pseudacteon brevicauda* | Kitee | 62.045 | 30.261 | 19-Jul-2016 | MaT |
| ZMUO.076046 | BOLD:ACD2464 | *Borophaga carinifrons* | Kitee | 62.045 | 30.261 | 19-Jul-2016 | MaT |
| ZMUO.076088 | BOLD:ACI8210 | *Phora convergens* | Kitee | 62.045 | 30.261 | 22-Aug-2016 | MaT |
| ZMUO.076246 | BOLD:ABA7001 | *Diplonevra glabra* | Kitee | 62.045 | 30.261 | 25-May-2016 | MaT |
| ZMUO.076258 | BOLD:ACF6067 | *Phora dubia* | Kitee | 62.045 | 30.261 | 25-May-2016 | MaT |
| ZMUO.076286 | BOLD:ADM5152 | *Triphleba antricola* | Lappeenranta | 61.061 | 28.745 | 21-Jun-2021 | MaT |
| ZMUO.076289 | BOLD:ABA7001 | *Diplonevra glabra* | Lappeenranta | 61.061 | 28.745 | 21-Jun-2021 | MaT |
| ZMUO.076292 | BOLD:ABA7001 | *Diplonevra glabra* | Lappeenranta | 61.061 | 28.745 | 21-Jun-2021 | MaT |
| ZMUO.076304 | BOLD:ACF6067 | *Phora dubia* | Lappeenranta | 61.061 | 28.745 | 21-Jun-2021 | MaT |
| ZMUO.076313 | BOLD:ACF6067 | *Phora dubia* | Lappeenranta | 61.061 | 28.745 | 21-Jun-2021 | MaT |
| ZMUO.076328 | BOLD:ABA7001 | *Diplonevra glabra* | Lappeenranta | 61.061 | 28.745 | 21-Jun-2021 | MaT |
| ZMUO.076329 | BOLD:ADM5152 | *Triphleba antricola* | Lappeenranta | 61.061 | 28.745 | 21-Jun-2021 | MaT |
| ZMUO.076335 | BOLD:AAG3236 | *Diplonevra nitidula* | Lappeenranta | 61.061 | 28.745 | 21-Jun-2021 | MaT |
| ZMUO.076336 | BOLD:ACD2261 | *Phora artifrons* | Lappeenranta | 61.061 | 28.745 | 21-Jun-2021 | MaT |
| ZMUO.076337 | BOLD:ABA7002 | *Phora tincta* | Lappeenranta | 61.061 | 28.745 | 21-Jun-2021 | MaT |
| ZMUO.076338 | BOLD:ACD2261 | *Phora artifrons* | Lappeenranta | 61.061 | 28.745 | 21-Jun-2021 | MaT |
| ZMUO.076348 | BOLD:ACD2261 | *Phora artifrons* | Lappeenranta | 61.061 | 28.745 | 21-Jun-2021 | MaT |
| ZMUO.076354 | BOLD:ACD2261 | *Phora artifrons* | Lappeenranta | 61.061 | 28.745 | 21-Jun-2021 | MaT |
| ZMUO.076355 | BOLD:ADM5152 | *Triphleba antricola* | Lappeenranta | 61.061 | 28.745 | 21-Jun-2021 | MaT |
| ZMUO.076356 | BOLD:ABA7002 | *Phora tincta* | Lappeenranta | 61.061 | 28.745 | 21-Jun-2021 | MaT |
| ZMUO.076358 | BOLD:ACD2261 | *Phora artifrons* | Lappeenranta | 61.061 | 28.745 | 21-Jun-2021 | MaT |
| ZMUO.076368 | BOLD:ACD2261 | *Phora artifrons* | Lappeenranta | 61.061 | 28.745 | 21-Jun-2021 | MaT |
| ZMUO.076372 | BOLD:ADM5152 | *Triphleba antricola* | Lappeenranta | 61.061 | 28.745 | 21-Jun-2021 | MaT |
| ZMUO.076373 | BOLD:ADM5152 | *Triphleba antricola* | Lappeenranta | 61.061 | 28.745 | 21-Jun-2021 | MaT |
| ZMUO.076391 | BOLD:ADM5152 | *Triphleba antricola* | Lappeenranta | 61.061 | 28.745 | 21-Jun-2021 | MaT |
| ZMUO.076417 | BOLD:ACD2261 | *Phora artifrons* | Lappeenranta | 61.061 | 28.745 | 21-Jun-2021 | MaT |
| ZMUO.076419 | BOLD:ABA7001 | *Diplonevra glabra* | Lappeenranta | 61.061 | 28.745 | 21-Jun-2021 | MaT |

| | | | | | | | |
|---|---|---|---|---|---|---|---|
| ZMUO.076422 | BOLD:ABA7001 | *Diplonevra glabra* | Lappeenranta | 61.061 | 28.745 | 21-Jun-2021 | MaT |
| ZMUO.076454 | BOLD:AAG3236 | *Diplonevra nitidula* | Lappeenranta | 61.061 | 28.745 | 21-Jun-2021 | MaT |
| ZMUO.076463 | BOLD:ADM5152 | *Triphleba antricola* | Lappeenranta | 61.061 | 28.745 | 21-Jun-2021 | MaT |
| ZMUO.076470 | BOLD:ACD2261 | *Phora artifrons* | Lappeenranta | 61.061 | 28.745 | 21-Jun-2021 | MaT |
| ZMUO.076474 | BOLD:ADM5152 | *Triphleba antricola* | Lappeenranta | 61.061 | 28.745 | 21-Jun-2021 | MaT |
| ZMUO.076488 | BOLD:ADM5152 | *Triphleba antricola* | Lappeenranta | 61.061 | 28.745 | 21-Jun-2021 | MaT |
| ZMUO.076499 | BOLD:ABA7002 | *Phora tincta* | Lappeenranta | 61.061 | 28.745 | 19-Jul-2021 | MaT |
| ZMUO.076513 | BOLD:ABA7001 | *Diplonevra glabra* | Lappeenranta | 61.061 | 28.745 | 19-Jul-2021 | MaT |
| ZMUO.076516 | BOLD:AAP9864 | *Triphleba subcompleta* | Lappeenranta | 61.061 | 28.745 | 19-Jul-2021 | MaT |
| ZMUO.076525 | BOLD:AAG3236 | *Diplonevra nitidula* | Lappeenranta | 61.061 | 28.745 | 19-Jul-2021 | MaT |
| ZMUO.076530 | BOLD:ADM5152 | *Triphleba antricola* | Lappeenranta | 61.061 | 28.745 | 19-Jul-2021 | MaT |
| ZMUO.076541 | BOLD:AAG3236 | *Diplonevra nitidula* | Lappeenranta | 61.061 | 28.745 | 19-Jul-2021 | MaT |
| ZMUO.076544 | BOLD:AAG3236 | *Diplonevra nitidula* | Lappeenranta | 61.061 | 28.745 | 19-Jul-2021 | MaT |
| ZMUO.076552 | BOLD:ADM5152 | *Triphleba antricola* | Lappeenranta | 61.061 | 28.745 | 19-Jul-2021 | MaT |
| ZMUO.076554 | BOLD:ACZ8391 | *Diplonevra pilosella* | Lappeenranta | 61.061 | 28.745 | 19-Jul-2021 | MaT |
| ZMUO.076561 | BOLD:ACD2463 | *Borophaga agilis* | Lappeenranta | 61.061 | 28.745 | 19-Jul-2021 | MaT |
| ZMUO.076564 | BOLD:AAP9864 | *Triphleba subcompleta* | Lappeenranta | 61.061 | 28.745 | 19-Jul-2021 | MaT |
| ZMUO.076568 | BOLD:AAG3236 | *Diplonevra nitidula* | Lappeenranta | 61.061 | 28.745 | 19-Jul-2021 | MaT |
| ZMUO.076569 | BOLD:ADM5152 | *Triphleba antricola* | Lappeenranta | 61.061 | 28.745 | 19-Jul-2021 | MaT |
| ZMUO.076586 | BOLD:AAG3236 | *Diplonevra nitidula* | Lappeenranta | 61.061 | 28.745 | 19-Jul-2021 | MaT |
| ZMUO.076597 | BOLD:ADM5152 | *Triphleba antricola* | Lappeenranta | 61.061 | 28.745 | 19-Jul-2021 | MaT |
| ZMUO.076599 | BOLD:ABA7002 | *Phora tincta* | Lappeenranta | 61.061 | 28.745 | 19-Jul-2021 | MaT |
| ZMUO.076609 | BOLD:ABA7002 | *Phora tincta* | Lappeenranta | 61.061 | 28.745 | 19-Jul-2021 | MaT |
| ZMUO.076617 | BOLD:AAG3236 | *Diplonevra nitidula* | Lappeenranta | 61.061 | 28.745 | 19-Jul-2021 | MaT |
| ZMUO.076631 | BOLD:AAG3236 | *Diplonevra nitidula* | Lappeenranta | 61.061 | 28.745 | 19-Jul-2021 | MaT |
| ZMUO.076633 | BOLD:ADM5152 | *Triphleba antricola* | Lappeenranta | 61.061 | 28.745 | 19-Jul-2021 | MaT |
| ZMUO.076635 | BOLD:ACD2444 | *Hypocera mordellaria* | Lappeenranta | 61.061 | 28.745 | 19-Jul-2021 | MaT |
| ZMUO.076641 | BOLD:ACD2463 | *Borophaga agilis* | Lappeenranta | 61.061 | 28.745 | 19-Jul-2021 | MaT |
| ZMUO.076666 | BOLD:ADM5152 | *Triphleba antricola* | Lappeenranta | 61.061 | 28.745 | 19-Jul-2021 | MaT |
| ZMUO.076682 | BOLD:ADM5152 | *Triphleba antricola* | Lappeenranta | 61.061 | 28.745 | 19-Jul-2021 | MaT |
| ZMUO.076685 | BOLD:ACI8210 | *Phora convergens* | Lappeenranta | 61.061 | 28.745 | 19-Jul-2021 | MaT |
| ZMUO.076703 | BOLD:ACN6664 | *Phora pubipes* | Lappeenranta | 61.061 | 28.745 | 19-Jul-2021 | MaT |
| ZMUO.076711 | BOLD:ACQ9655 | *Anevrina unispinosa* | Lappeenranta | 61.061 | 28.745 | 19-Jul-2021 | MaT |
| ZMUO.076724 | BOLD:ACN6664 | *Phora pubipes* | Lappeenranta | 61.061 | 28.745 | 19-Jul-2021 | MaT |
| ZMUO.076729 | BOLD:ADM5152 | *Triphleba antricola* | Lappeenranta | 61.061 | 28.745 | 19-Jul-2021 | MaT |
| ZMUO.076730 | BOLD:ACN6664 | *Phora pubipes* | Lappeenranta | 61.061 | 28.745 | 19-Jul-2021 | MaT |
| ZMUO.076745 | BOLD:AAP9864 | *Triphleba subcompleta* | Lappeenranta | 61.061 | 28.745 | 19-Jul-2021 | MaT |
| ZMUO.076753 | BOLD:ADM5152 | *Triphleba antricola* | Lappeenranta | 61.061 | 28.745 | 19-Jul-2021 | MaT |
| ZMUO.076756 | BOLD:ADM5152 | *Triphleba antricola* | Lappeenranta | 61.061 | 28.745 | 19-Jul-2021 | MaT |
| ZMUO.076760 | BOLD:ADM5152 | *Triphleba antricola* | Lappeenranta | 61.061 | 28.745 | 19-Jul-2021 | MaT |
| ZMUO.076763 | BOLD:ACD2463 | *Borophaga agilis* | Lappeenranta | 61.061 | 28.745 | 19-Jul-2021 | MaT |
| ZMUO.076813 | BOLD:AFE7151 | *Anevrina thoracica* | Lappeenranta | 61.063 | 28.735 | 21-Jun-2021 | MaT |
| ZMUO.076837 | BOLD:ACD2261 | *Phora artifrons* | Lappeenranta | 61.063 | 28.735 | 21-Jun-2021 | MaT |
| ZMUO.076860 | BOLD:ACQ9655 | *Anevrina unispinosa* | Lappeenranta | 61.063 | 28.735 | 21-Jun-2021 | MaT |
| ZMUO.076892 | BOLD:ACF6067 | *Phora dubia* | Lappeenranta | 61.063 | 28.735 | 21-Jun-2021 | MaT |
| ZMUO.076910 | BOLD:ABA7002 | *Phora tincta* | Lappeenranta | 61.063 | 28.735 | 21-Jun-2021 | MaT |
| ZMUO.076914 | BOLD:ACF6067 | *Phora dubia* | Lappeenranta | 61.063 | 28.735 | 21-Jun-2021 | MaT |
| ZMUO.076958 | BOLD:ACE2645 | *Conicera floricola* | Lappeenranta | 61.063 | 28.735 | 19-Jul-2021 | MaT |

| | | | | | | | |
|---|---|---|---|---|---|---|---|
| ZMUO.076963 | BOLD:ACR0978 | *Pseudacteon formicarum* | Lappeenranta | 61.063 | 28.735 | 19-Jul-2021 | MaT |
| ZMUO.076996 | BOLD:AFE7151 | *Anevrina thoracica* | Lappeenranta | 61.063 | 28.735 | 19-Jul-2021 | MaT |
| ZMUO.077036 | BOLD:AAP9864 | *Triphleba subcompleta* | Lappeenranta | 61.063 | 28.735 | 19-Jul-2021 | MaT |
| ZMUO.077099 | BOLD:ACB3701 | *Diplonevra concinna* | Lappeenranta | 61.063 | 28.735 | 19-Jul-2021 | MaT |

S-Table 2. Non-*Megaselia* species identified using BOLD Identification Engine with >98% match, with the number of specimens from which *COI* barcode was recovered. Retrieved November 2023.

| Species | Specimen count |
|---|---|
| *Aenigmatias lubbockii* | 1 |
| *Anevrina thoracica* | 7 |
| *Borophaga agilis* | 21 |
| *Borophaga carinifrons* | 50 |
| *Borophaga femorata* | 1 |
| *Borophaga subsultans* | 3 |
| *Chaetopleurophora erythronota* | 1 |
| *Conicera dauci* | 1 |
| *Conicera floricola* | 9 |
| *Conicera schnittmanni* | 1 |
| *Conicera similis* | 2 |
| *Diplonevra concinna* | 12 |
| *Diplonevra florescens* | 1 |
| *Diplonevra freyi* | 1 |
| *Diplonevra glabra* | 52 |
| *Diplonevra nitidula* | 66 |
| *Gymnophora arcuata* | 1 |
| *Hypocera mordellaria* | 4 |
| *Metopina oligoneura* | 76 |
| *Phalacrotophora fasciata* | 6 |
| *Phora artifrons* | 27 |
| *Phora atra* | 1 |
| *Phora convergens* | 37 |
| *Phora dubia* | 10 |
| *Phora edentata* | 4 |
| *Phora hamata* | 1 |
| *Phora holosericea* | 5 |
| *Phora obscura* | 2 |
| *Phora occidentata* | 1 |
| *Phora pubipes* | 131 |
| *Phora stictica* | 1 |
| *Phora tincta* | 108 |
| *Pseudacteon brevicauda* | 2 |
| *Pseudacteon fennicus* | 1 |
| *Spiniphora excisa* | 2 |
| *Triphleba aequalis* | 4 |
| *Triphleba bicornuta* | 4 |
| *Triphleba citreiformis* | 1 |
| *Triphleba distinguenda* | 30 |
| *Triphleba lugubris* | 93 |

| | |
|---|---|
| *Triphleba nudipalpis* | 9 |
| *Triphleba papillata* | 1 |
| *Triphleba subcompleta* | 7 |

S-Table 3. Known species representation among samples.

| The Finnish Checklist | Malaise trap material |
|---|---|
| *Abaristophora arctophila* | Not found |
| *Abaristophora kolaensis* | Not found |
| *Aenigmatias franzi* | Not found |
| *Aenigmatias lubbockii* | Present |
| *Aenigmatias picipes* | Not found |
| *Anevrina thoracica* | Present |
| *Anevrina unispinosa* | Present |
| *Anevrina urbana* | Not found |
| *Borophaga agilis* | Present |
| *Borophaga bennetti* | Not found |
| *Borophaga carinifrons* | Present |
| *Borophaga femorata* | Present |
| *Borophaga incrassata* | Not found |
| *Borophaga irregularis* | Not found |
| *Borophaga subsultans* | Present |
| *Chaetopleurophora bohemanni* | Not found |
| *Chaetopleurophora erythronota* | Present |
| *Chaetopleurophora spinosissima* | Not found |
| *Conicera dauci* | Present |
| *Conicera floricola* | Present |
| *Conicera schnittmanni* | Present |
| *Conicera similis* | Present |
| *Conicera tarsalis* | Not found |
| *Conicera tibialis* | Not found |
| *Diplonevra concinna* | Present |
| *Diplonevra florescens* | Present |
| *Diplonevra freyi* | Present |
| *Diplonevra funebris* | Not found |
| *Diplonevra glabra* | Present |
| *Diplonevra nitidula* | Present |
| *Diplonevra oldenbergi* | Not found |
| *Diplonevra pilosella* | Present |
| *Dohrniphora cornuta* | Not found |
| *Gymnophora arcuata* | Present |
| *Gymnophora bifida* | Not found |
| *Gymnophora distincta* | Not found |
| *Gymnophora forresteri* | Not found |
| *Gymnophora healeyae* | Not found |

| | |
|---|---|
| *Gymnophora nigripennis* | Not found |
| *Gymnophora perpropinqua* | Not found |
| *Gymnophora quartomollis* | Not found |
| *Gymnophora winqvisti* | Not found |
| *Gymnoptera longicostalis* | Not found |
| *Hypocera mordellaria* | Present |
| *Menozziola obscuripes* | Not found |
| *Menozziola schmitzi* | Not found |
| *Metopina galeata* | Not found |
| *Metopina oligoneura* | Present |
| *Metopina pileata* | Not found |
| *Microselia forsiusi* | Not found |
| *Phalacrotophora berolinensis* | Not found |
| *Phalacrotophora beuki* | Not found |
| *Phalacrotophora fasciata* | Present |
| *Phora artifrons* | Present |
| *Phora atra* | Present |
| *Phora bullata* | Present |
| *Phora convallium* | Not found |
| *Phora convergens* | Present |
| *Phora dubia* | Present |
| *Phora edentata* | Present |
| *Phora hamata* | Present |
| *Phora holosericea* | Present |
| *Phora hyperborea* | Not found |
| *Phora indivisa* | Not found |
| *Phora obscura* | Present |
| *Phora occidentata* | Present |
| *Phora penicillata* | Not found |
| *Phora praepandens* | Not found |
| *Phora pubipes* | Present |
| *Phora stictica* | Present |
| *Phora tincta* | Present |
| *Plectanocnema nudipes* | Present |
| *Pseudacteon fennicus* | Present |
| *Pseudacteon formicarum* | Present |
| *Pseudacteon lundbecki* | Present |
| *Spiniphora bergenstammi* | Not found |
| *Spiniphora dorsalis* | Not found |
| *Spiniphora excisa* | Present |
| *Spiniphora jugorum* | Not found |
| *Spiniphora maculata* | Not found |
| *Triphleba admirabilis* | Not found |
| *Triphleba aequalis* | Present |

| | |
|---|---|
| *Triphleba antricola* | Present |
| *Triphleba autumnalis* | Not found |
| *Triphleba bicornuta* | Present |
| *Triphleba citreiformis* | Present |
| *Triphleba cumsetae* | Not found |
| *Triphleba dentata* | Not found |
| *Triphleba distinguenda* | Present |
| *Triphleba excisa* | Not found |
| *Triphleba gilvipes* | Not found |
| *Triphleba gracilis* | Not found |
| *Triphleba hyalinata* | Not found |
| *Triphleba inaequalis* | Not found |
| *Triphleba intermedia* | Not found |
| *Triphleba lugubris* | Present |
| *Triphleba luteifemorata* | Not found |
| *Triphleba minuta* | Not found |
| *Triphleba nudipalpis* | Present |
| *Triphleba opaca* | Not found |
| *Triphleba pachyneurella* | Not found |
| *Triphleba palposa* | Not found |
| *Triphleba papillata* | Present |
| *Triphleba renidens* | Not found |
| *Triphleba salmelai* | Not found |
| *Triphleba subcompleta* | Present |
| *Triphleba sunnmorkensis* | Not found |
| *Triphleba trinervis* | Not found |
| *Trucidophora ewardurskae* | Not found |
| *Veruanus oldenbergi* | Not found |
